# PHARMACOLOGICAL PROTEIN INACTIVATION BY TARGETING FOLDING INTERMEDIATES

**DOI:** 10.1101/2020.03.31.018069

**Authors:** Giovanni Spagnolli, Tania Massignan, Andrea Astolfi, Silvia Biggi, Paolo Brunelli, Michela Libergoli, Alan Ianeselli, Simone Orioli, Alberto Boldrini, Luca Terruzzi, Giulia Maietta, Marta Rigoli, Nuria Lopez Lorenzo, Leticia C. Fernandez, Laura Tosatto, Luise Linsenmeier, Beatrice Vignoli, Gianluca Petris, Dino Gasparotto, Maria Pennuto, Graziano Guella, Marco Canossa, Hermann Clemens Altmeppen, Graziano Lolli, Stefano Biressi, Manuel Martin Pastor, Jesús R. Requena, Ines Mancini, Maria Letizia Barreca, Pietro Faccioli, Emiliano Biasini

**Affiliations:** Department of Cellular, Computational and Integrative Biology (CIBIO), University of Trento, 38123 Povo, Trento TN, ITALY; Dulbecco Telethon Institute, University of Trento, 38123 Povo, Trento TN, ITALY; Department of Pharmaceutical Sciences, University of Perugia, 06123 Perugia PG, ITALY; Department of Physics, University of Trento, Povo, Trento TN, ITALY; INFN-TIFPA, University of Trento, Povo, Trento TN, ITALY; Sibylla Biotech SRL, 37121 Verona VR, ITALY; RIAIDT, University of Santiago de Compostela-IDIS, 15782 SPAIN; CIMUS Biomedical Research Institute & Department of Medical Sciences, University of Santiago de Compostela-IDIS, 15782 SPAIN; Institute of Biophysics, National Council of Research, 38123 Povo, Trento TN, Italy; Institute of Neuropathology, University Medical Center Hamburg-Eppendorf, 20246 Hamburg, GERMANY; Department of Biomedical Sciences (DBS), University of Padova, 35131 Padova PD, ITALY

## Abstract

Recent computational advancements in the simulation of biochemical processes allow investigating the mechanisms involved in protein regulation with realistic physics-based models, at an atomistic level of resolution. Using these techniques to study the negative regulation of the androgen receptor (AR), we discovered a key functional role played by non-native metastable states appearing along the folding pathway of this protein. This unexpected observation inspired us to design a completely novel drug discovery approach, named Pharmacological Protein Inactivation by Folding Intermediate Targeting (PPI-FIT), based on the rationale of negatively regulating protein expression by targeting folding intermediates. Here, PPI-FIT was tested for the first time on the cellular prion protein (PrP), a cell surface glycoprotein playing a key role in fatal and transmissible neurodegenerative pathologies known as prion diseases. We predicted the all-atom structure of an intermediate appearing along the folding pathway of PrP, and identified four different small molecule ligands for this conformer, all capable of selectively lowering the expression of the protein by promoting its degradation. Our data support the notion that the level of target proteins could be modulated by acting on their folding pathways, implying a previously unappreciated role for folding intermediates in the biological regulation of protein expression.

## INTRODUCTION

Protein expression and function in eukaryotic cells are tightly harmonized processes modulated by the combination of different layers of regulation. These may occur at the level of gene transcription, processing, stability, and translation of the mRNA as well as by assembly, post-translational modifications, sorting, recycling, and degradation of the corresponding polypeptide ^1^. In addition, the expression of a small subset of proteins is known to be regulated by the presence of specific ligands, which are required along the folding pathway to reach the native, functional state ^2-7^. For example, the folding of the kinase-inducible domain of the cAMP-response-element-binding protein is known to proceed only upon interaction with the CREB-binding protein ^8^. Other typical examples of ligand-dependent folding mechanism include nuclear receptors, such as estrogen and androgen receptors ^9^. The integration between these pathways and the protein quality control machinery, deputed to avoid the production and accumulation of aberrantly folded proteins, is known as proteostasis ^1,10^. Mechanisms by which proteostasis is ensured include chaperone-assisted protein folding as well as re-routing and degradation of misfolded protein conformers. These processes play a fundamental role during development, aging, tolerance to environmental stresses, and minimize alterations of proteome homeostasis induced by pathogens ^11^. Consistently, a large percentage of human pathologies are linked to alterations of proteostasis, including a broad spectrum of age-related brain disorders, known as neurodegenerative diseases, characterized by the accumulation of aberrantly folded proteins in the central nervous system ^12,13^. These range from highly frequent disorders, such as Alzheimer’s and Parkinson’s diseases, to rather rare conditions such as amyotrophic lateral sclerosis and prion diseases. The latter, also known as transmissible spongiform encephalopathies, are caused by the conformational conversion of a cell surface glycoprotein, named the cellular prion protein (PrP), into an aggregated, pathogenic form (called PrP^Sc^) capable of propagating like an infectious agent (prion) by templating the structural conversion of its physiological counterpart ^14-16^. Growing evidence indicates that prions exemplify a mechanism of protein-based propagation of information exerted by many other amyloids in pathological and physiological contexts ^17,18^. Unfortunately, the absence of detailed information regarding misfolding processes, as well as ineffective clearance by the proteostasis machinery, have so far hampered the development of therapies for the vast majority of neurodegenerative disorders.

Fundamental insights into the regulation of protein expression, folding and misfolding could theoretically be provided by the full reconstruction of the conformational transitions underlying folding pathways. However, experimental approaches aimed at addressing the dynamics of protein folding are seriously affected by the tradeoff between temporal and spatial resolutions of available biophysical techniques ^19^. Computer-based technologies, such as Molecular Dynamics (MD) simulations, could, in principle, overcome these limitations and help to predict the evolution in time of protein conformations. In practice, however, classical MD simulations are not applicable to study many fundamental molecular processes, such as the cascade of events underlying the folding of a typical, biologically-relevant polypeptide bigger than 80 amino acids and with folding times longer than a few milliseconds ^20^. The reason is that the timescales which can be simulated by MD, even on the most powerful supercomputers, are still orders of magnitude shorter than those at which major structural transitions take place ^21^. To overcome some of these limitations, we have developed algorithms that reduce computational costs by exploiting available experimental information about native protein structures ^22,23^. Here, we employed recent technological advancements in the field of computational biochemistry to shed light on a mechanism of negative regulation of the androgen receptor (AR) and identified a role for intermediate folding species appearing before the protein acquires the native state. These observations inspired us to design a completely novel drug discovery paradigm (named PPI-FIT) based on the rationale of negatively regulating the expression of a given protein by targeting folding intermediates. The PPI-FIT approach was tested for the first time on PrP, as compelling genetic and experimental evidence indicates that lowering the expression of this protein could produce therapeutic benefits in prion diseases without causing major side effects ^24-27^.

## RESULTS

### Elucidation of an AR folding regulation mechanism at atomistic resolution

The AR is responsible for hormone-dependent transcription of genes involved in prostate cell proliferation and differentiation ^28^. Once produced in the Leydig cells of testes, testosterone circulates in the bloodstream bound to serum sex hormone-binding globulin. The free form of this hormone can cross the cell membrane and may be converted into 5-dihydrotestosterone (DHT) into the cytoplasm. The inactive form of the AR resides into the cytoplasm bound to heat shock proteins, which are then displaced upon binding of the AR to DHT, leading the receptor to reach its functional conformation, translocate into the nucleus and activate transcription of target genes ^29^. Ligand binding to the AR stabilizes its conformation by inducing an intramolecular structural re-organization of helix 12 that caps the ligand-binding pocket (Supp. Figure 1). Additionally, the AR could be regulated by a number of post-translational modifications, including phosphorylation, ubiquitination, SUMOylation, acetylation, and methylation ^30^. In one of these cases, it has been shown that serine 791 (S791) within the ligand-binding domain of the AR could be phosphorylated by the Protein Kinase B ^31^. This modification inhibits ligand interaction and promotes polyubiquitination and degradation of the AR by the proteasome ^32^. Interestingly, in natively folded, ligand-bound AR, the S791 residue is buried inside the protein core, suggesting that regulation of this phosphorylation site likely happens before the AR binds to its ligand (Supp. Table 1 and Supp. Figure 2). Based on this conclusion, we hypothesized that phosphorylation of S791 might occur on a metastable state along the folding pathway of the AR. In order to test this possibility, we employed the bias functional (BF) method to reconstruct the folding pathway of the AR (see Methods). The BF algorithm is a recently developed computational scheme that allows the simulation of protein folding transitions at all-atom resolution using physics-based models ^22,33^. This method has been validated against both plain MD simulations and biophysical experiments ^34-36^. The application of the BF approach requires to provide as input the structure attained at the end of the transition. In this case, we used as a reference structure the ligand-binding domain of the AR (PDB 3B68, residue 671 to 917) without including the atoms of the ligand. First, we generated 3 unfolded conformations by thermal unfolding. Next, 20 folding trajectories for each conformation were generated using the ratchet and pawl molecular dynamics (rMD) algorithm. The resulting ensemble (60 trajectories) was used to generate a probability distribution plot (Figure 1A). The BF scheme was then applied to rank all generated trajectories in order to select the most realistic prediction for each set (i.e., the so-called least biased trajectories). This procedure enables us to keep at a minimum the systematic errors introduced by the rMD scheme. Finally, conformations laying within observed energy wells were sampled from each least biased trajectory. These analyses yielded 3 sequential, relevant non-native metastable states, each represented by 3 conformations: (i) a partially unfolded state (U), characterized by the presence of α-helical structures largely lacking native contacts; (ii) an intermediate folding state (I1), partially resembling the native conformation; (iii) a second intermediate folding state (I2), characterized by a higher degree of native-like packing (Figure 1B). We observed that U is the only state in which S791 is exposed in all the conformations, whereas this residue is completely buried in both I1 and I2 states (Figure 1C-D). These data indicate that phosphorylation of the S791 most likely occurs when the protein is in a U-like state. Interestingly, the ligand-binding pocket in state I1 is already properly structured and shows favorable druggability descriptors, and this site is not hindered by helix-12. Therefore, the I1 conformations represent viable ligand-binding species of the AR (Supp. Figure 3 and Supp. Table 2). These data suggest that at least two regulatory mechanisms of the AR activity target intermediate species appearing along the folding pathway of the protein.

**Figure 1.**
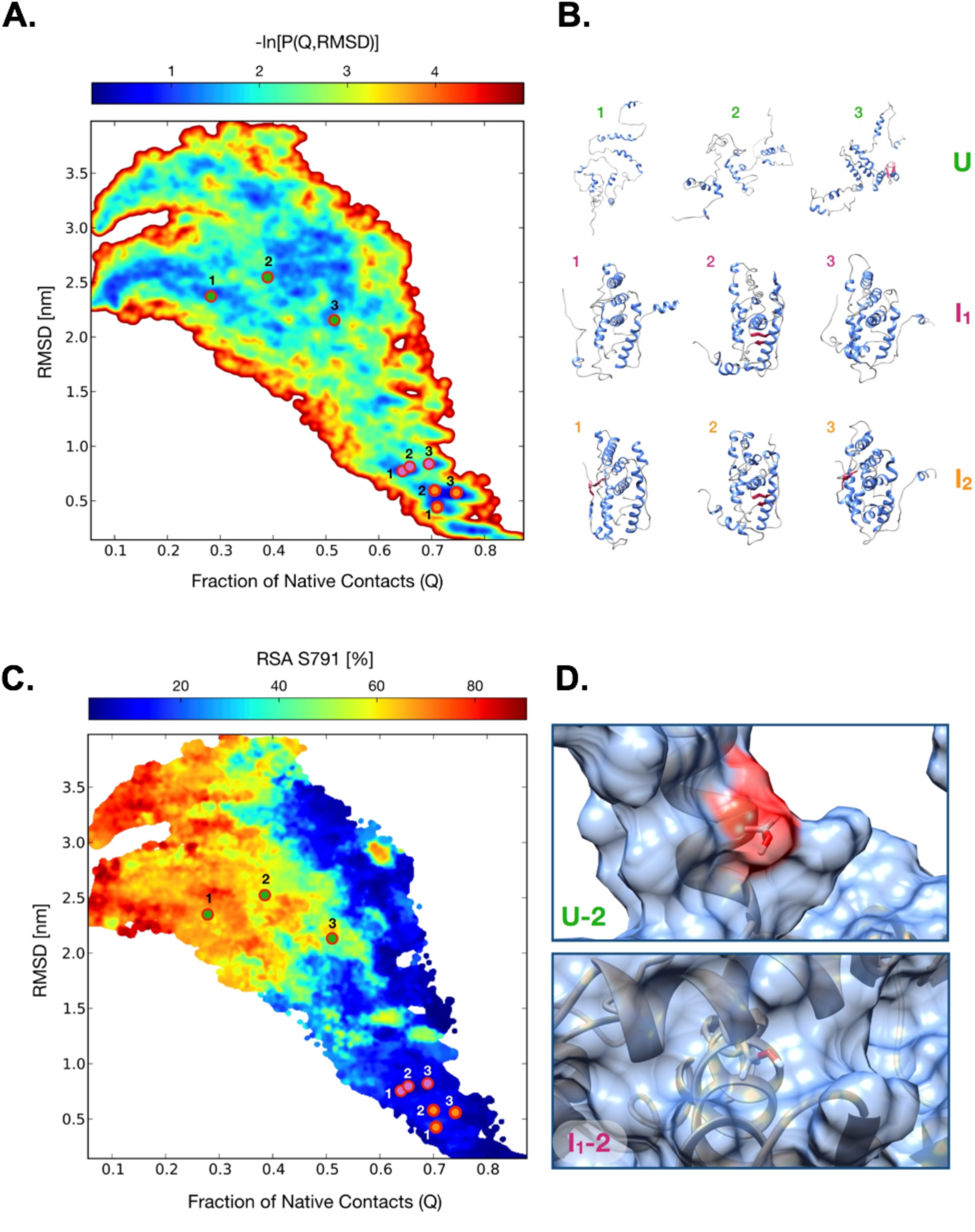
All-atom reconstruction of the AR ligand-binding domain folding pathway. **A**. Lower-bound approximation of the free energy landscape of AR folding obtained from 60 rMD trajectories, plotted as a negative logarithm of the probability distribution expressed as a function of the collective variables Q (fraction of native contacts) and RMSD. **B**. Representative AR structures of non-native (RMSD > 4 Å) conformations U, I_1_ and I_2_, each sampled from 3 least biased trajectories belonging to relevant energy wells (G ≤ 2.5 k_B_T). **C**. Relative surface area of residue S791 along the AR folding pathway expressed as a function of Q and RMSD. **D**. Images show the solvent-exposed surface of S791 (depicted in red) in the U-2 (upper panel) or I_1_-2 (lower panel) conformations, taken as representative examples of solvent accessibility or non-accessibility of the residue in the U and I_1_ states, respectively.

### Identification of a PrP folding intermediate

The observation that the regulation of the AR may occur at the level of its folding pathway inspired us to test the possibility of pharmacologically modulating the expression of a given protein by targeting folding intermediates. We named this new experimental paradigm as Pharmacological Protein Inactivation by Folding Intermediate Targeting (PPI-FIT). The approach is based on designing compounds against the most kinetically- and thermodynamically-relevant folding intermediate of a given protein, in order to stabilize such intermediate and inhibit its transition to the native form. In a cellular environment, a stabilized folding intermediate could be recognized by the protein folding quality control machinery as an improperly folded polypeptide and sent to degradation ^37-39^. Therefore, the underlying rationale of PPI-FIT is to design compounds capable of post-translationally decreasing the expression of a target protein. We sought to test this new drug discovery paradigm on PrP, based on four aspects: (i) several previous reports indicate that at least one intermediate is generated along the folding pathway of PrP ^40-42^; (ii) the genetic suppression of PrP in animal models has been shown to produce little side effects only later in life ^43^. This conclusion is also supported by loss-of-function PRNP alleles observed in healthy human subjects ^26^; (iii) strong experimental evidence demonstrates that lowering the expression of PrP would confer therapeutic benefits in prion diseases, and possibly other neurodegenerative disorders linked to the toxicity-transducing activity of the protein ^24,25,27,44-48^; (iv) however, multiple previous failures to target the native conformation of PrP pharmacologically suggest that this protein could be an undruggable factor ^49,50^. In light of these considerations, we decided to identify the folding intermediate of PrP by reconstructing the entire sequence of events underlying the folding pathway of this protein at an atomistic level of resolution. The protein consists of a flexible, N-terminal segment (residues 24-120, human sequence), and a structured, C-terminal domain (residues 121-231) comprising three α-helices and two short β-strands flanking helix 1. The N-terminal segment is intrinsically disordered, a feature precluding its use for in-silico approaches ^51^. Thus, as a reference structure, we used the globular domain of human PrP (residues 125-228, PDB 1QLX; Supp. Figure 4). We generated 9 unfolded PrP conformations by thermal unfolding and used rMD to produce 20 folding trajectories for each conformation (making a total of 180 trajectories plotted for their bidimensional probability distribution in Figure 2A). The BF scheme was then employed to define the most statistically-significant folding pathway for each set of trajectories, leading to 9 least biased folding pathways (an example is shown in the Supp. Movie). Conformations residing within the observed energy wells in the bidimensional distribution were sampled from each least biased trajectory (Figure 2B). The analyses revealed the existence of an on-pathway PrP folding intermediate structurally close to the native state, and explored by all least biased pathways. Subsequent clustering analysis of the ensemble of conformers populating the intermediate state enabled the definition of an all-atom structure for the PrP folding intermediate (Supp. Figure 5). As compared to the PrP native conformation, this structure is characterized by a displaced helix-1 missing its docking contacts with helix 3, leaving several hydrophobic residues natively buried inside the protein core exposed to the solvent (e.g., Y157, M206, V209; Figure 2C).

**Figure 2.**
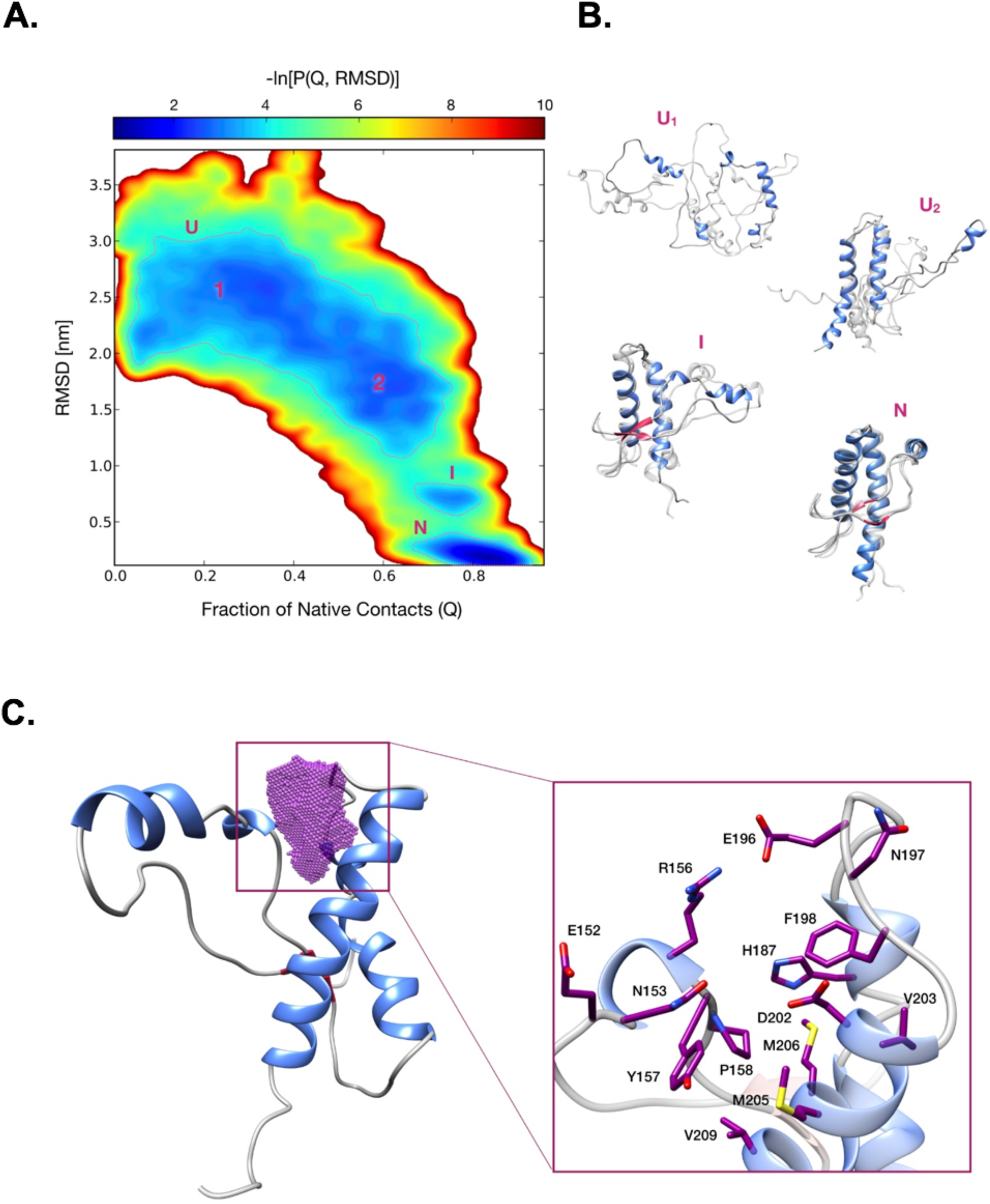
All-atom reconstruction of the PrP folding pathway. **A**. Lower-bound approximation of the free energy landscape of the PrP folding obtained from 180 rMD trajectories, plotted as a negative logarithm of the probability distribution expressed as a function of the collective variables Q (fraction of native contacts) and RMSD. The dashed lines delimit the metastable regions of interest (G ≤ 3.7 k_B_T). **B**. Representative PrP structures of unfolded (U_1_ and U_2_), folding intermediate (I) and native conformations (N). Each conformation represents the cluster center of one of the corresponding populated regions (structures depicted in transparency are sampled from the same region, showing conformations variability): the first two (corresponding to the unfolded states U_1_ and U_2_) are characterized by RMSD > 1.8 nm and a Q < 0.5 (U_1_), and 1.2 < RMSD < 1.8 nm and a 0.5 < Q < 0.75 (U_2_); a third one (corresponding to the intermediate folding state I) with 0.55 < RMSD < 0.90 nm and a 0.65 < Q < 0.85; a fourth one (corresponding to the native state N) with RMSD < 0.40 nm and Q > 0.80. **C**. Ribbon diagram of the PrP intermediate highlighting the ligand-binding pocket (purple dots) as identified by SiteMap and DogSiteScorer tools. The purple volume maps the unique druggable pocket identified in the PrP folding intermediate I. The box shows specific residues defining the site.

### In silico identification of high-affinity ligands for the PrP folding intermediate

We employed in silico modeling and virtual screening techniques to identify small ligands for the PrP folding intermediate. First, we searched for druggable binding sites unique in the structure of the latter and not present in the native PrP conformation. This step was carried out by means of SiteMap (Schrödinger Release 2017-4: SiteMap, Schrödinger, LLC, New York, NY, 2017) ^52^ and DoGSiteScorer ^53^ software. The results highlighted the presence of a possible ligand-binding region, placed between the beginning of helix α1 and the loop that connects the α2 and α3 helixes (Supp. Table 3). MD simulations refined the structure of a solvent-accessible, druggable pocket (Supp Table 3) defined by 14, non-continuous residues (152, 153, 156, 157, 158, 187, 196, 197, 198, 202, 203, 205, 206, 209) (Figure 2C). This site was the target of a virtual screening campaign performed by using the BioSolveIT software (www.biosolveit.de) and employing the Asinex Gold & Platinum collection of small molecules (∼3.2 × 10^5^ commercially available compounds, www.asinex.com). The obtained poses were filtered according to the predicted estimated affinity, physicochemical, and ADMET (absorption, distribution, metabolism, excretion, and toxicity) properties as well as 3D visualization/assessment of the ligand-binding interactions. In addition, molecules potentially acting as pan-assay interference compounds were filtered, and chemoinformatics methods for similarity and clustering analysis were applied to support a diversity-based selection. Finally, 30 compounds were selected as promising virtual hits and purchased for biological validation (Supp. Table 4 and Supp. Figure 6).

### In vitro validation of in silico hits

PrP biogenesis follows a trafficking pathway typical of glycosylphosphatidylinositol (GPI)-anchored polypeptides ^54-56^. The protein is synthesized directly in the lumen of the endoplasmic reticulum (ER), where it folds and receives post-translational processing of the primary structure (removal of the signal peptide and anchoring of the GPI moiety) as well as the addition of two high-mannose N-linked glycans (at Asn-181 and Asn-197) (see ^57^ for a review). Based on the role of lysosomes in the quality control of PrP expression, a compound binding to a PrP folding intermediate may produce a long-living, immature conformer that could be recognized by the ER quality control (ERQC) machinery, leading to its degradation by the ER-associated lysosome-dependent autophagy ^58^. Following this principle, the 30 putative ligands were directly tested in cells for their ability to lower the expression of PrP at a post-translational level. Each compound was administered for 48 h at different concentrations (1-30 µM) to HEK293 cells stably expressing mouse PrP. The expression and post-translational alterations of PrP were detected by western blotting. Compounds showing an ability to lower the amount or alter the post-translational processing of PrP (≥ 30%) at any of the concentrations were tested against a control protein, the Neuronal Growth Regulator 1 (NEGR-1), a member of the immunoglobulin LON family. This GPI-anchored molecule follows the same biosynthesis pathway as PrP, thus representing an ideal control to evaluate compound specificity ^59^. We validated 4 compounds (coded SM930, SM940, SM950, and SM875), all of them capable of decreasing the levels of PrP in HEK293 cells without lowering the expression of the control protein NEGR-1 (Figure 3 A-D). Among these, compound SM875 was selected for further analyses, based on its potency (a docking poses of the compound bound to the PrP folding intermediate is shown in Figure 3E).

**Figure 3.**
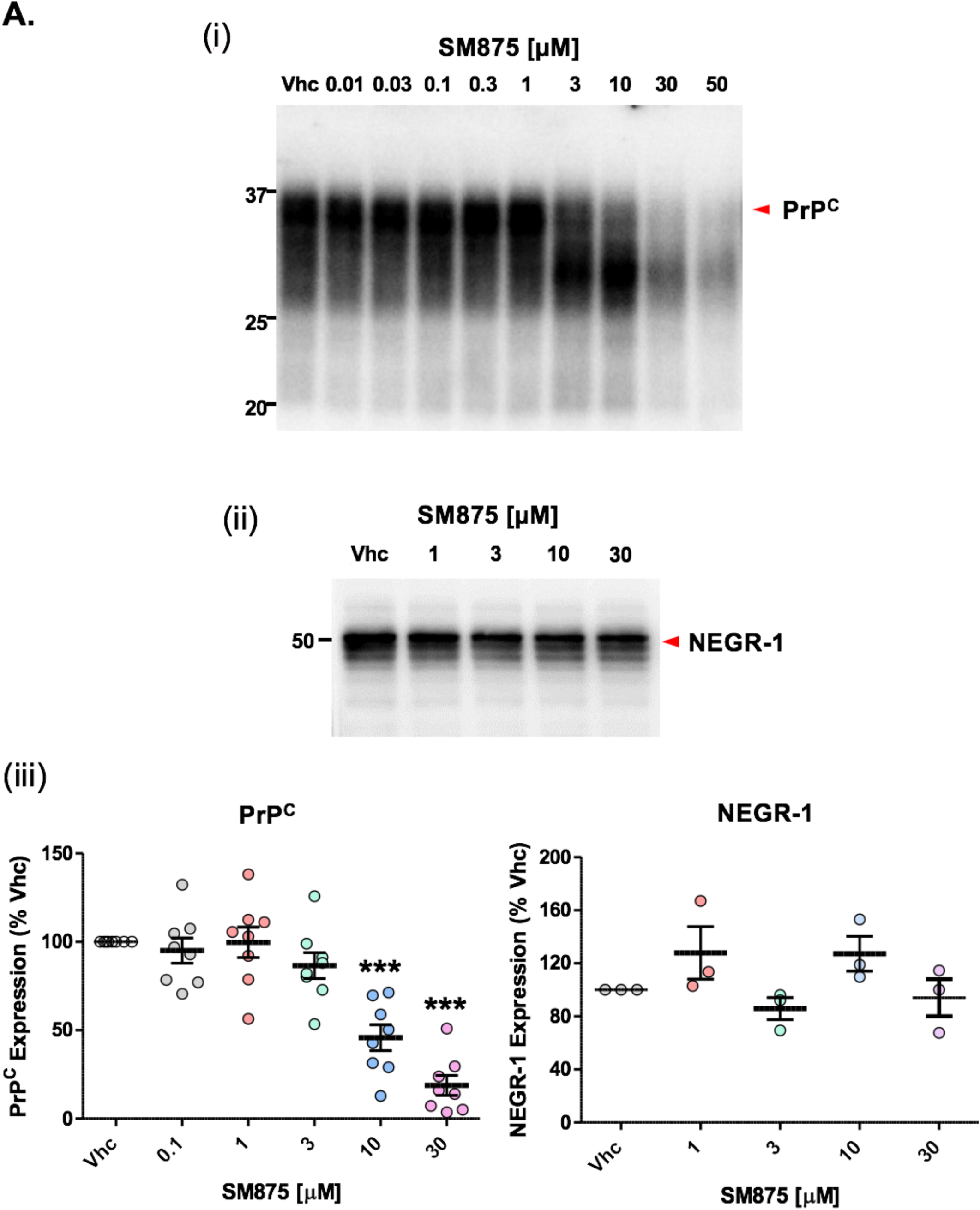
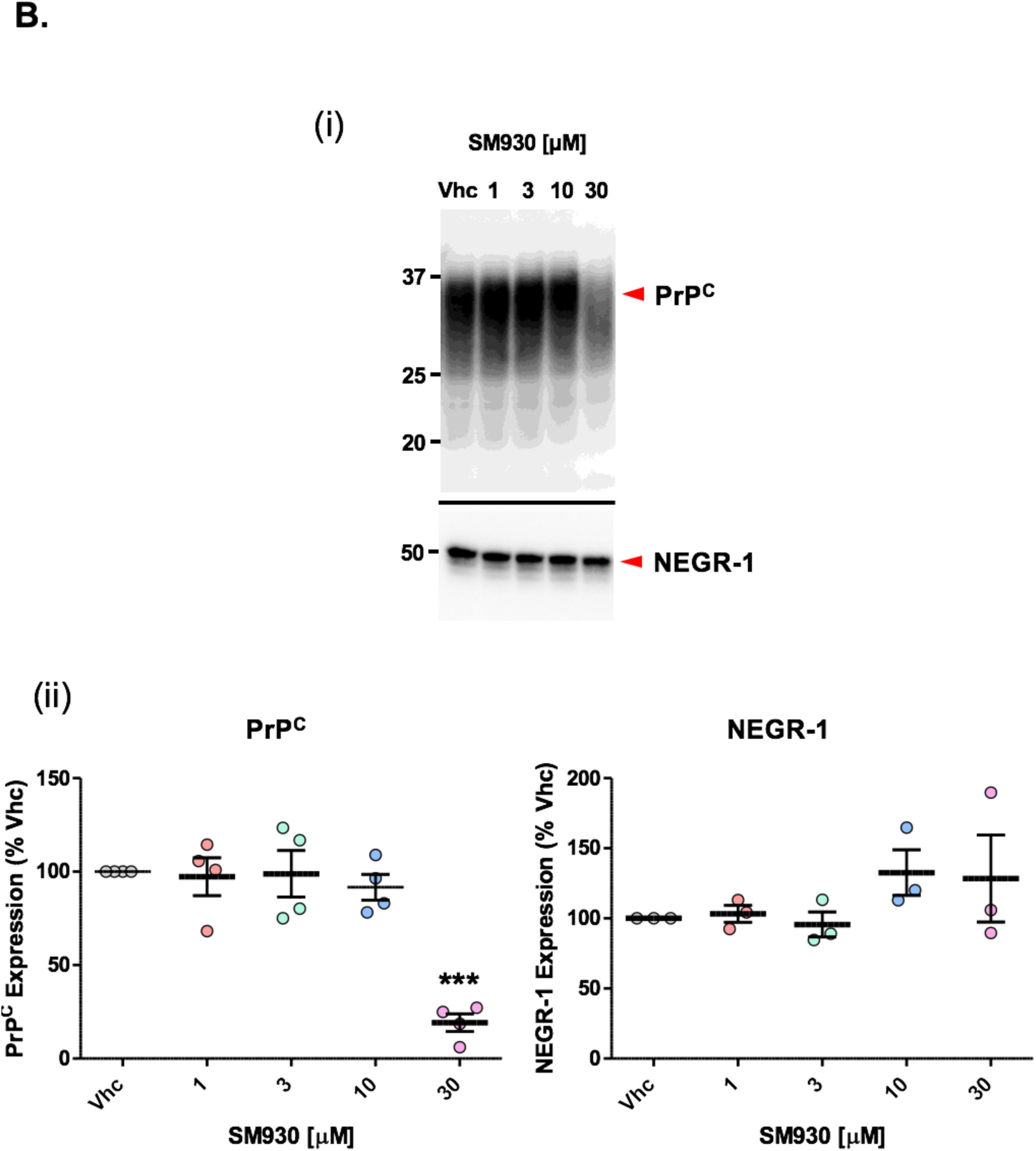
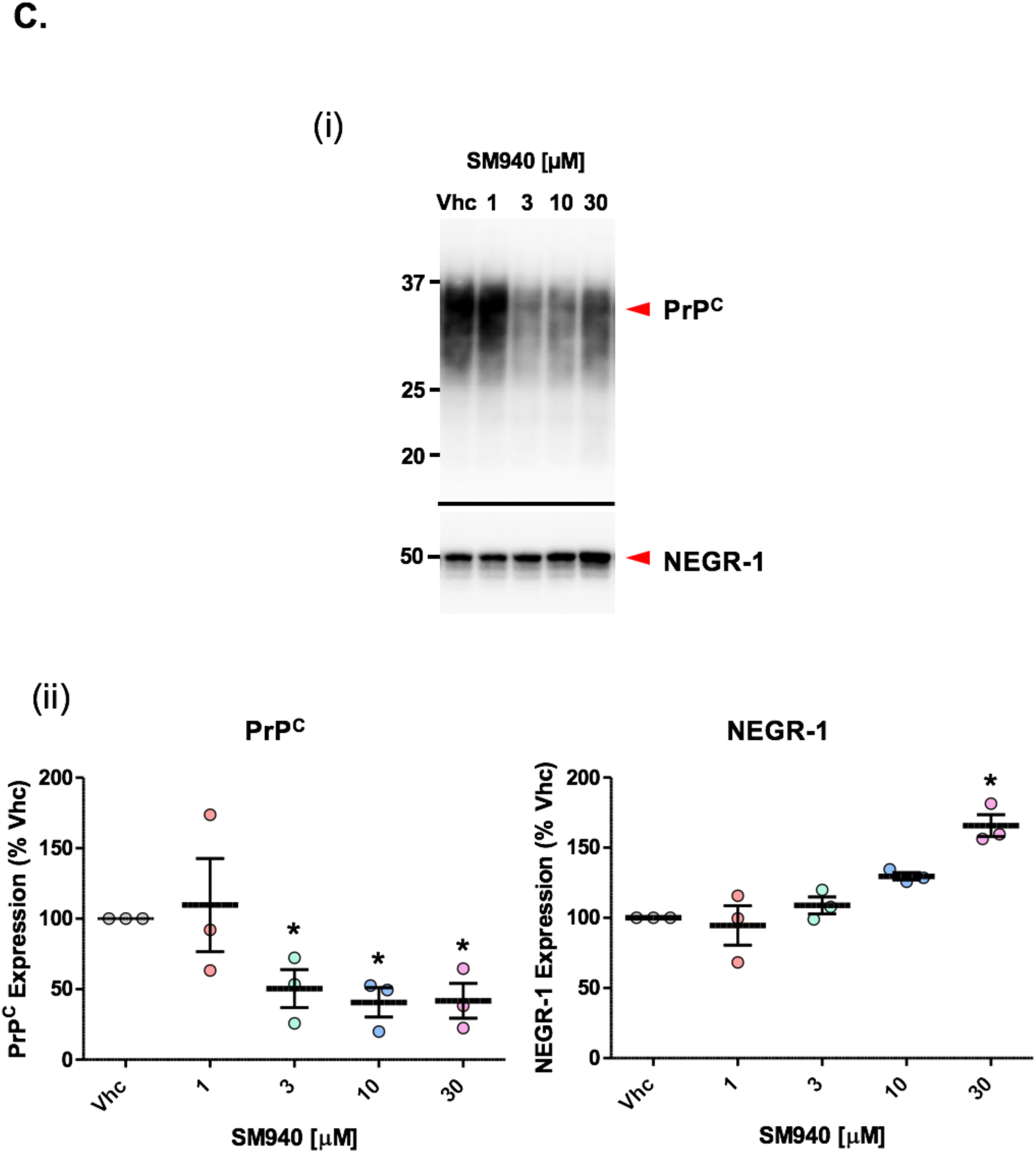
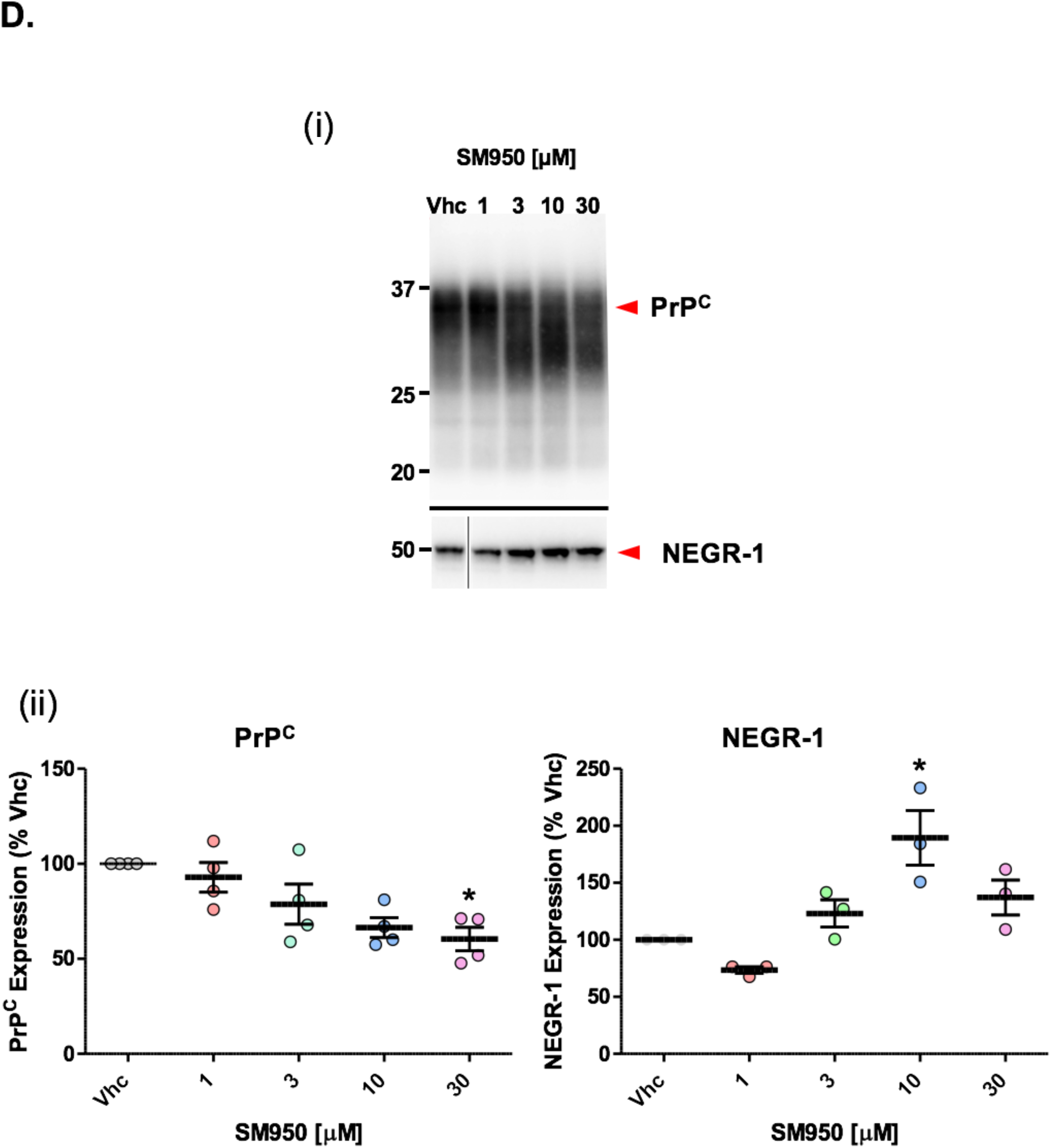
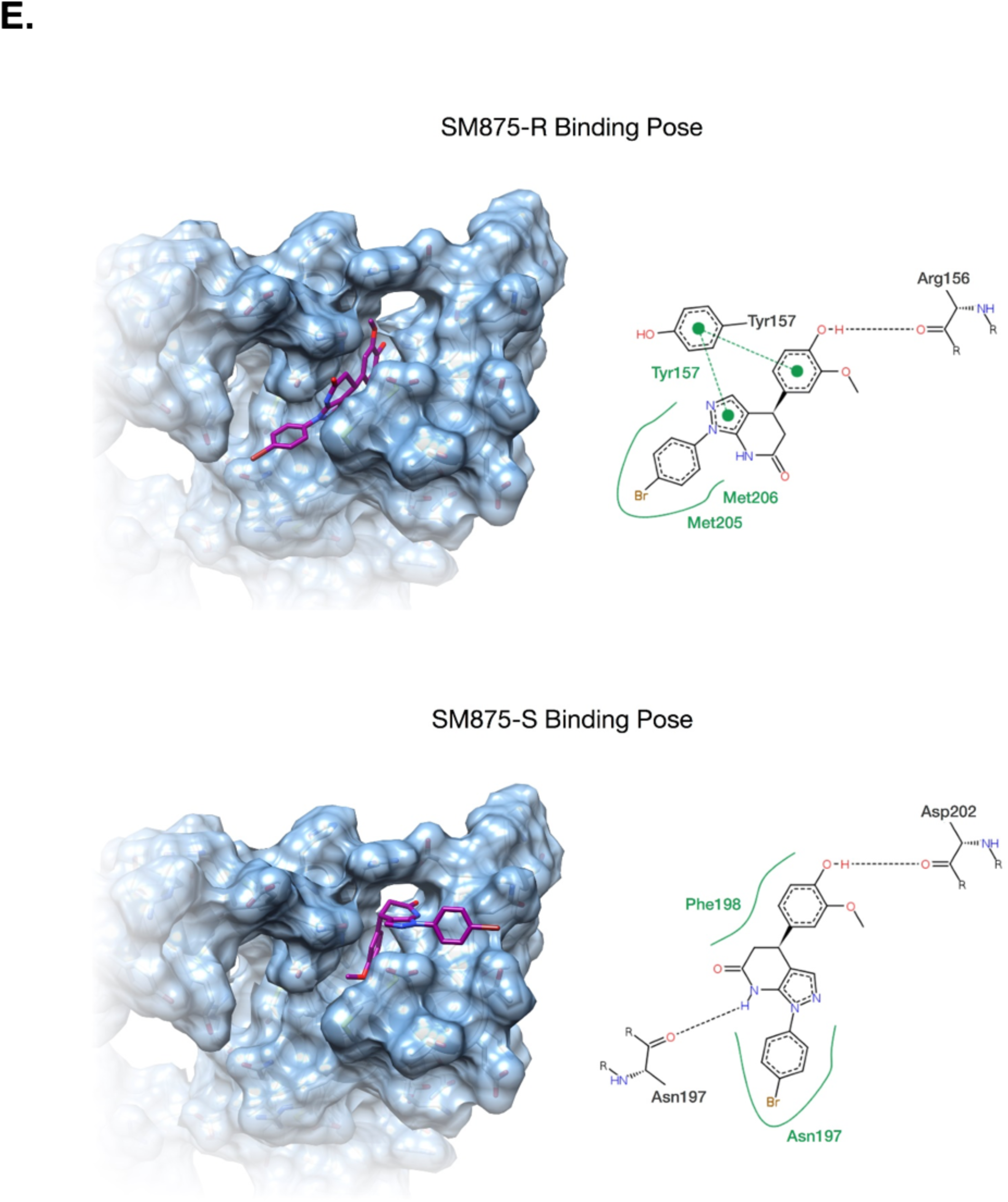
In vitro validation of selected hits. **A**. HEK293 cells (ATCC, CRL-1573) stably transfected with mouse PrP or NEGR-1 were exposed to different concentrations of SM875 (indicated) or vehicle (DMSO, volume equivalent) for 48 h, lysed and analyzed by western blotting. Signals were detected by using a specific anti-PrP (D18) or anti-NEGR-1 primary antibodies, relevant HRP-coupled secondary antibodies, and revealed using a ChemiDoc Touch Imaging System (Bio-Rad, CA, USA). Red arrows indicate the expected sizes of mature, fully glycosylated forms of PrP and NEGR-1. **(i)** At the lowest active concentrations (3-10 µM), compound SM875 shows a dose-dependent effect on the expression of PrP, accompanied by a progressive shift of the signal from the expected ∼35 kDa of the full-length, di-glycosylated form of the protein, to lower molecular weight bands, likely corresponding to cleaved PrP forms. At the highest active concentrations (30-50 µM), the whole PrP signal progressively disappears, possibly consistent with increased degradation of all the forms of the protein; **(ii)** the compound shows no effect on NEGR-1 levels when tested at concentrations similar to those at which it lowers PrP expression; **(iii)** the graphs show the densitometric quantification of the levels of full-length PrP (left) or NEGR-1 (right) from different independent replicates (n = 8 for PrP; n = 3 for NEGR-1). Each signal was normalized on the corresponding total protein lane (detected by UV, and allowed the enhanced tryptophan fluorescence technology of stain-free gels) and expressed as the percentage of the level in vehicle (Vhc)-treated controls (*** p < 0.005). **B**. HEK293 cells stably transfected with mouse PrP or NEGR-1 were exposed to different concentrations of SM930 (indicated) or vehicle (DMSO, volume equivalent) for 48 h, lysed and analyzed by western blotting; **(i)** Red arrows indicate the expected sizes of mature, fully glycosylated forms of PrP and NEGR-1. Compound SM930 decreases the whole expression of PrP, but not NEGR-1, at the highest dose tested (30 µM); **(ii)** the graphs show the densitometric quantification of the levels of full-length PrP (left) or NEGR-1 (right) from different independent replicates (n = 4 for PrP; n = 3 for NEGR-1). Each signal was normalized on the corresponding total protein lane (detected by UV) and expressed as the percentage of the level in vehicle (Vhc)-treated controls (*** p < 0.005). **C**. HEK293 cells stably transfected with mouse PrP or NEGR-1 were exposed to different concentrations of SM940 (indicated) or vehicle (DMSO, volume equivalent) for 48 h, lysed and analyzed by western blotting. **(i)** Red arrows indicate the expected sizes of mature, fully glycosylated forms of PrP and NEGR-1. Compound SM940 shows a lowering effect on the whole expression of PrP, starting from the concentration of 3 µM. The molecule also induces an unexpected, significant increase of NEGR-1 expression at 30 µM. The reason for this effect has not been further investigated; **(ii)** the graphs show the densitometric quantification of the levels of full-length PrP (left) or NEGR-1 (right) from different independent replicates (n = 3 n = 3 for PrP and NEGR-1). Each signal was normalized on the corresponding total protein lane (detected by UV) and expressed as the percentage of the level in vehicle (Vhc)-treated controls (* p < 0.05). **D**. HEK293 cells stably transfected with mouse PrP (i) or NEGR-1 (ii) were exposed to different concentrations of compound SM950 (indicated) or vehicle (DMSO, volume equivalent) for 48 h, lysed and analyzed by western blotting. **(i)** Red arrows indicate the expected sizes of mature, fully glycosylated forms of PrP and NEGR-1. The compound shows a dose-dependent effect on the expression of PrP, starting from the concentration of 3 µM, accompanied by a progressive shift of the signal from the expected ∼35 kDa of the full-length, di-glycosylated form of the protein, to lower molecular weight bands, likely corresponding to cleaved PrP forms. SM950 shows an unexpected, significant increase of NEGR-1 expression at the single concentration of 10 µM, an observation not investigated further in this manuscript; **(ii)** the graphs show the densitometric quantification of the levels of full-length PrP (left) or NEGR-1 (right) from different independent replicates (n = 4 for PrP; n = 3 for NEGR-1). Each signal was normalized on the corresponding total protein lane (detected by UV) and expressed as the percentage of the level in vehicle (Vhc)-treated controls (* p < 0.05). **E**. The picture illustrates the predicted ligand binding pose of the R (upper panel) and S (lower panel) SM875 enantiomer into the PrP intermediate druggable pocket. For each enantiomer, 10 poses were generated by FlexX and then rescored using the HYDE algorithm. Only the best rescored pose for each enantiomer was retained and analyzed (SM875-R:405 nM < K_iHYDE_ < 40275 nM; SM875-S: 2252nM < K_iHYDE_ < 223796 nM). On the left side of each panel, the SM875 enantiomer (purple stick) is bound to the druggable pocket (light blue surface), whereas the simplified 2D representation of the intermolecular ligand-protein interactions is reported on the right side. The interaction pattern is composed of hydrogen bonds, visualized as black dashed lines, π interactions, shown as green dashed lines with dots denoting the participating π systems, and hydrophobic contacts, which are represented by the residue labels and segments along the contacting hydrophobic ligand surface.

### Synthesis and biological characterization of SM875

In order to obtain large amounts of the compound for additional assays, we designed and performed a novel synthesis scheme for SM875 (Supp. Figure 7). The correct structure of the molecule was then verified by nuclear magnetic resonance (NMR), mass spectrometry, and infrared (IR) spectroscopy (Supp. Figure 8A-D and Supp. Table 5). The dose-dependent, PrP-lowering activity of the newly synthesized SM875 molecule was validated using a larger concentration range (0.01-50 µM) in stably transfected HEK293 cells, allowing the calculation of the inhibitory concentration at 50% in the low micromolar range (IC50 = 7.87 ± 1.17 µM; Figure 3A). Next, to rule out possible artifacts due to the exogenous transfection of PrP, we tested the effect of SM875 in a human breast cancer cell line (ZR-75) endogenously expressing human PrP. These cells were identified among the NCI collection of human tumor cell lines as high PrP expression (Supp. Figure 9). Compound SM875 decreased PrP levels in a dose-dependent fashion also in these cells, in the exact same concentration range (1-30 µM) observed in transfected HEK293 cells (Figure 4A). Importantly, SM875 did not alter the expression of Thy-1, another GPI-anchored protein endogenously expressed in ZR-75 and used as a control (Figure 4A) ^60^. Consistent with these data, SM875 reduced the expression of PrP in two additional untransfected cell lines, L929 mouse fibroblasts, and mouse N2a neuroblastoma cells (Figure 4B-C). Interestingly, in N2a cells SM875 did not show a dose-response effect, as PrP decrease was observed only at lower concentrations (1-10 µM) while the protein showed normal expression levels at the highest concentration (30 µM). In order to address this discrepancy, we assessed the levels of PrP mRNA by quantitative real-time PCR (RT-PCR). We found that treatment with SM875 did not decrease PrP mRNA in the different cell lines even at the highest concentrations, confirming the original rationale that a compound targeting a folding intermediate of PrP should lower its expression at a post-translational level (Figure 4D). Importantly, only in N2a cells, we observed a robust (>500% at 30 µM), dose-dependent increase of PrP mRNA upon treatment with SM875, which directly explains the discrepancy observed at the protein level. This observation also suggests that the compound triggers an over-production of PrP mRNA in these cells, likely representing a compensatory response aimed at restoring PrP levels. The compound showed variable intrinsic toxicity in the different cell lines, with more prominent cytotoxicity in PrP-transfected HEK293 cells, and progressively lower toxicity in untransfected HEK293, N2a, ZR-75, and L929 cells, respectively (Supp. Figure 10). To further corroborate the observed effects of SM875 on PrP expression, we checked whether the compound also decreases the amount of the protein at the cell surface. HEK293 cells stably transfected with an EGFP-tagged PrP construct were exposed to SM875, and the localization of PrP was monitored by detecting the intrinsic green fluorescence of EGFP. In control conditions, EGFP-PrP localizes almost entirely in the Golgi apparatus and at the plasma membrane, with the latter giving rise to a typical “honeycomb-like” staining of the cell surface (Figure 4E) ^61^. Compounds altering PrP trafficking, as the phenothiazine derivative chlorpromazine, have previously been shown to alter such localization pattern ^62^. Incubation with SM875 for 24 h induced a drastic reduction of cell surface EGFP-PrP at concentrations as low as 1 µM. Collectively, these data confirm that SM875 selectively reduces the expression of PrP at the post-translational level and in a cell-independent fashion.

**Figure 4.**
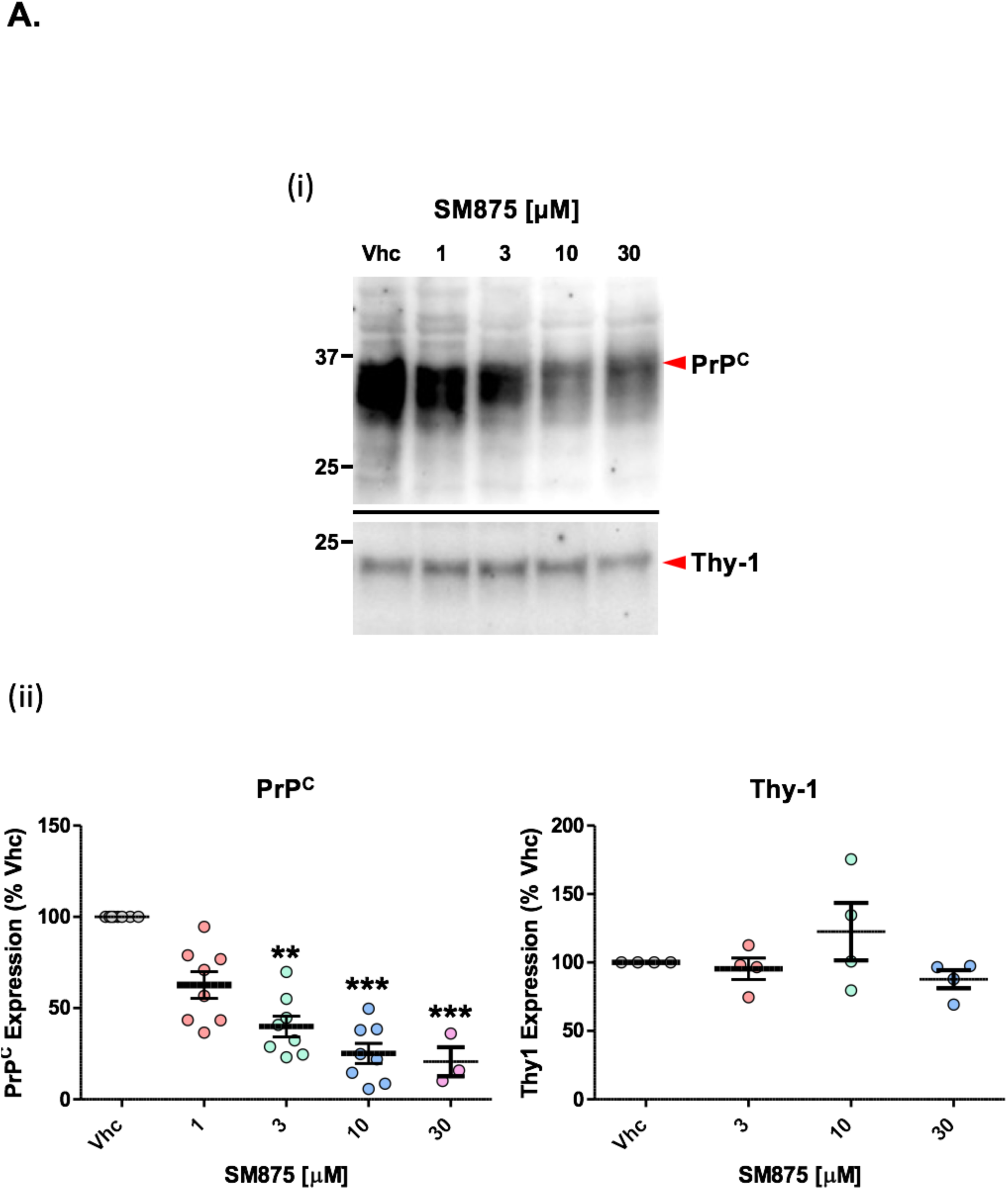
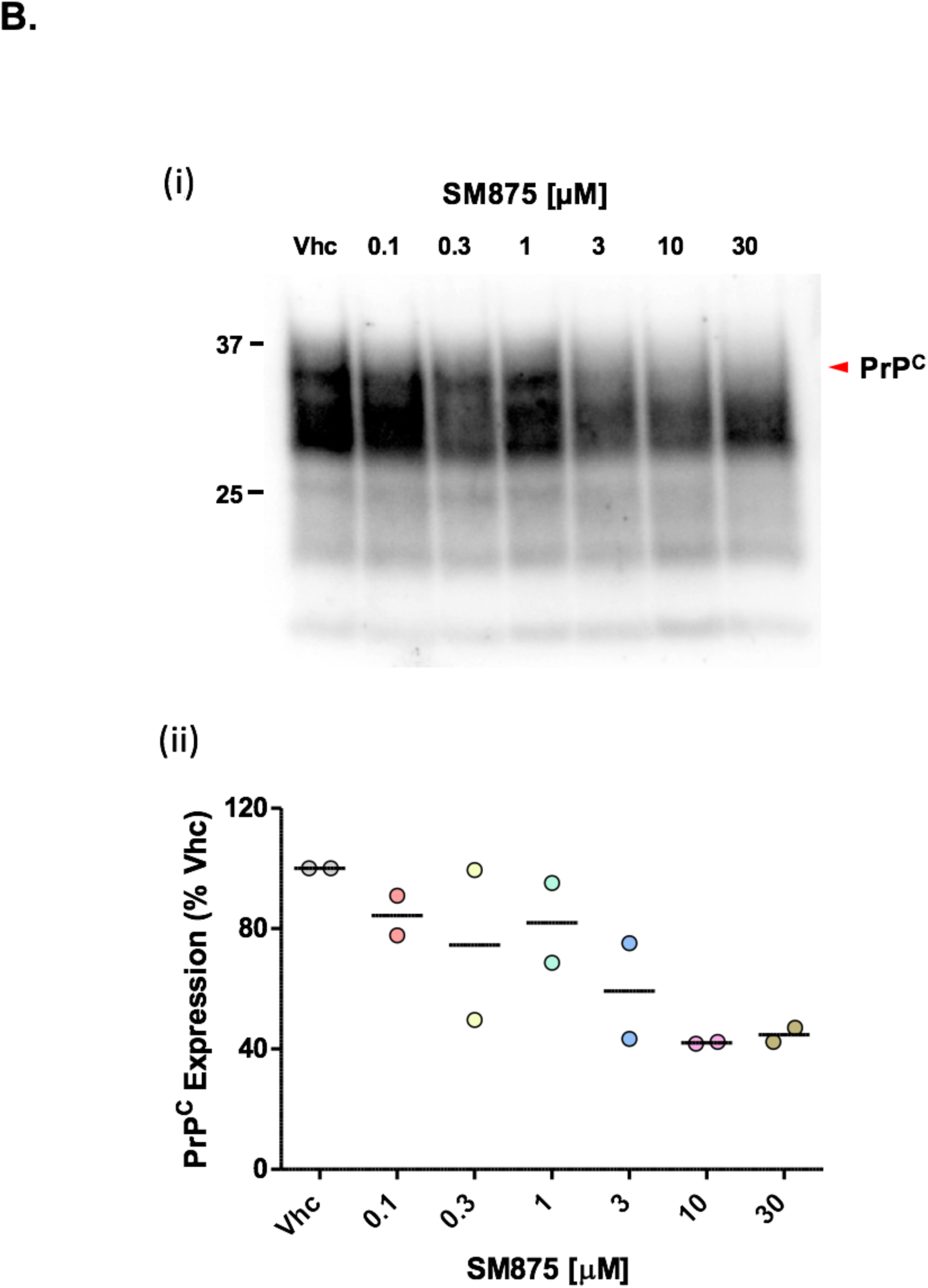
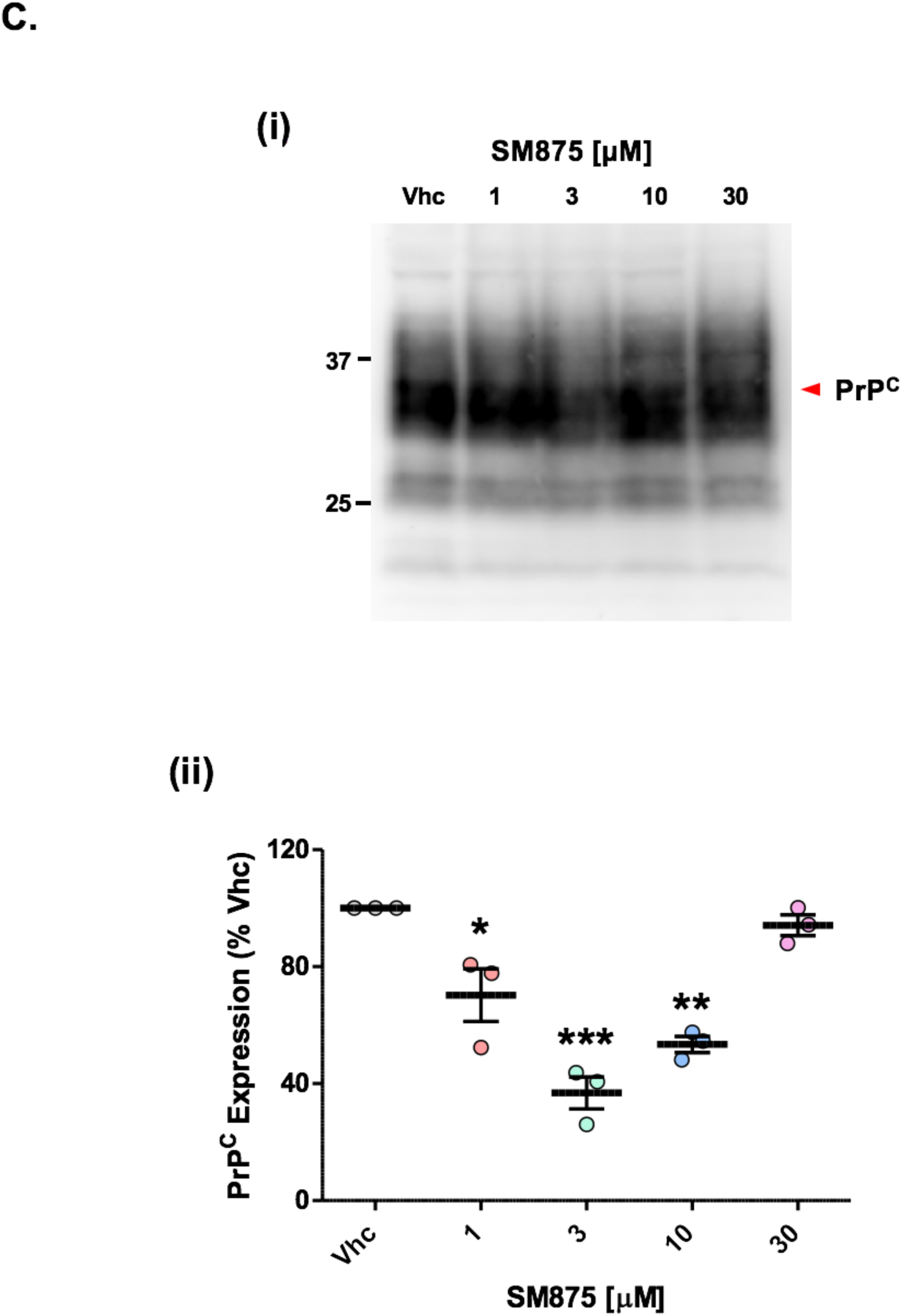
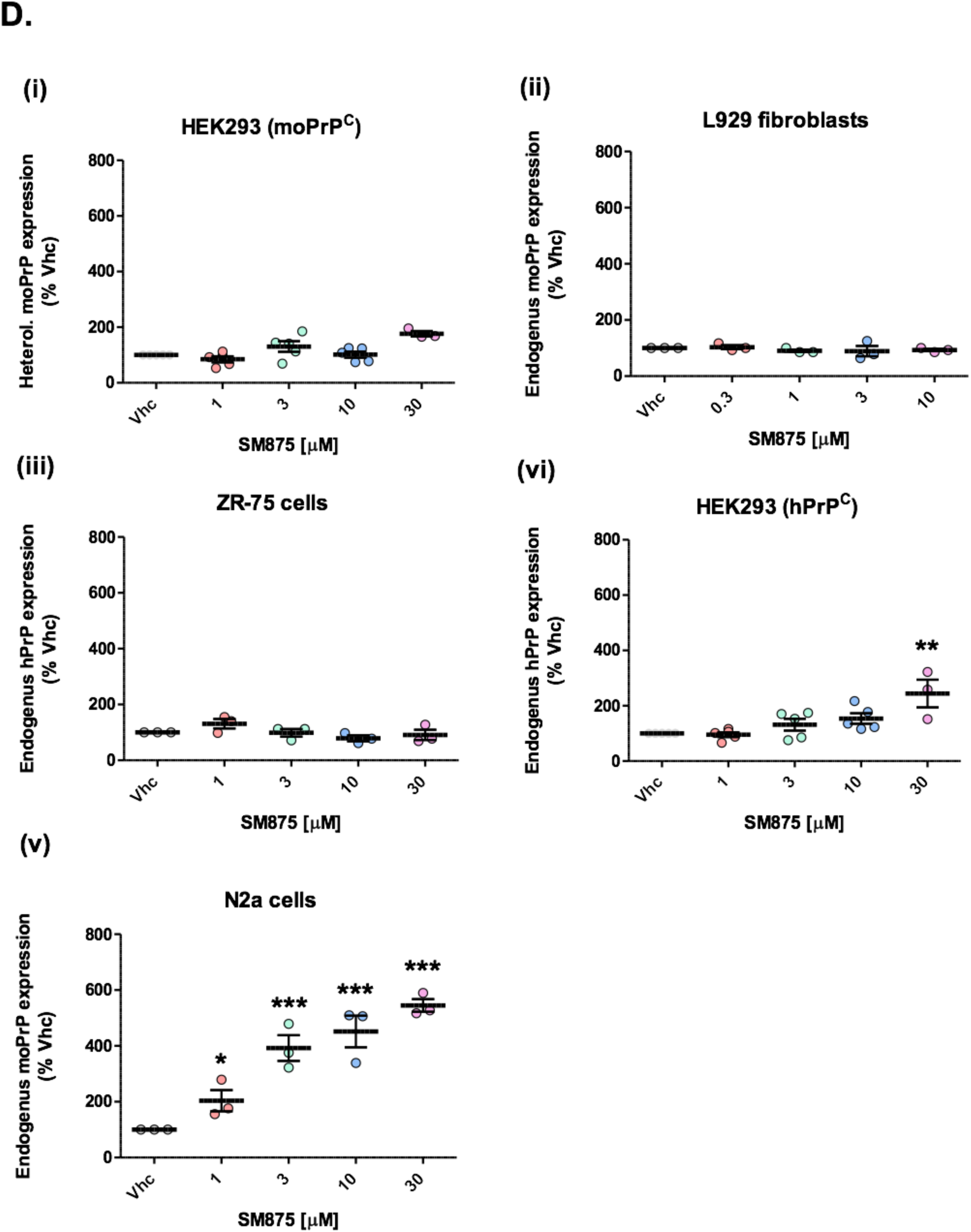
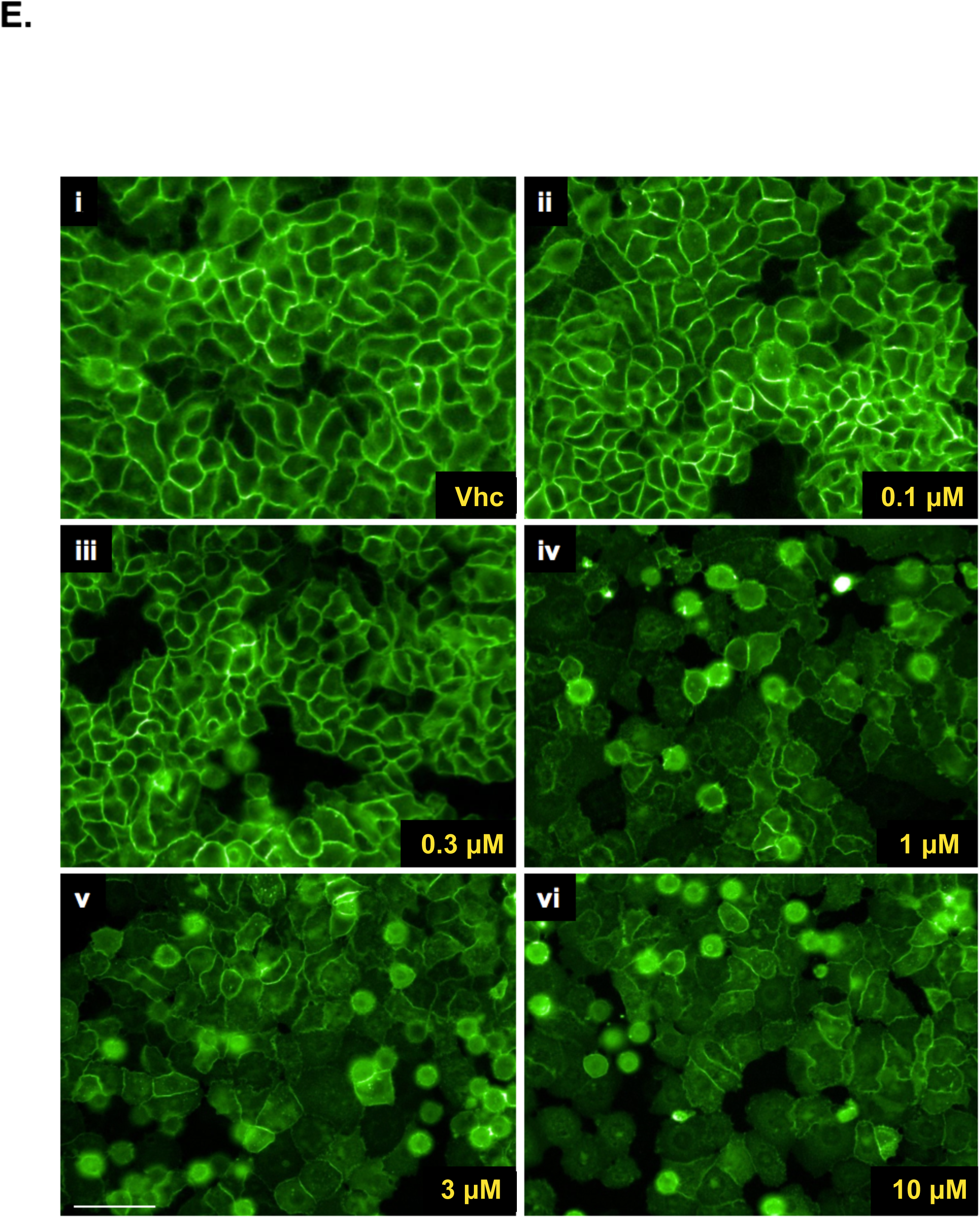
SM875 lowers the expression of PrP at a post-translational level in different cell lines. **A**. ZR-75 cells were exposed to different concentrations of SM875 (indicated) or vehicle (DMSO, volume equivalent) for 48 h, lysed, and analyzed by western blotting. Signals were detected by using a specific anti-PrP (D18) or anti-Thy-1 primary antibodies, relevant HRP-coupled secondary antibodies, and revealed using a ChemiDoc Touch Imaging System. Red arrows indicate the expected sizes of mature, fully glycosylated forms of PrP and Thy-1; **(i)** In ZR-75 cells, SM875 shows a dose-dependent lowering effect of the expression of PrP, in a concentration range of 1-10 µM, although the effect is not accompanied by the appearance of the lower molecular weight bands, as observed in HEK293 cells. The compound shows no effect on Thy-1 levels when tested at the same concentrations at which it lowers PrP expression; **(ii)** the graphs show the densitometric quantification of full-length PrP (left) or Thy-1 (right) levels from independent replicates (n = 8 for PrP; n = 4 for Thy-1). Each signal was normalized on the corresponding total protein lane (detected by UV) and expressed as the percentage of vehicle (Vhc)-treated controls (** p < 0.01, *** p < 0.005). **B**. L929 fibroblasts were exposed to different concentrations of SM875 (indicated) or vehicle (DMSO, volume equivalent) for 48 h, lysed and analyzed by western blotting. Signals were detected by using a specific anti-PrP (D18) primary antibody, HRP-coupled secondary antibody, and revealed using a ChemiDoc Touch Imaging System. The red arrow indicates the expected size of mature, fully glycosylated PrP; **(i)** In these cells, SM875 shows a dose-dependent lowering effect of the expression of PrP, in a concentration range of 3-10 µM, accompanied by the appearance of low molecular weight bands, similarly to HEK293 cells; **(ii)** the graph shows the densitometric quantification of full-length PrP levels from two independent replicates. **C**. N2a cells were exposed to different concentrations of SM875 (indicated) or vehicle (DMSO, volume equivalent) for 48 h, lysed and analyzed by western blotting. Signals were detected by using a specific anti-PrP (D18) primary antibody, HRP-coupled secondary antibody, and revealed using a ChemiDoc Touch Imaging System. The red arrow indicates the expected size of mature, fully glycosylated PrP; **(i)** In N2a cells, SM875 shows a dose-dependent lowering effect of the expression of PrP at 1-10 µM. However, in contrast to the other cell lines, the compound showed no effect at 30 µM; **(ii)** the graph shows the densitometric quantification of full-length PrP levels from independent replicates (n = 3). Each signal was normalized on the corresponding total protein lane (detected by UV) and expressed as the percentage of vehicle (Vhc)-treated controls (* p < 0.05, ** p < 0.01, *** p < 0.005). **D**. In light of the observed effects of SM875 on the amount and intracellular localization of PrP, we sought to test whether the compound could act by decreasing the mRNA of the protein. The levels of PrP mRNA upon treatment with SM875 was evaluated by RT-PCR in the same cell lines and at the same concentrations (indicated) used to evaluate the PrP-lowering effects of the compound. Following treatments, cells were harvested, and RNA was extracted. An aliquot corresponding to 800 ng of total RNA per sample was reverse transcribed using High Capacity cDNA Reverse Transcription kit (Applied Biosystems), according to the manufacturer’s instructions. Quantitative RT-PCR was performed in a CFX96 TouchTM thermocycler (Bio-Rad), using PowerUpTM SYBR Green Master mix (Invitrogen) with a program of 40 cycles amplification. Specific forward and reverse primers were used to amplify endogenous or exogenous, mouse or human PrP transcripts (see Materials and Methods). Relative quantification was normalized to mouse or human HPRT (hypoxanthine-guanine phosphoribosyltransferase). Results show that SM875 does not alter the levels of PrP mRNA in transfected HEK293 cells **(i)** or L929 fibroblasts **(ii)**, expressing mouse PrP (moPrP) exogenously or endogenously, respectively, as well as in ZR-75 cells **(iii)**, endogenously expressing human PrP (hPrP). Interestingly, the molecule induces a slight increase of endogenous hPrP mRNA in untransfected HEK293 cells **(iv)**, an effect that is even more prominent on moPrP endogenously expressed in N2a cells **(v)**. This observation suggests a specific compensatory response increasing PrP synthesis in these cells, caused by the SM875-induced, post-translational decrease of PrP. Such an effect likely explains the observed absence of a dose-dependent decrease of PrP in N2a cells upon treatment with SM875 (reported in panel C). The graphs show the quantification of independent replicates (n ≥ 3). Statistical analyses refer to the comparison with vehicle controls (** p < 0.01, *** p < 0.005). **E**. Based on the observed decrease of PrP expression upon SM875 treatment, we sought to test whether the compound also lowers the total amount of the protein at the cell surface. HEK293 cells were stably transfected with a PrP form tagged with a monomerized EGFP molecule at its N-terminus (EGFP-PrP), and incubated with vehicle (DMSO) control or SM875 at different concentrations **(ii-vi)**. Fluorescence of the EGFP protein was then visualized with an Operetta Imaging System (Perkin Elmer). In control (Vhc) conditions, EGFP-PrP localizes almost entirely at the plasma membrane, giving rise to a typical “honeycomb-like” staining of the cell surface. Conversely, incubation with SM875 for 24 h induced a drastic reduction of cell surface EGFP-PrP, at the same concentrations (1-10 µM) at which the molecule decreases PrP expression. These results are consistent with the idea that SM875 affects PrP expression by altering its biosynthesis. Scale bar 50 µM.

### SM875 induces the degradation of PrP by the lysosomes

Proteins trafficking through the ER that fail to fold properly are usually degraded to prevent the accumulation of toxic aggregates. Most of these aberrantly folded polypeptides undergo retro-translocation into the cytosol, poly-ubiquitination, and degradation by the 26S proteasome, a process known as ER-associated degradation (ERAD) ^38,39,63-65^. While ERAD is the main degradation pathway for most misfolded ER proteins, GPI-anchored proteins like PrP are primarily cleared by the so-called ER-to-lysosome-associated degradation pathway (ERLAD) ^66-68^. For example, expression of disease-associated, aggregation-prone mutants of PrP in different cells increases several markers of the autophagy-lysosomal pathway, such as the autophagosome-specific marker microtubule-associated protein 1A/1B-light chain 3-II (LC3-II) ^58^. Here, we sought to test whether SM875 lowers PrP levels by inducing its lysosomal degradation. First, by comparing PrP-transfected and untransfected HEK293 cells, we observed that SM875 (10-30 µM) increases LC3-II levels in a PrP-dependent manner (Figure 5A). Next, we directly tested the role of lysosomal clearance for the observed PrP-lowering effect of SM875. Since previous data showed that inhibiting degradation pathways in transfected cells could possibly generate over-transcriptional artifacts of the exogenous genes ^67,69^, for these experiments, we relied on ZR-75 cells, expressing PrP endogenously. We found that autophagy inhibitor Bafilomycin A1 completely rescues SM875-induced PrP decrease in these cells (Figure 5B). Of note, the natural disaccharide trehalose, reported to increase de novo formation of autophagosomes, did not alter the PrP-lowering effects of SM875 (Supp. Figure 11) ^70^. Collectively, these data indicate that SM875 promotes the degradation of PrP by the autophagy-lysosomal pathway.

**Figure 5.**
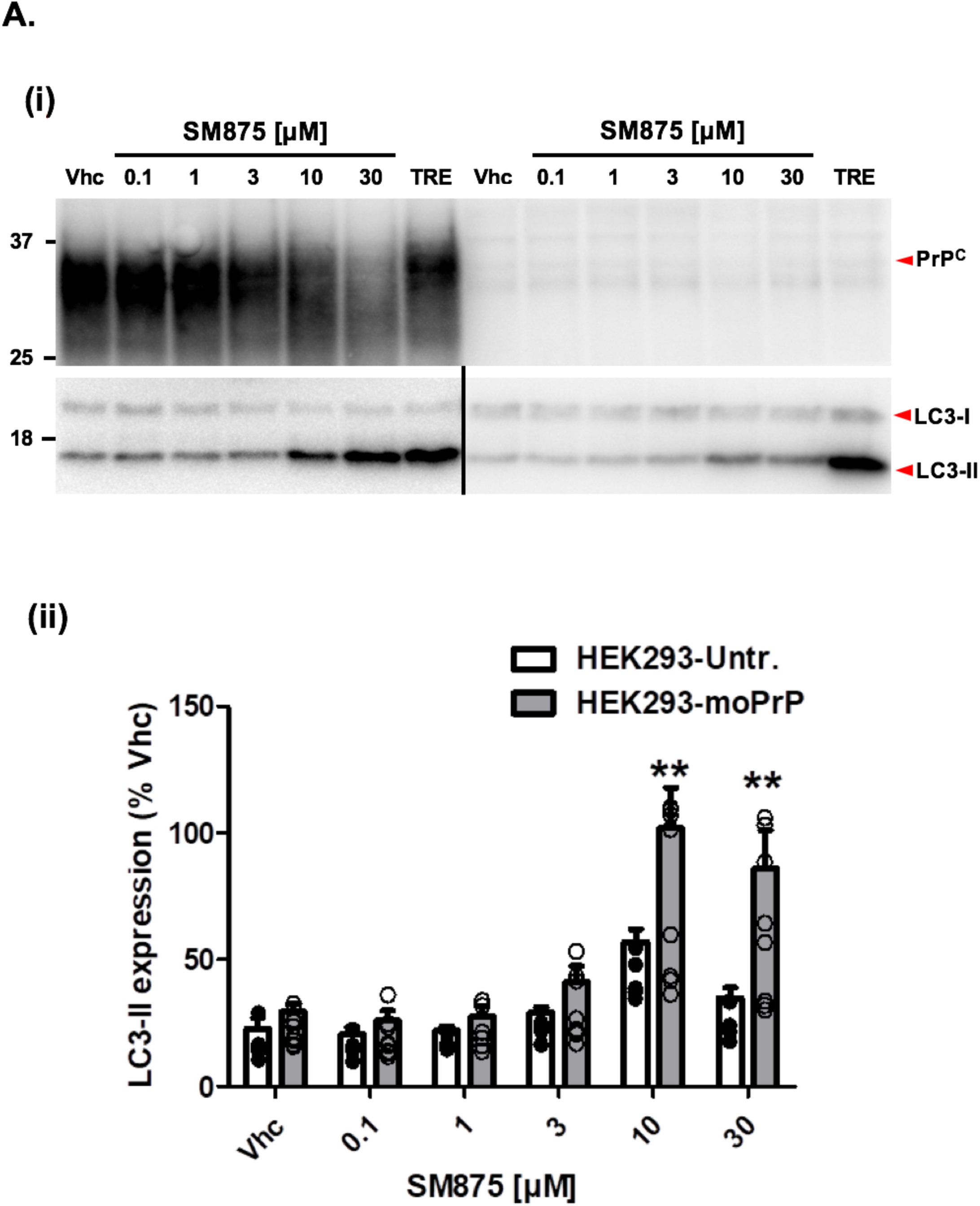
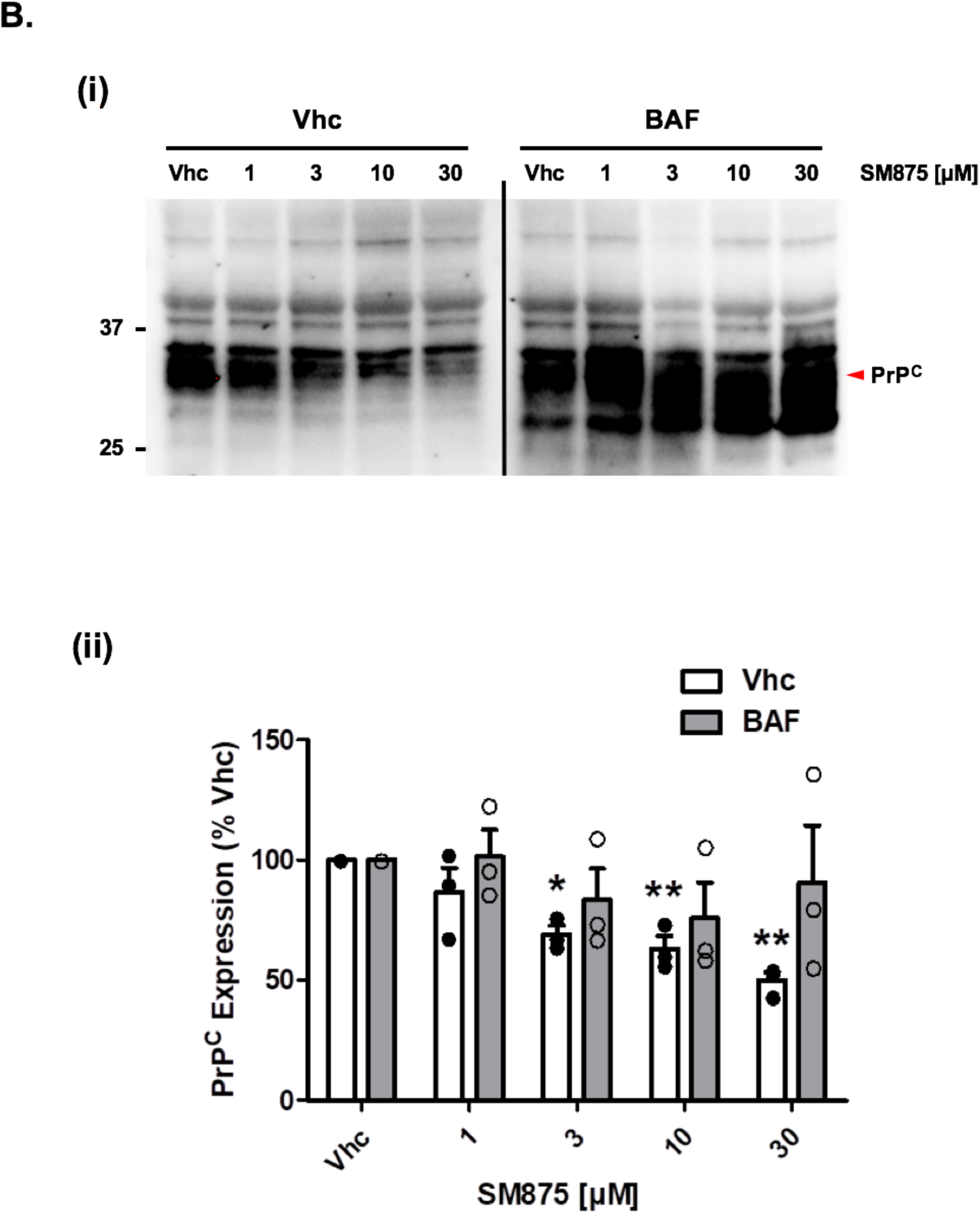
SM875 decreases PrP expression via the autophagy-mediated lysosomal degradation. **A**. Based on the evidence that SM875 lowers PrP expression by acting at a post-translational level, we sought to directly test the hypothesis that the compound promotes the degradation of the protein. Previous data indicate that newly generated misfolded PrP molecules carrying disease-associated mutations are re-routed from the ER to lysosomal degradation, an event associated with the increase of autophagy-lysosomal markers. During the formation of autophagosomal membranes, the cytosolic form of LC3 (known as LC3-I) is conjugated to phosphatidylethanolamine (known as LC3-II) and then recruited to autophagosomal membranes. Consequently, the quantification of LC3-II level is a direct method for monitoring autophagy activation. Since our transfected HEK293 cells highly over-express PrP (as shown in the upper panel), we hypothesized that a molecule promoting its degradation by the ER-associated autophagy could lead to an overall increase of LC3-II. Thus, we treated PrP-expressing as well as untransfected HEK293 cells with different concentrations of SM875 (indicated) or vehicle (DMSO, volume equivalent) and detected PrP, LC3-I, and LC3-II by western blotting (indicated by the red arrows, **i**). As a positive control, we treated cells with 100 µM trehalose (TRE), a disaccharide previously reported to induce autophagy. Consistently, we found that TRE strongly elevates LC3-II levels in both transfected and untransfected HEK293 cells. However, SM875 increases the amount of LC3-II in a dose-dependent fashion almost exclusively in PrP-expressing cells, and not in untransfected HEK293 cells. These results are coherent with the hypothesis that SM875 promotes the degradation of PrP into the lysosomal compartment through an autophagy-mediated mechanism; **(ii)** graphs show the densitometric quantification of LC3-II from independent replicates (n = 8 for PrP-expressing HEK293 cells; n =5 for untransfected HEK293 cells). Each signal was normalized on the corresponding total protein lane (detected by UV) and expressed as the percentage of vehicle (Vhc)-treated controls (** p < 0.01). **B**. In order to corroborate the direct involvement of the lysosomal compartment in the SM875-induced decrease of PrP, we sought to test the effect of the compound in the presence or absence of bafilomycin A1 (BAF, 10 µM), a macrolide antibiotic known to interfere with the autophagy-lysosomal degradation by inhibiting the acidification of the lysosomal vesicles through its interaction with the proton pump V-ATPase. Since blocking protein degradation in transfected cell lines could lead to confounding artifacts ^69^, for these experiments, we employed ZR-75 cells, which express hPrP endogenously. As expected, in the absence of BAF, SM875 induces the decrease of PrP in a dose-dependent fashion, as evaluated by western blotting (**i**). Importantly, co-treatment with BAF fully counteracted the PrP-lowering effects of SM875; (**ii**) the graphs show the densitometric quantification of full-length PrP from independent replicates (n = 3). Each signal was normalized on the corresponding total protein lane (detected by UV) and expressed as the percentage of vehicle (Vhc)-treated controls (** p < 0.01).

### SM875 acts exclusively on nascent PrP

Based on the initial hypothesis that SM875 acts on an intermediate appearing along the folding pathway of PrP, we decided to rule out the possibility that the molecule instead targets native PrP. First, we used dynamic mass redistribution (DMR), a technique previously employed to characterize small molecule ligands of PrP ^71^. In contrast to Fe^3+^-TMPyP, an iron tetrapyrrole known to interact with PrP ^71^, SM875 (0.1-100 µM) showed no detectable binding to either human full-length (23-231) or mouse N-terminally truncated (111-230) recombinant PrP molecules (Figure 6A). These data indicate that the compound has no detectable affinity for natively folded PrP. In order to further validate this conclusion in a cell system, we turned to previously described RK13 cells expressing mouse PrP in a doxycycline-inducible fashion (Supp. Figure 12) ^72^. In the first set of experiments, we turned on the expression of PrP with doxycycline, in the presence of SM875 (10 µM), brefeldin-1A (10 µM), an inhibitor of the ER-to-Golgi protein trafficking ^73^, or vehicle control (DMSO, volume equivalent). Cells were then lysed and analyzed by western blotting at different time points (2, 4, 8, or 24 h, Figure 6B panel i; Supp. Figure 13A). As expected, in contrast to the typical size of full-length, diglycosylated PrP (∼35 kDa) obtained in the control samples, brefeldin-1A induced the accumulation of an immature, low molecular weight band (∼20 kDa), previously described to correspond to a N-terminally-cleaved PrP molecule formed by a lysosomal-dependent event ^74^. Interestingly, SM875 induced the accumulation of a similar PrP band, compatible with its hypothesized activity of targeting a folding intermediate of the protein (Figure 6B, panel i). Next, we sought to directly test the effect of the molecule on pre-synthesized, mature PrP. To pursue this objective, we induced the expression of PrP for 24 h with doxycycline. We then removed the inducer, waited 4 h, treated the cells with SM875 or vehicle control, and analyzed PrP content at different time points (5, 19, and 24 h). In this case, we found no difference between compound- and control-treated samples (Figure 6B, panel ii; and Supp. Figure 13B). The results were further validated by high content imaging analysis of immunostained PrP, which showed that in the same doxycycline-induced RK13 cells, SM875 causes the rapid (as early as 4 h) accumulation of PrP species in intracellular compartments, preventing their correct delivery to the cell surface (Figure 6C). Collectively, these data show that SM875 acts exclusively on nascent, immature PrP molecules, while the compound exerts no effect on pre-synthesized, mature PrP.

**Figure 6.**
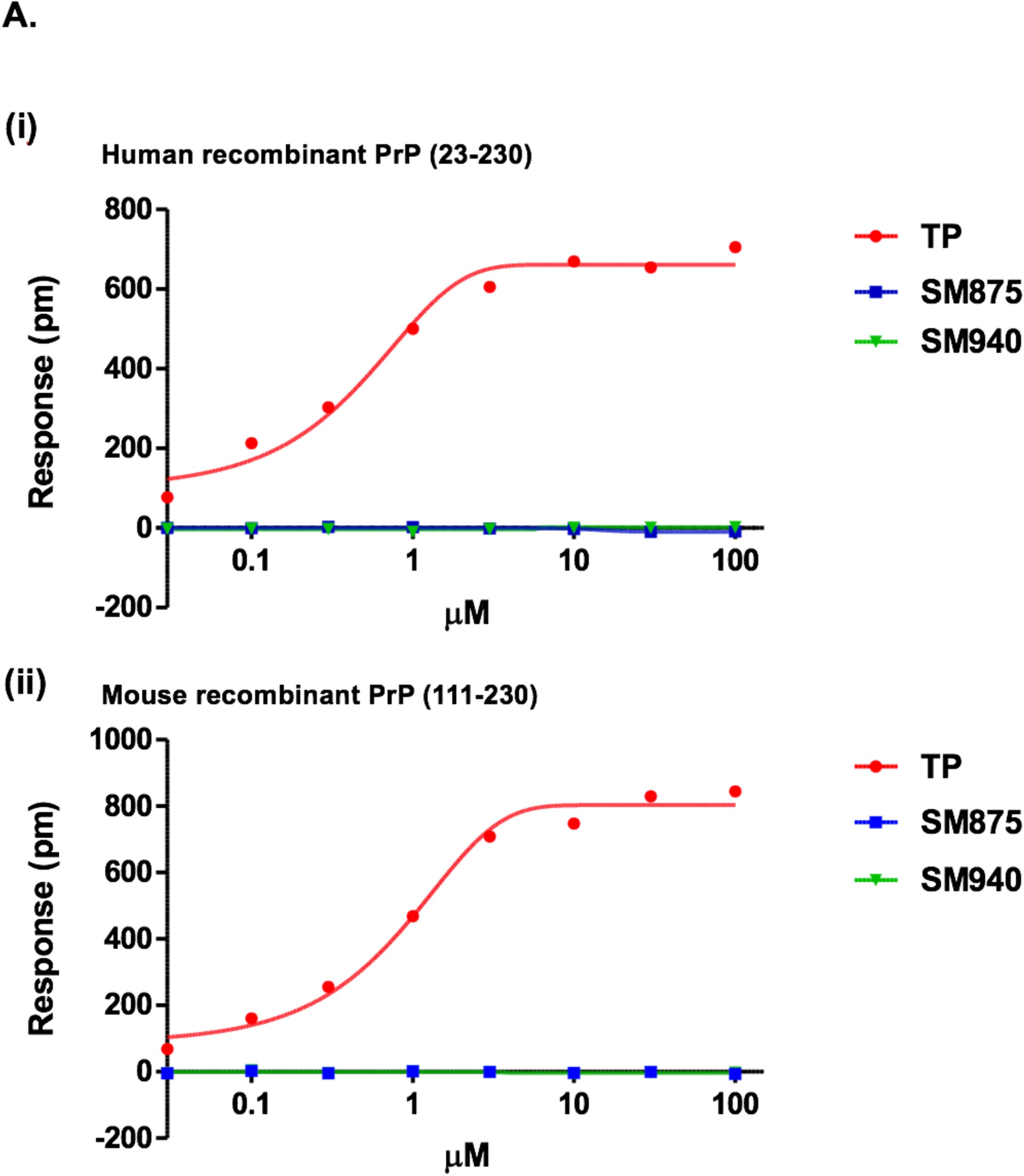
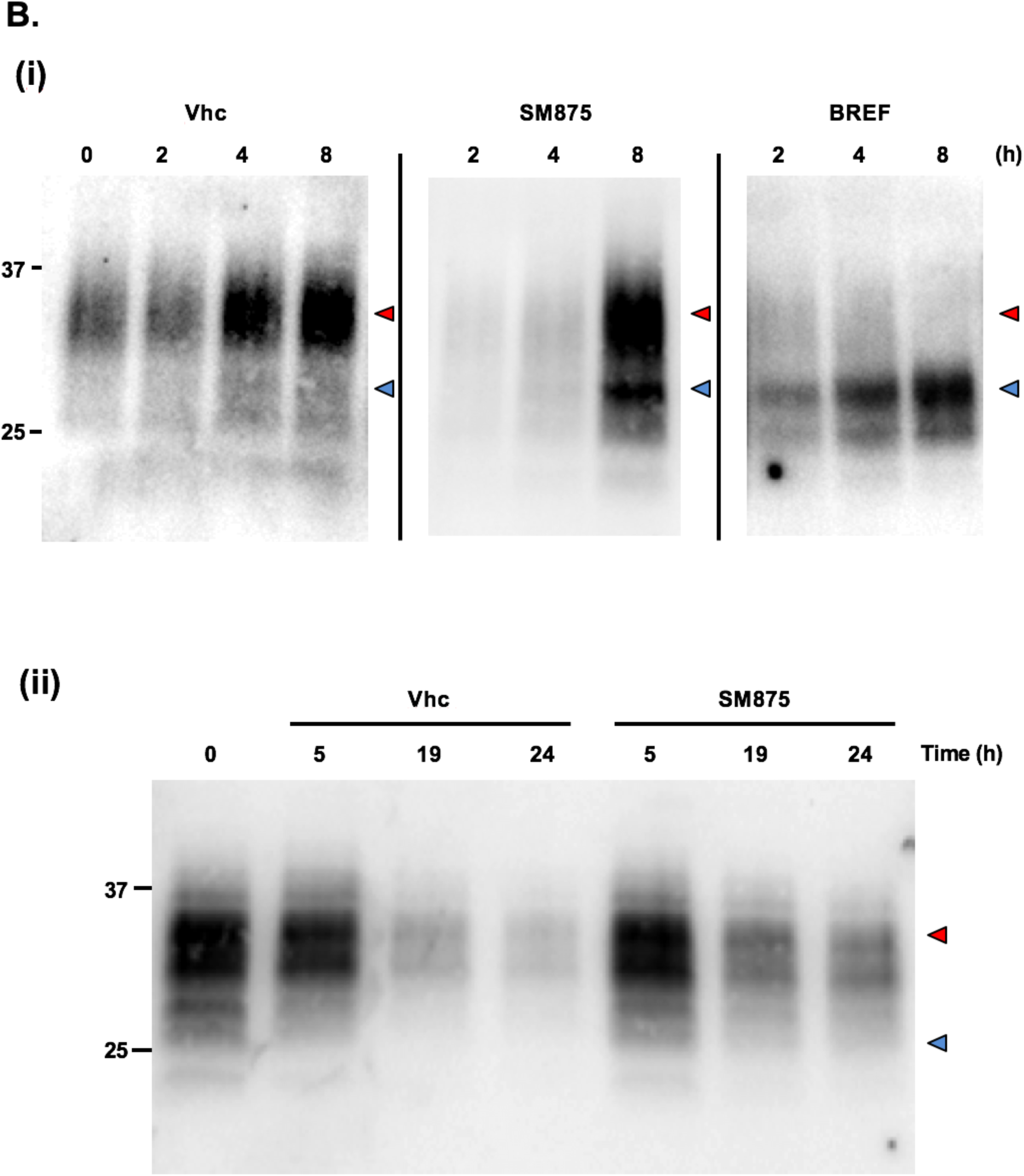
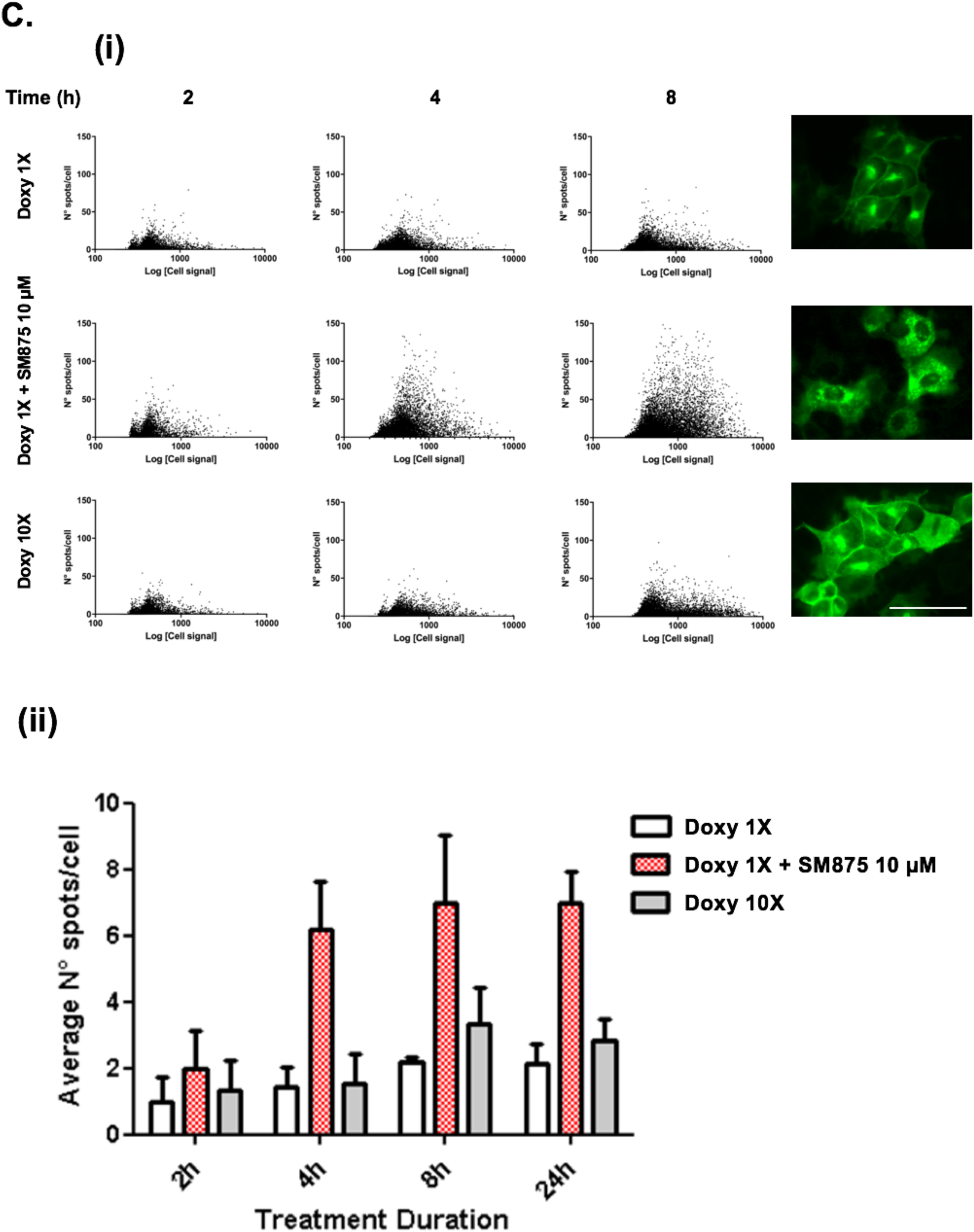
SM875 acts exclusively on non-native, nascent synthesized PrP. **A**. We employed DMR, a biophysical technique previously used to test the interaction of small molecules to recombinant PrP, to assess whether SM875 has an affinity for the native conformation of the protein. Different concentrations (0.03 -100 μM) of SM875, SM940 or PrP ligand Fe^3+^-TMPyP (TP), used as a control, were added to label-free microplate well surfaces on which either **(i)** full-length, human recombinant PrP (23-230) or N-terminally deleted mouse recombinant PrP (111-230) had previously been immobilized. Measurements were performed before (baseline) and after adding each compound. The output signal for each well was obtained by subtracting the signal of the protein-coated reference area to the signal of the uncoated area. The final response (pm) was obtained by subtracting the baseline output to the average of three final output signals. All signals were fitted (continuous lines), when possible, to a sigmoidal function using a 4PL non-linear regression model. As expected, TP shows a detectable affinity for both full-length, human, and N-terminally deleted, mouse PrP molecules, with an affinity in sub-micromolar range (for full-length PrP, K_d_ = 0.67 ± 0.05, R^2^=0.99). Conversely, we failed to detect an interaction with PrP molecules for SM875 or SM940. These results indicate that SM875 (and SM940) are not ligands of native PrP. **B**. In order to dissect the effect of SM875 on nascent vs mature, native PrP molecules, we turned to RK13 cells expressing mouse PrP under control a doxycycline-inducible promoter. **(i)** In the first set of experiments, PrP expression was induced over 8 h, in the presence of SM875 (10 µM), brefeldin-1A (BREF, 10 µM) or vehicle (Vhc) control, samples collected at different time points (indicated) and PrP signals visualized by western blotting. Signals were detected by probing membrane blots with anti-PrP antibody (D18). As expected, in control cells, the expression of full-length PrP (red arrows) increases in a time-dependent fashion. Conversely, a lower molecular weight band (blue arrows) is detected in brefeldin-treated cells. This form had previously been reported to correspond to immature PrP species resulting from the inhibitory action of brefeldin-1A on the ER-to-Golgi transition ^73^. Interestingly, SM875 induces the appearance of the same PrP band upon 8 h from induction, although less potently than brefeldin-1A. These results indicate that SM875 targets nascent PrP molecules; **(ii)** Next, we designed an experiment to test the effect of SM875 exclusively on mature, natively folded PrP. PrP expression was induced for 24 h, in the absence of any additional treatment. Doxycycline was then removed, and after 4 h without inducer, the cells were exposed to SM875 or Vhc control, and subsequently lysed at different time points (indicated). In this experimental setting, cells are exposed to SM875 only when all PrP molecules are synthesized and likely in transit to, or already reached the plasma membrane. In these conditions, normal PrP patterns appear in both compound-treated and Vhc-treated cells. These results indicate that SM875 exerts no detectable effects on pre-synthesized, mature PrP. **C**. To further validate the results, we applied a high-content approach to the same experimental setting described above by analyzing the localization of PrP after immunostaining with an anti-PrP antibody (D18) coupled to an Alexa 488 secondary antibody. (i) The expression of PrP was induced for 2, 4 or 8 h by doxycycline 1x (0.01 mg/mL) or 10x (0.1 mg/mL) and the effect of SM875 (10 µM) incubated together with doxycycline 1x was measured by Harmony software after the image acquisition performed by Operetta Imaging System. Green spots detected in cells were quantified and plotted against the total green fluorescence relative to each cell. While the extended incubation time and doxycycline concentration, es expected, affect the total green fluorescence by increasing the levels of PrP, only SM875 elevates also the number of intracellular spots, preventing the correct delivery of PrP to the cell surface. Representative images on the right were acquired at 8 h of incubation, scale bar 50 µM. (ii) Quantification of the average number of green spots per cell in wells incubated for 2, 4, 8, and 24 h with doxycycline 1X (white bars), doxycycline 10X (gray bars) and doxycycline 1X + SM875 (red bars). While the augmented PrP synthesis induced by doxycycline 10X induces just a modest rise in the number of spots, SM875 significantly enhances the aggregation of PrP. Quantification of at least 3 independent experiments. N° spots/cell below the 10% percentile and above the 90% percentile have been excluded. Average N° spots/cell values have been normalized on doxycycline 1X 2h, considered as a baseline condition. Unpaired *t test*. 4h doxycycline 1X vs doxycycline 1X + 875 *p= 0.0147; 8h doxycycline 1X vs doxycycline 1X + 875 *p=0.0183; 24h doxycycline 1X vs doxycycline 1X + 875 **p= 0.0024.

### SM875 suppresses prion replication in mouse L929 fibroblasts

One of the most solid concepts in prion diseases is that suppressing PrP should abrogate prion replication ^75^. The most direct way to achieve this objective is to silence PrP expression, either through a selective decrease of its synthesis or increase of its clearance. Following the observation that SM875 lowers the expression of PrP by inducing the degradation of a folding intermediate, we tested the ability of the compound to inhibit the replication of the Rocky Mountain Laboratories (RML) prion strain in persistently infected L929 mouse fibroblasts ^76^. We found that SM875 inhibits prion replication in a dose-dependent fashion, decreasing prion loads similarly to anti-prion molecule Fe^3+^-TMPyP, used as a control (Figure 7) ^71^. These results lay the groundwork for the chemical optimization of SM875 into an analog administrable in vivo, which could provide a new therapeutic tool against prion diseases.

**Figure 7.**
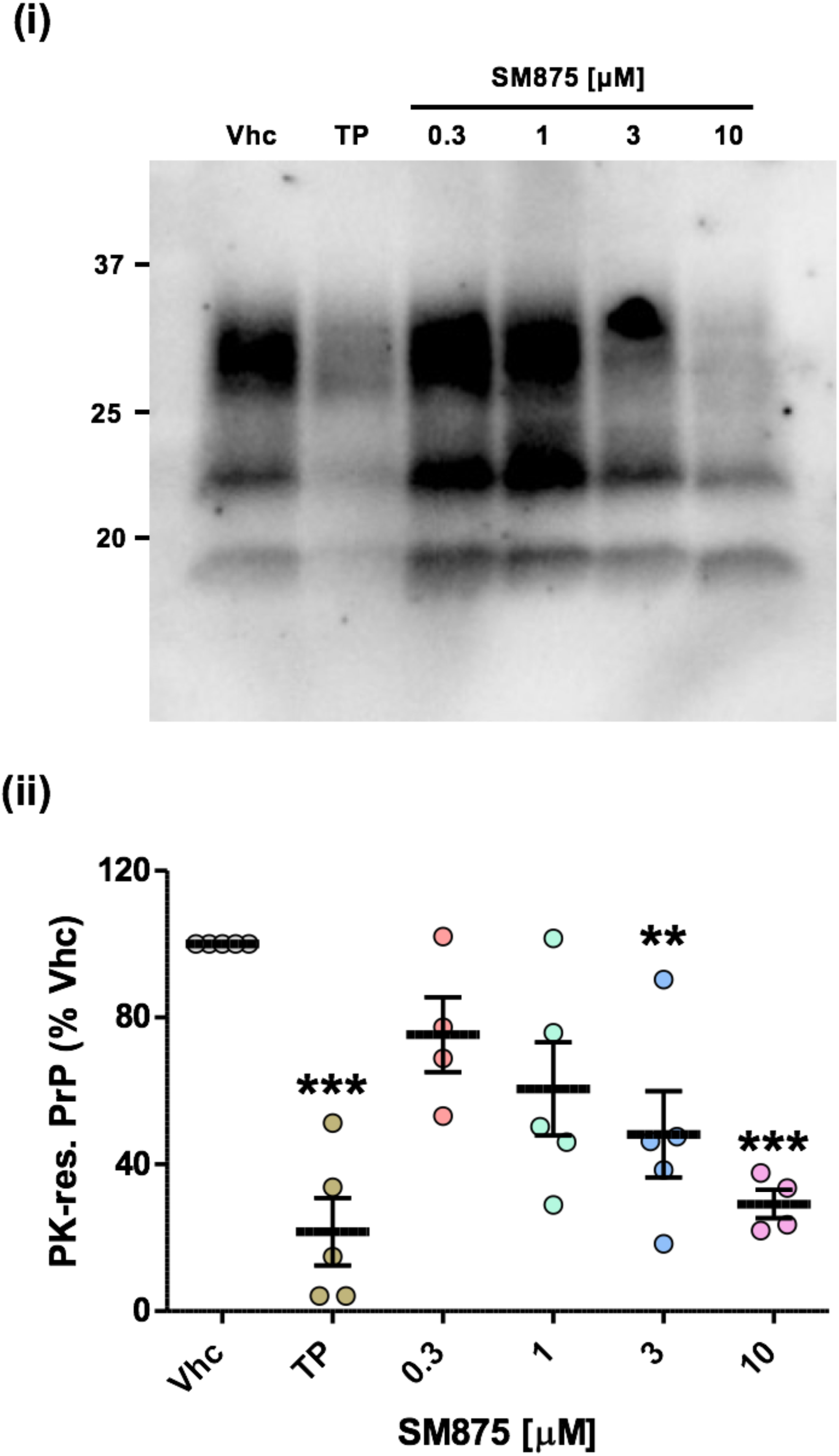
SM875 inhibits prion replication in L929 mouse fibroblasts. Based on the results showing that SM875 consistently lowers PrP expression in a variety of cell lines, we sought to test the ability of the molecule to inhibit the replication of PrP^Sc^ in prion-infected cells. N2a cells, a widely used cell model for prion replication, were not suitable for this experiment, as we observed a compensatory overexpression of PrP mRNA in these cells upon treatment with SM875. Thus, we turned to L929 mouse fibroblasts. These cells were infected with the RML prion strain (derived from a brain extract of a prion-infected mouse) and then propagated for five sequential passages before exposure to SM875 (indicated concentrations), anti-prion compound Fe^3+^-TMPyP (TP, 10 µM) or vehicle (DMSO) for 48 h. PrP^Sc^ loads were then estimated by treating cell lysates with PK and analyzing PrP content by western blotting **(i)**. Signals were detected by probing membrane blots with anti-PrP antibody (D18). Results show that SM875 inhibits prion replication in a dose-dependent fashion, with the maximal effect (obtained at 10 µM) comparable to that of TP; **(ii)** the graph shows the densitometric quantification of PK-resistant PrP species from independent replicates (n = 4). Quantification was obtained by densitometric analysis of the western blots, normalizing each signal on the corresponding PK-untreated lane and expressed as the percentage of vehicle (Vhc)-treated controls (** p < 0.01, *** p < 0.005).

### SM875 induces the aggregation of partially denatured PrP

Despite the different pieces of evidence collected in-silico and in cell-based assays, the formal demonstration that SM875 directly targets a folding intermediate of PrP would require a structural characterization of the complex. Unfortunately, solving the structure of folding intermediates is limited by the transient nature of these protein conformers. In an attempt to overcome this problem, we designed an experimental paradigm aimed at inducing the appearance of non-native conformers of PrP by thermally-induced partial denaturation (Supp. Figure 14). The underlying hypothesis is that the PrP folding intermediate predicted by our in-silico analyses could also be reached from the native state by overcoming an energy barrier so that SM875 could bind and stabilize it. Here, we explored this possibility by trying to crystallize the temperature-induced, semi-denatured PrP in complex with SM875. Recombinant mouse PrP (800 µM) was heated to 45 °C and slowly cooled to 20°C using a thermal cycler, either in the presence or absence of SM875 (2 mM). Unexpectedly, we observed massive precipitation (∼40%) of the protein as soon as SM875 was added to the solution, as assayed by UV absorbance. Conversely, no appreciable precipitation was detected in vehicle-treated controls. The remaining soluble PrP molecules gave rise to “thin needles”-like crystals, appearing after 2-4 days. Importantly, such crystals were obtained irrespectively of whether the protein was incubated with SM875 or vehicle. Crystals were weakly and anisotropically diffracting to 3.7 Å in the best directions, allowing determination of orthorhombic crystal system and unit cell dimensions, highly similar to those reported for apo human PrP in the protein databank (PDB 3HAK) (Supp. Table 6). These data suggest that SM875 targets a thermally-induced PrP population, promoting its precipitation. Conversely, the more native-like PrP molecules do not bind to SM875 and remain in solution, allowing crystallization.

The observed precipitation of partially-denatured PrP induced by SM875 could reflect an intrinsic propensity of a PrP folding intermediate to expose hydrophobic residues normally buried in the native state. In order to better characterize this effect, we performed a detergent-insolubility assay to detect insoluble aggregates of mouse recombinant PrP (111-230) in the presence or absence of SM875 upon temperature shift (25, 37, 45, 55°C; of note, the reported melting temperature for recombinant PrP is approximately 65°C) ^77^. We found that, differently from vehicle-treated samples, SM875 induced the aggregation of recombinant PrP in a temperature-dependent fashion (Figure 8). Importantly, similar results were obtained with compound SM940 but not SM935, respectively found positive and negative for PrP downregulation in cells (Figure 3C and Supp. Figure 15). Collectively, these data add direct evidence for the interaction of SM875 with a non-native PrP conformer.

**Figure 8.**
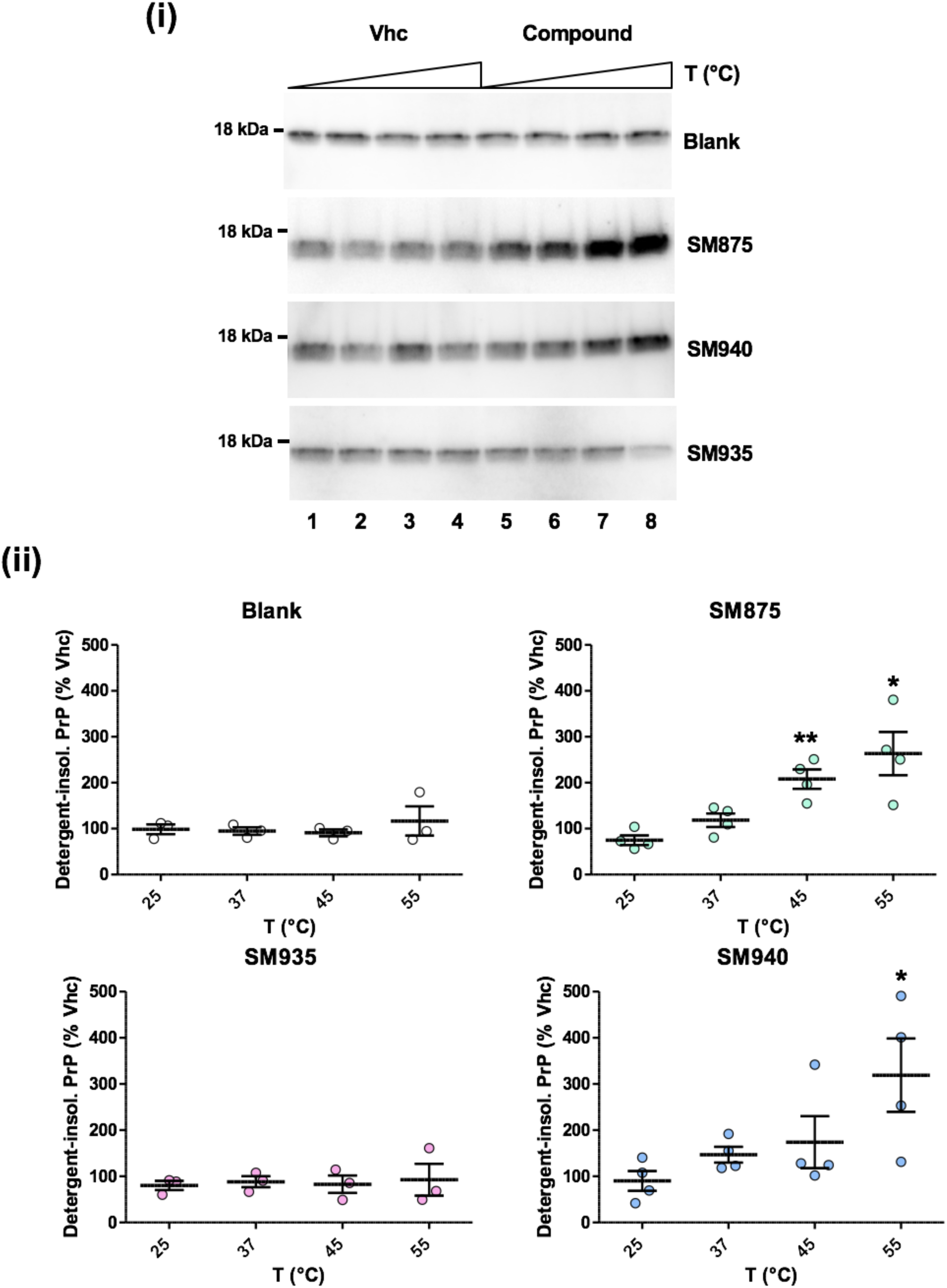
SM875 induces the aggregation of PrP in a temperature-dependent fashion. In order to detect a direct effect of SM875 on the PrP folding intermediate, we implemented the experimental paradigm adopted for crystallization by adding additional temperature points (25, 37, 45 and 55 °C). Recombinant PrP (111-231) was diluted in a detergent buffer (final concentration of 0.5 µM), placed at one of the indicated temperatures and incubated for 1 h with either vehicle (Vhc, lanes 1-4) or (lanes 5-8) assay buffer (blank), SM875 (10 µM), SM940 (10 µM) or SM935 (10 µM). After incubation, samples were subjected to ultracentrifugation, and the resulting detergent-insoluble pellets, corresponding to the aggregated fraction of PrP, were analyzed by western blotting **(i)**. Signals were detected by probing membrane blots with anti-PrP antibody (D18). These experiments revealed a temperature-dependent increase of PrP aggregation upon incubation with SM875 and SM940. Conversely, both the blank control and SM935, a compound found in the initial screening not to exert PrP-lowering effects in cells and used here as a negative control, showed no detectable changes in PrP aggregation after temperature increase; **(ii)** the graphs show a densitometric quantification of recombinant PrP band from independent replicates (n ≥ 3). Each signal was normalized and expressed as the percentage of the corresponding Vhc-treated sample (* p < 0.05, ** p < 0.01).

## DISCUSSION

We have applied computational techniques to shed light on the role of folding intermediates in a previously reported mechanism of regulation of the AR expression. The observation that protein expression may be tuned by acting on folding pathways prompted us to test an entirely novel method for selectively reducing the level of target proteins, which we called PPI-FIT. This technology is based on the concept of targeting folding intermediates of proteins rather than native conformations. The approach is made possible by the recent development of computational algorithms allowing the full atomistic reconstruction of the entire sequence of events underlying the folding of a polypeptide into its native state. In the PPI-FIT method, druggable pockets appearing in specific folding intermediates observed along the folding pathway of a given protein are used as targets for virtual screening campaigns aimed at identifying small ligands for such regions. The underlying rationale is that stabilizing a folding intermediate of a protein with small ligands could promote its degradation by the cellular quality control machinery, which could recognize such artificially stabilized intermediates as improperly folded species. In this manuscript, we applied this technology to PrP, identified a specific intermediate along the folding pathway of the protein, and then carried out a virtual screening campaign that led to the discovery of four different chemical scaffolds all capable of selectively lowering the expression of PrP. Extensive characterization of one of these molecules, called SM875, provided strong experimental support for the notion that targeting folding intermediates could represent a novel pharmacological paradigm to modulate protein expression. From a broader perspective, our data reveal the existence of a previously unappreciated layer of regulation of protein expression occurring at the level of folding pathways.

### Evidence supporting the PPI-FIT mode of action for the identified compounds

The demonstration that the expression of a protein could be regulated by targeting a folding intermediate, the concept underlying the PPI-FIT method, would ultimately require a direct biophysical characterization of the interaction between the identified ligands and the predicted folding conformer. Unfortunately, available high-resolution techniques such as X-ray crystallography or NMR can only be applied to stable molecular species, while protein folding intermediates are transient and highly dynamic by definition ^78^. Conversely, techniques designed to study single molecules, like optical tweezers, or to capture very rapid molecular transitions, such as spectroscopic detection coupled to stopped-flow methods, are intrinsically low resolution. In fact, the existence of a folding intermediate of PrP was previously detected by using similar approaches but never characterized at a structural level ^40-42^. Despite the intrinsic approximations of all in-silico techniques, our computational platform overcomes the *spatio-temporal* resolution limits, allowing us to define the atomistic structure of the putative PrP folding intermediate, as well as to use this information to carry out a virtual screening campaign aimed at identifying small ligands for such conformer. Then, in light of the aforementioned experimental limits, we designed an extensive in vitro and cell-based workflow to validate the PPI-FIT method. First, we identified four compounds, displaying different chemical scaffolds, all capable of selectively lowering the expression of PrP. Next, we found that SM875, the most potent among the positive hits, reduces PrP loads in several different cell lines, either expressing the protein endogenously or exogenously, without decreasing PrP mRNA levels. Our data also show that SM875 promotes the degradation of nascent PrP molecules by the autophagy-lysosomal pathway, while the molecule neither binds nor exerts any effect on mature, cell surface PrP. Finally, in an attempt to co-crystallize the PrP folding intermediate bound to SM875, we found that the small molecule induces the precipitation of PrP species appearing upon mild thermal unfolding. Collectively, even though we do not formally demonstrate the interaction of SM875 with the predicted binding pocket, our data provide strong experimental support for the notion that the compound acts as by targeting a folding intermediate of PrP (Figure 9).

**Figure 9.**
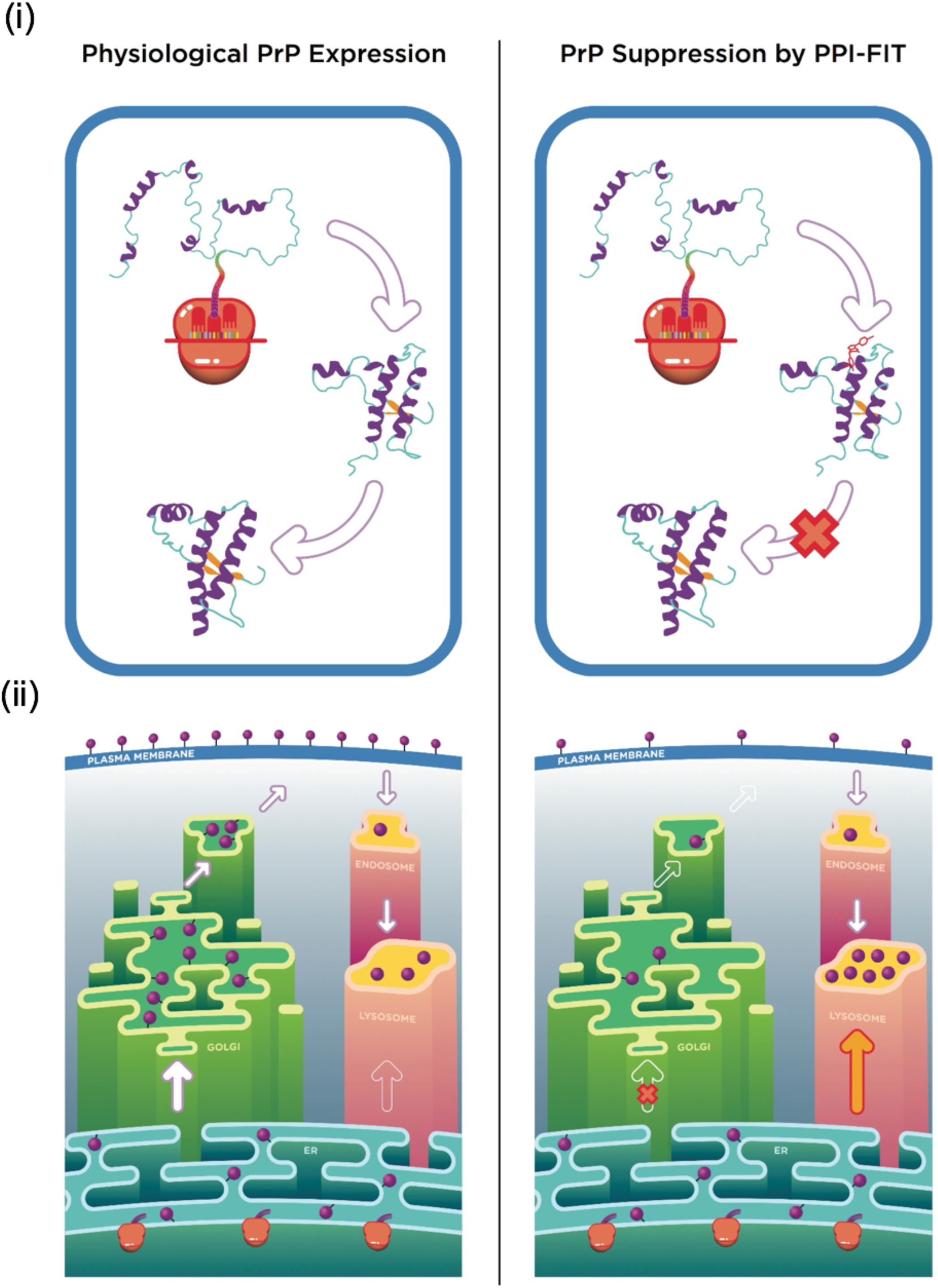
Model for the PPI-FIT-based suppression of PrP. The model schematically illustrates the rationale underlying the PPI-FIT method applied to PrP. (i) The schematics highlight PrP folding in the presence (right) or absence (left) of a small molecule targeting the PrP intermediate. (ii) PrP follows a typical expression pathway for GPI-anchored proteins (left panel). The polypeptide is directly synthesized into the lumen of the ER, once properly folded and post-translationally modified it traffics to the Golgi apparatus, where sugar moieties are matured, and then delivered to the cell surface. From the plasma membrane, PrP molecules could enter into the endosomal recycling pathways, and eventually recycled into the lysosomes. Aberrantly folded conformers of PrP, as the case for some previously reported mutant, could be re-routed directly from ER to the lysosomes. In the PPI-FIT method, a small molecule interferes with PrP expression by binding to a folding intermediate, which gets delivered to the lysosomes, leading to an overall decrease of the protein at the cell surface.

### Therapeutic potentials of SM875

Prion diseases represented a convenient experimental ground for testing the PPI-FIT approach, as compelling evidence indicates that prion replication and toxicity can be inhibited by targeting a single protein (PrP) ^24,25,44,45^. Theoretical options to achieve this goal include the ablation of the PrP gene, for example by using the CRISPR technology ^79,80^, decreasing its synthesis with specific antisense oligonucleotides (ASOs) ^27^, or acting directly on cell surface PrP ^81,82^, for example with ligands stabilizing the native fold and inhibiting its conversion into PrP^Sc^ ^50,77^. Despite encouraging results obtained with ASOs recently, so far, none of these approaches has been translated into effective therapies for prion diseases. Moreover, multiple unsuccessful attempts to identify small molecules specific for PrP suggest that this protein could be an undruggable target ^49,50^. Conceptually, a compound like SM875 capable of selectively promoting the degradation of PrP by targeting a folding intermediate, could be the ideal candidate to tackle prion diseases pharmacologically, and possibly synergize with other PrP-silencing strategies. Our data show that SM875 alters the correct maturation of the PrP polypeptide, similarly to ER-to-Golgi trafficking inhibitor brefeldin 1A. Moreover, in prion-infected cells, the PrP-lowering activity of SM875 produces a strong inhibition of prion replication, comparable to that of the potent anti-prion compound Fe^3+^-TMPyP. However, in contrast to brefeldin 1A and Fe^3+^-TMPyP, which are known to be rather non-specific ^50,83,84^, the effect of SM875 is selective for PrP, as demonstrated by the lack of alterations of other two GPI-anchored proteins (NEGR-1 and Thy-1). These results indicate that the further development of PPI-FIT-derived molecules into compounds administrable in vivo and their subsequent validation in animal models of prion diseases could represent fundamental steps to move these new classes of anti-prion compounds to the clinical phase.

### Pharmacological relevance of targeting folding intermediates

A major aspect of the PPI-FIT method is that it capitalizes on the cellular quality control machinery to promote the degradation of the target polypeptide. A similar rationale is exploited by emerging pharmacological strategies like the proteolysis targeting chimeras (PROTACs) ^85,86^. This approach builds on the principle of designing bi-functional degraders composed by two covalently-linked chemical moieties, one interacting with the target protein and the second engaging the E3 ubiquitin ligase, which then targets the polypeptide to degradation by the proteasome. Notably, two PROTAC compounds against the estrogen receptor and the AR have recently reached the clinical phase (trial numbers NCT03888612 and NCT04072952). However, compounds like PROTACs target native structures of proteins, therefore suffering from the same limits as classical pharmacological approaches when applied to undruggable targets like PrP. The PPI-FIT method theoretically expands the range of druggable conformations of a polypeptide by targeting alternative structures explored by the protein along its folding pathway. In fact, the PrP folding intermediate provided a solvent-exposed, druggable pocket hidden in the native conformation. This feature may result in an additional advantage: a common problem in pharmacology is posed by proteins with a high structural similarity of their native conformation, a factor that greatly decreases the chances of developing selective drugs. However, proteins with highly similar three-dimensional native folds do not necessarily share folding pathways, and even proteins with identical folding pathways may differ for the lifetimes of folding intermediates, which would then display different affinities for the same compound ^35,87^. Thus, the PPI-FIT approach could intercept structurally and/or kinetically-different folding intermediates and ultimately allow the identification of selective ligands even in the case of highly similar proteins. Finally, it will be interesting to evaluate in the future whether existing drugs that lower the expression of specific proteins by unknown mechanisms could be acting in a PPI-FIT-like manner by targeting folding intermediates.

### Do folding intermediates represent a layer of regulation of protein expression?

The observation that the expression of the AR could be modulated by altering the metabolism of non-native conformers of the protein appearing along the folding pathway suggests that similar mechanisms of regulation could be exploited in biology to manipulate the expression of several other proteins. Interestingly, such a mechanism could be directly interconnected with the protein quality control machinery, potentially conferring an additional degree of complexity for the spatial and temporal regulation of proteostasis. Regulating the homeostasis of a folding intermediate of a protein, rather than just acting on transcription, translation or clearance of its mature form, could be energetically favorable and allow the cell to respond to stimuli more rapidly. Similar mechanisms may play important roles in several other disease contexts beyond the one addressed here. For example, altering the folding pathways of specific homeostatic regulators or immunoregulatory factors could be a way for cancer cells to overcome cellular checkpoints, or for pathogens to evade host defenses. On an even broader perspective, the existence of functional folding intermediates of proteins implies that the Darwinian evolution of polypeptide sequences does not act exclusively on native conformations, but also on alternative protein conformers transiently appearing along the folding pathways.

## METHODS

### Bias Functional Methods for Folding Pathways Simulations

#### General Features and Software

The Bias Functional (BF) method is a three-steps procedure that enables the simulation of protein folding pathways at atomistic level of resolution and consists in: (i) generation of denatured condition by thermal unfolding, (ii) productions of folding trajectories starting from the unfolded states and (iii) scoring of the folding trajectories based on a variational principle. The software to perform these simulations relies on the molecular dynamics (MD) engine of Gromacs 4.6.5 patched with the plugin for collective variable analysis Plumed 2.0.2.

#### Generation of Denatured Conditions

Unfolded conformations are generated by thermal unfolding starting from the native structure. This is achieved by performing independent 3-5 ns trajectories of molecular dynamics at high temperature (800K) in the NVT ensemble. For each trajectory, a single denatured conformation is extracted.

#### Generation of Folding Pathways

For each denatured conformation, a set of folding trajectories are generated by employing the ratchet and pawl molecular dynamics (rMD) algorithm. In this scheme, the folding progress is described as a function of a reaction coordinate, defined as *z(X)*:

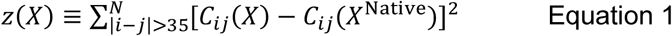

Where *C*_*ij*_*(X)* is the contact map of the instantaneous system configuration and *C*_*ij*_*(X*^*Native*^) Is the contact map in the reference native state. The reference native state is obtained by energy minimizing the experimental structure retrieved from the protein data bank. The *C*_*ij*_*(X)* entries of *z(X)* interpolate smoothly between 0 and 1 according to the following function:

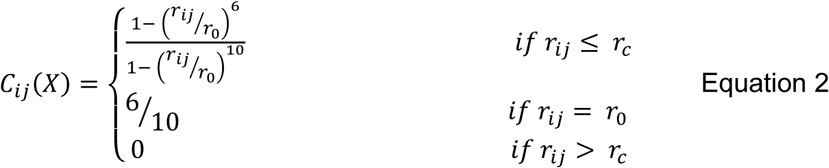

Where *r*_*ij*_ is the Euclidean distance between the *i*_*th*_ and the *j*_*th*_ atom, *r*_*0*_ is a typical distance defying residue contacts (set to 7.5 _Å_) and *r*_*c*_ is a cutoff distance (set to 12.3 Å) beyond which the contact is set to 0. In rMD, the protein evolves according to plain-MD as long as the reaction spontaneously proceeds towards the native state (i.e. lowering the value of the *z(x)* coordinate). On the other hand, when the chain tries to backtrack along *z(X)*, an external biasing force is introduced that redirects the dynamics towards the native state. The biasing force acting on a given atom, **F**_i_^rMD^, is defined as:

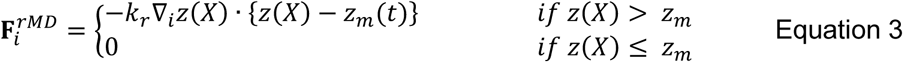

Where *z*_*m*_*(t)* indicates the smallest value of the reaction coordinate *z(X)* up to time *t* and *k*_*r*_ is a coupling constant.

#### Selection of the Least Biased Trajectory

For each set of trajectories starting from the same initial condition, the folding pathway with the highest probability to realize in the absence of external biasing force is selected. This scheme is applied by first defining a folding threshold: a trajectory is considered to have reached the folded state if its root mean squared deviation of atomic positions (RMSD) compared to the native target structure is ≤ 4 Å. Then, the trajectories successfully reaching the native state are scored by their computed bias functional *T*, defined as:

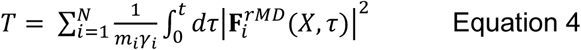

Where *t* is the trajectory folding time, *m*_*i*_ and *γ*_*i*_ are the mass and the friction coefficient of the *i*_*th*_ atom and **F**_i_^rMD^ is the force acting on it. The folding trajectory minimizing the bias functional for each set is referred to as Least Biased (LB) trajectory.

### Computational Analyses of the Androgen Receptor

#### Structure and Topology of AR Ligand Binding Domain

Atomistic structure of the AR ligand binding domain was retrieved from PDB 3B68, the structure spans from residue 671 to 917 and contains the receptor bound to SARM-S4 (an AR agonist). Ligand structure was retained in the plain-MD simulations, while it was removed to perform folding pathway simulations. Protein topology was generated in Gromacs 4.6.5 using Amber99SB-ILDN force field in TIP3P water. Ligand topology was generated by employing the AM1-BCC method in Antechamber.

#### Plain MD Simulations of AR Ligand Binding Domain

Setup and MD production was performed in Gromacs. The structure of AR ligand binding domain in complex with SARM-S4 was positioned in a dodecahedral box with 12 Å minimum distance from the walls. The box was filled with TIP3P water molecules and neutralized with 5 Cl^-^ ions. The system was energy minimized using the steepest descent algorithm. NVT equilibration was then performed for 500 ps at 310 K using the V-rescale thermostat (_T_T = 0.1 ps) with positional restraints on heavy atoms. The system was then equilibrated in NPT for 500 ps at 310 K and 1 Bar by employing the V-rescale thermostat and the Parrinello Rahman barostat (τP = 2 ps) with positional restraints on heavy atoms. Finally, 500 ns of plain MD were performed in the NPT ensemble by releasing the positional restraints. NVT, NPT and MD simulations were performed by using a leap-frog integrator with time-step equal to 2 fs, bond constraints were treated using LINCS. Van-der Waals and Coulomb cutoff was set to 12 Å while long-range electrostatics was treated using Particle Mesh Ewald. Atomic configurations were saved every 200 ps.

#### Analysis of Plain MD Simulation

RMSD of the MD trajectory was computed using Gromacs, while the solvent accessible surface area (SASA) of serine 791 was computed by employing the Shrake-Rupley algorithm in the python library MDTraj. The relative surface area (RSA), was calculated by normalizing the SASA of S791 with the SASA value of the serine in a GSG linear tripeptide.

#### Folding Simulations of the AR Ligand Binding Domain

The native structure of the AR ligand binding domain (PDB 3B68) was deprived of the ligand and positioned in a dodecahedral box with 40 Å minimum distance from the walls. The box was filled with TIP3P water molecules and neutralized with 5 Cl^-^ ions. The system was energy minimized using the steepest descent algorithm. NVT equilibration was then performed for 500 ps at 800 K using the V-rescale thermostat with positional restraints on heavy atoms. Restraints were then removed and three independent 5 ns of plain MD were performed in the NVT ensemble at 800 K, yielding three independent denatured conformation. Each initial condition was then repositioned in a dodecahedral box with 15 Å minimum distance from the walls, energy minimized using the steepest descent algorithm and then equilibrated first in the NVT ensemble (using the V-rescale thermostat at 350 K, τT = 0.1 ps) and then in the NPT ensemble (using the V-rescale thermostat at 350 K, τT = 0.1 ps, and the Parrinello-Rahman barostat at 1 bar, τP = 2 ps). For each initial condition 20 trajectories were generated by employing the rMD algorithm in the NPT ensemble (350 K, 1 bar). Each trajectory consists in 2.5 · 10^6^ rMD steps with integration time of 2 fs. Frames were saved every 5· 10^3^ steps. The ratchet constant *k*_*r*_ was set to 5 · 10^−4^ kJ/mol. Non-bonded interactions were treaded as follow: Van-der-Waals and Coulomb cutoff was set to 12 Å, while Particle Mesh Ewald was employed for long-range electrostatics. For each set, the bias functional scheme was applied yielding a total of 3 LB trajectories.

#### Analysis of Trajectories

RMSD was computed using Gromacs while the fraction of native contacts (Q) was computed using VMD 1.9.2. A lower-bound approximation of the energy landscape *G(Q, RMSD)* was generated by plotting the negative logarithm of the 2D probability distribution of the collective variables Q and RMSD, obtained from the 60 rMD trajectories (115×115 bins). Protein conformations belonging to the LB trajectories and spanning over the energetic wells of interest (*G* < 2.5 k_B_T) were sampled. The RSA was calculated as in the AR plain MD analysis employing the Shrake-Rupley algorithm in the python library MDTraj. Data were represented using the Matplotlib library in python, the 2D [Q, RMSD] energy plot was smoothed with a Gaussian kernel while the 2D RSA plot of serine 791 was interpolated using an inverse distance weighting with a quartic Lorentzian metrics. Images of the protein conformations were produced using UCSF Chimera.

### Computational Analyses of the Cellular Prion Protein

#### Structure and Topology of the C-terminal Domain of the Cellular Prion Protein (PrP)

The native structure of the C-terminal domain of human PrP was retrieved from PDB 1QLX, the structure spans from residue 125 to 228 and contain the structured globular domain of PrP. Protein topology was generated in Gromacs 4.6.5 using Amber99SB-ILDN force field in TIP3P water.

#### Folding Simulations of PrP

The native structure of the C-terminal domain of human PrP (PDB 1QLX) was positioned in a dodecahedral box with 40 Å minimum distance from the walls. The box was filled with spc216 water molecules and neutralized with 3 Na^+^ ions. The system was energy minimized using the steepest descent algorithm. NVT equilibration was then performed for 500 ps at 800 K using the V-rescale thermostat with positional restraints on heavy atoms. Restraints were then removed and 9 independent 3 ns of plain MD were performed in the NVT ensemble at 800 K, yielding 9 denatured conformation. Each initial condition was repositioned in a dodecahedral box with 15 Å minimum distance from the walls, energy minimized using the steepest descent algorithm and then equilibrated first in the NVT ensemble (using the Nosé-Hoover thermostat at 350 K, τT = 1 ps) and then in the NPT ensemble (using the Nosé-Hoover thermostat at 350 K, τT = 1 ps, and the Parrinello-Rahman barostat at 1 bar, τP = 2 ps). For each initial condition 20 trajectories were generated by employing the rMD algorithm in the NPT ensemble (350 K, 1 bar). Each trajectory consists in 1.5 · 10^6^ rMD steps generated with a leap-frog integrator with time-step of 2 fs. Frames were saved every 5· 10^2^ steps. The ratchet constant *k*_*r*_ was set to 5 · 10^−4^ kJ/mol. Non-bonded interactions were treaded as follow: Van-der-Waals and Coulomb cutoff was set to 16 Å, while Particle Mesh Ewald was employed for long-range electrostatics. For each set of trajectories, the bias functional scheme was applied with an additional filtering on the secondary structure content for folding definition. In particular, trajectories reaching a final conformation with less than 85% of average secondary structure content compared to the NMR structure were not considered in the ranking.

#### Analysis of Trajectories

RMSD was computed using Gromacs while the fraction of native contacts (Q) was computed using VMD 1.9.2. A lower-bound approximation of the energy landscape *G(Q, RMSD)* was generated by plotting the negative logarithm of the 2D probability distribution of the collective variables Q and RMSD, obtained from the 180 rMD trajectories (115×115 bins). Protein conformations belonging to the LB trajectories and spanning over the energetic wells of interest (G < 3.7 k_B_T) were sampled. Conformations belonging to the intermediate state were clustered by using a k-mean clustering in R-Studio ^88^ employing the following metrics for defining a distance between two structures:

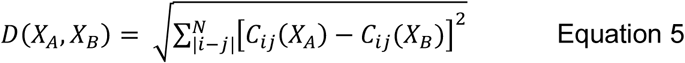

Where *D(X*_*A*_, *X*_*B*_) is the distance metrics between two protein conformations, *C*_*ij*_*(X*_*A*_) and *C*_*ij*_*(X*_*B*_) are the contact map entries of the conformations *A* and *B* respectively (defined in equation 2). The appropriate number of cluster (*k* = 3) was selected using the elbow method. The representative configuration of each cluster was selected by calculating the average contact map of the cluster conformations and then extracting the structure minimizing the distance *D(X*_*A*_, *X*_*B*_) between itself and the average contact map. Data were represented using the Matplotlib library in python, the 2D [Q, RMSD] energy plot was smoothed with a Gaussian kernel. Images of the protein conformations were produced using UCSF Chimera.

### Computer-Aided Drug Discovery Analyses

#### Consensus Approach for Druggable Ligand Binding Site Identification

A consensus approach relying on SiteMap (Schrödinger Release 2017-4: SiteMap, Schrödinger) ^52^ and DoGSiteScorer ^53^ analysis has been applied to find and evaluate druggable binding pocket. Default parameters were used for both tools. Specific structural properties, namely volume, depth, enclosure/exposure and balance and different druggability scores were computed for each identified site. In particular, the exposure, enclosure and depth properties provide a different measure of how the site is prone to be a deep pocket. For the exposure property, the lower the score, the better the site, with the average value for tight-binding sites found to be 0.49. Conversely, higher scores are preferred for the enclosure descriptor, with the average enclosure score for a tight-binding site being 0.78. On the other hand, the balance property of SiteMap expresses the ratio between the relative hydrophobic and hydrophilic character of a pocket. This fraction has proven to be a highly discriminating property in distinguishing between druggable and undruggable pockets, with a mean value for tight-binding site equal to 1.6. Besides the global pocket descriptors, the applied tools provide automated methods for a quantitative estimation of druggability. SiteMap predicts a site score (SiteScore) and druggabilty score (DScore) through a linear combination of only three single descriptors: the size of the binding pocket, its enclosure, and a penality for its hydrophilicity. The two scores differ in the coefficients, which are based on different training sets and strategies. A score value of ≥ 0.8 and ≥0.98 for SiteScore and DScore, respectively, has been reported to accurately distinguish between drug-binding and non-drug-binding sites. DoGSiteScorer also generates two scores, i.e Simple Score (SimpleScore) and a druggability score (DrugScore) which range from zero to one. The druggability cut-off for both scores is set to 0.5, indicating that targets with score above this value are considered as being druggable. Against this backdrop and considering the innovative character of the target herein reported (i.e. a folding intermediate structure), less stringent thresholds were thus selected in the search of druggable pockets to be explored in virtual screening: volume ≥ 300 Å3; depth ≥ 10 Å; balance≥1.0; exposure ≤ 0.5; enclosure ≥ 0.70; SiteScore ≥ 0.8; Dscore ≥ 0.90; DrugScore ≥ 0.5; SimpleScore ≥ 0.5 (Supp. Table 2 and 3).

#### Identification of a Druggable Pocket in the State I_1-3_ of AR

The N- and C-terminal residues of the intermediate folding state I_1-3_ of AR were capped with ACE and NMA groups, respectively. The obtained structure was submitted to the consensus approach in the search of a druggable ligand-binding site, leading to the collection of a set of global and druggability descriptors (Supp. Table 2).

#### Identification of a Druggable Pocket in the PrP Folding Intermediate

The previously described consensus approach allowed the identification of a possible ligand-binding region in the PrP folding intermediate, placed between the beginning of helix α1 and the loop that connects the α2 and α3 helices (Supp. Table 2). MD simulation were carried out to extract the most druggable conformation of this region, suitable for our virtual screening purpose. The folding intermediate was prepared with the Schroedinger’s Protein Preparation Wizard. During the preparation, the hydrogen bonding networks were optimized through an exhaustive sampling of hydroxyl and thiol groups. The N- and C-terminal residues were capped with ACE and NMA groups, respectively. Then, hydrogen atoms and protein side chains were energy-minimized using the OPLS3 force field. The obtained structure was (i) solvated by TIP3P water molecules in a cubic simulation box of 12.5 Å distant from the protein in every direction, (ii) neutralized by addition of three Na+ ions, and (iii) equilibrated for 100 ps of MD simulation (NPT ensemble) at 300 K using the Langevin thermostat. In such a simulation, the relative positions of the Cα atoms were kept fixed (force constant 1 Kcal/mol), in order to exclusively sample the arrangement of the side chains. Short-range electrostatic interactions were cut off at 9 Å, a RESPA integrator was used with a time step of 2 fs, and long-range electrostatics were computed every 6 fs. MD simulations were performed using the OPLS3 force field in Desmond 5.0 software (Schrödinger Release 2017-4: Desmond Molecular Dynamics System, D. E. Shaw Research, New York, NY, 2017) and run for 50ns. Recording interval was set to 50ps, allowing the collection of 1001 frames. The trajectory was clustered using the “Desmond trajectory clustering” tool in Maestro (Maestro-Desmond Interoperability Tools, Schrödinger) based on the RMSD of residues 152, 153, 156, 157, 158, 187, 196, 197, 198, 202, 203, 205, 206 and 209 (i.e. the residues composing the interested site). A hierarchical clustering was performed to obtain 10 clusters of the explored site. The centroid of each cluster was then selected as representative structure and subjected to in silico ligand binding site prediction and druggability assessment by using the above-mentioned consensus methods involving DogSiteScorer and SiteMap analysis (Supp Table 3).

#### Virtual Chemical Library Preparation

The Asinex Gold & Platinum Library was downloaded from the Asinex webpage (∼3.2 × 10^5^ commercially available compounds, www.asinex.com). A first round of ligand preparation was performed in LigPrep (Schrödinger Release 2017-4: LigPrep, Schrödinger). In this step, the different tautomeric forms for undefined chiral centers were created. By contrast, for specified chirality, only the specified enantiomer was retained. Subsequently, the compounds were imported within SeeSAR (SeeSAR Version 5.6, BioSolveIT GmbH), that assigned the proper geometry, the protonation state and the tautomeric form of the compounds using the ProToss method ^89^. A final library of ∼4.3 × 10^5^ docking clients was thus obtained.

#### Identification of In Silico Hits Through Virtual Screening

The virtual screening workflow was developed by using the KNIME analytic platform and the BioSolveIT KNIME nodes. Specifically, the workflow was organized as follows: (i) the “Prepare Receptor with LeadIT” node was used for protein preparation and docking parameters definition in LeadIT (LeadIT version 2.2.0; BioSolveIT GmbH, www.biosolveit.de/LeadIT). The binding site was defined on the basis of the residues composing the identified druggable pocket (Supp Fiure 2C). The residue protonation states, as well as the tautomeric forms, were automatically assessed in LeadIT using the ProToss method, that generates the most probable hydrogen positions on the basis of an optimal hydrogen bonding network using an empirical scoring function; (ii) the “Compute LeadIT Docking” node was selected to perform the docking simulations of the ∼4.3 × 105 docking clients by using the FlexX algorithm ^90^. Ten poses for each ligand were produced; (iii) the “Assess Affinity with HYDE in SeeSAR” node generated refined binding free energy (i.e. ΔG) and estimated HYDE affinity (K_iHYDE_) for each ligand pose using the HYDE rescoring function ^91^; (iv) for each ligand, the pose with the lowest K_iHYDE_ was extracted. Only compounds with a predicted K_iHYDE_ range below 5uM were retained for the following steps; (v) the rescored poses were filtered based on physicochemical and ADME filters using the Optibrium models integrated in SeeSAR (Optibrium 2018, www.optibrium.com/stardrop). In particular the following filters were used: 2 ≤ LogP ≤ 5, where LogP is the calculated octanol/water partition coefficient; 1.7 ≤ LogD ≤ 5, where logD is the calculated octanol/water distribution coefficient; TPSA ≤ 90; where TPSA is the topological polar surface area; LogS ≥ 1, where a LogS corresponds to intrinsic aqueous solubility greater than 10 µM; LogS7.4 ≥ 1, where LogS7.4 is the intrinsic aqueous solubility at pH of 7.4; HIA = +, where HIA is the classification for human intestinal absorption (predicts a classification of ‘+’ for compounds which are ≥30% absorbed and ‘-’ for compounds which are < 30% absorbed); 300 ≤ MW ≤ 500, where MW is molecular weight; number of rotable bonds ≤ 3; PgP category = -, where PgP category is the classification of P-glycoprotein transport (the compound must belong to the ‘-’ category to avoid the active efflux); number of hydrogen bond donors ≤ 3; number of stereocenters ≤ 1. In addition, molecules potentially acting as pan-assay interference compounds were discharged. This approach produced a list of 275 virtual hits, that were first submitted to a diversity-based selection. For each compound, a binary fingerprint was derived by means of the canvasFPgen utility provided by Schrödinger (Fingerprint type: MolPrint2D; precision:XP) (Schrödinger Release 2017-4: Canvas, Schrödinger). Using the created fingerprint, the 10 most different compounds (i.e. ASN 03578729, ASN 15755504, ASN 16356773, ASN 17325626, ASN 19380113, BAS 00312802, BAS 00340795, BAS 00382671, BAS 01058340, BAS 01849776) were extrapolated by applying the canvasLibOpt Schrodinger utility (Schrödinger Release 2017-4: Canvas, Schrödinger). In addition, visual inspection guided the selection of promising ligands based on the predicted binding mode and the interactions established with the identified binding pocket. In total, 30 molecules were selected, 8 from the diverse selection and 22 after visual inspection (Supp Figure 6 and Supp Table 4). Indeed, even though 10 diverse compounds were originally chosen, BAS 00340795 was not in stock and ASN 03578729 was later replaced by its close analogue ASN 05397475, selected by 3D visualization and endowed with a better predicted affinity.

### Chemical Synthesis of SM875

#### Reagents and Instrumentation

The reagents (Sigma Aldrich) and solvents (Merck) were used without purification. The reaction yields were not optimized and calculated after chromatographic purification. Thin layer chromatography (TLC) was carried out on Merck Kieselgel 60 PF254 with visualization by UV light. Microwave-assisted reactions were carried out using a mono-mode CEM Discover reactor in a sealed vessel. Preparative thin layer chromatography (PLC) on 20 × 20 cm Merck Kieselgel 60 F254 0.5 mm plates. HPLC purification was performed by a Merck Hitachi L-6200 apparatus, equipped with a diode array detector Jasco UVIDEC 100V and a LiChrospher reversed phase RP18 column, in isocratic conditions with eluent acetonitrile/water 1:1, flow 5 mL·min^-1^, detection at 254 nm. IR spectrum of the final product was recorded by using a FT-IR Tensor 27 Bruker spectrometer equipped with Attenuated Transmitter Reflection (ATR) device at 1 cm^-1^ resolution in the absorption region ΔV 4000 ÷ 1000 cm^-1^. A thin solid layer was obtained by the evaporation of the chloroform solution in the sample. The instrument was purged with a constant dry nitrogen flow. Spectra processing was made using Opus software packaging. NMR spectra were recorded on a Bruker-Avance 400 spectrometer by using a 5 mm BBI probe ^1^H at 400 MHz and ^13^C at 100 MHz in CDCl_3_ relative to the solvent residual signals δ_H_ 7.25 and δ_C_ 77.00 ppm, J values in Hz. Structural assignments are confirmed by heteronuclear multiple bond correlation (HMBC) experiment. ESI-MS spectra were taken with a Bruker Esquire-LC mass spectrometer equipped with an electrospray ion source, by injecting the samples into the source from a methanol solution MS conditions: source temperature 300 °C, nebulizing gas N_2_, 4 L·min^-1^, cone voltage 32 V, scan range m/z 100–900. LC-ESI-MS spectrum was acquired using a C-18 Kinetex 5 µm column, eluting with acetonitrile/ water 70:30, flow 1 mL·min^-1^ using ESI source as detector in positive ion mode. High-resolution ESI-MS measurement for the final product, including tandem MS^2^ fragmentation experiments, were obtained by direct infusion using an Orbitrap Fusion Tribrid mass spectrometer.

#### Synthesis of SM875

The target product SM875 was obtained according the synthetic strategy reported in Figure 7. The sequence involves the preparation of the precursor 1-(4-bromophenyl)-1H-pyrazol-5-amine [1], which was used in a following three-component reaction with 4-hydroxy-3-methoxybenzaldehyde and Meldrum acid (2,2-dimethyl-1,3-dioxane-4,6-dione) according to a modified method ^92^. The 1-(4-bromophenyl)-1H-pyrazol-5-amine was synthesized starting from 4-(bromophenyl) hydrazine that was obtained from the commercial hydrochloride by treatment with a saturated NaHCO_3_ aqueous solution (50 mL), followed by dichloromethane extraction (50 mL x3), treatment with anhydrous Na_2_SO_4_ and evaporation. To a magnetically stirred solution of 4-(bromophenyl)hydrazine (100 mg, 0.53 mM, in 5 mL ethanol 5), (ethyl 2-cyano-3-ethoxyacrylate (89.6 mg, 0.53 mM) was added and refluxed for 2 h. The reaction mixture was concentrated *in vacuo*, the residue was suspended in 1:1 methanol/ 2M NaOH aq. solution (5 mL) and refluxed for 1 h. After cooling, the mixture was neutralized with 1M HCl aq. solution (5 mL) and concentrated *in vacuo* using a water bath at 40°C. The residue was heated at 180°C for 10 min, suspended in ethanol after cooling and stored overnight at 4°C. The supernatant was recovered and concentrated to give a residue which was stirred in the presence of a NaHCO_3_ solution (10 mL). Extraction with ethyl acetate (10 mL x3), followed by the treatment with anhydrous Na_2_SO_4_ of the combined organic phases and concentration *in vacuo* gave the product (79 mg, 61%), which was used in the following three component reaction. The successful synthesis of 1-(4-bromophenyl)-1H-pyrazol-5-amine was verified by ^1^H-NMR [δ_H_ 7.58(d, J 8.7 Hz, H-3’and H-5’), 7.47(d, J 8.7 Hz, H-2’and H-6’), 7.41(s, H-3), 5.62 (s, H-4)] and ESIMS (m/z 239.8 [M+H]^+^). In the last reaction step, 1-(4-bromo-phenyl)-1H-pyrazol-5-amine (79 mg, 0.33 mM), 4-hydroxy-3-methoxybenzaldehyde (41 mg, 0.27 mM) and 2,2-dimethyl-1,3-dioxane-4,6-dione (46 mg, 0.32 mM) in ethanol (5 mL) were refluxed under stirring for 2.5 h, or alternatively by replacing the conventional heating with microwave irradiation at 110°C for 1 h. The reaction mixture was then cooled to room temperature (RT) and dried in vacuo. The raw product was purified by silica preparative thin layer chromatography (PLC) eluting with n-hexane / ethyl acetate (1:1). The band collected at retention factor 0.4 was first used for structural characterization and then injected into preparative HPLC (RP-18 column, acetonitrile/ water 1:1, UV detection at 254 nm, flow 5 mL·min^-1^, retention time 4.5 min) to give the target product (for use on cell cultures) as a white powder after evaporation of the eluent: 34 mg, 25% with reflux in ethanol; a yield of ∼25% is also obtained with microwave irradiation.

#### Structural Characterization of SM875

NMR, HRESI-MS spectra together with LC-MS (UV, EIC and MS) are reported in Figure 8 and Supp. Table 5. Figure 8B, relative to HRESI-MS, reports the results of single measurements, while the average of 80 measures in positive ion mode is reported here: HRESI(+)MS: m/z 414.0443 ± 0.0013, [M+H]^+^ (calculated for C_19_H_17_^79^BrN_3_O_3_ = 414.0448); HRESI(+)MS/MS on the fragment ions of 414.0443 :m/z 399.0205 ± 0.0019, [M+H-CH_3_]^+^ (calculated for C_18_H_14_^79^BrN_3_O_3_ = 399.0213); m/z 372.0335 ± 0.0019, [M+H-C_2_H_2_O]^+^ (calculated for C_17_H_15_^79^BrN_3_O_2_ = 372.0342); m/z 289.9918 ± 0.0017, [M+H-C_7_H_8_O_2_]^+^ (calculated for C_12_H_9_^79^BrN_3_O = 372.0342).

### Cell cultures and treatments

Cell lines used in this paper have been cultured in Dulbecco’s Minimal Essential Medium (DMEM, Gibco, #11960-044), 10% heat-inactivated fetal bovine serum (Δ56-FBS), Penicillin/Streptomycin (Pen/Strep, Corning #20-002-Cl), non-essential amino acids (NEAA, Gibco, #11140-035) and L-Glutamine (Gibco, #25030-024), unless specified differently. HEK293 and N2a cells were obtained from ATCC (ATCC CRL-1573 and CCL-131, respectively). We used a subclone (A23) of HEK293 stably expressing a mouse wild-type PrP or an EGFP-PrP construct, both already described and characterized previously ^62^. L929 mouse fibroblasts and inducible RK13 cells were kindly provided by Ina Vorberg (DZNE, Bonn, Germany) ^76^ and Didier Villette (INRA, Toulouse, France) ^72^, respectively. Human cancer cell lines (H358, ZR-75, A549, H460, MCF7, H1299, SKBR3 and T47D), all belonging to the NCI collection of human cancer cell lines, were kindly provided Valentina Bonetto (Mario Negri Institute, Milan, Italy). Cells were passaged in T25 flasks or 100 mm Petri dishes in media containing 200 µg/ml of Hygromycin or 500 µg/ml of G418 and split every 3-4 days. Every cell line employed in this study has not been passaged more than 20 times from the original stock. Compounds used in the experiments were resuspended at 30 or 50 mM in DMSO, and diluted to make a 1000X stock solution, which was then used for serial dilutions. A 1 µl aliquot of each compound dilution point was then added to cells plated in 1 mL of media with no selection antibiotics.

For pulse experiments, inducible RK13 cells were seeded on 24-well plates at a confluence of 50%. After 24 h cells were treated with doxyciclin (0.01 mg/ml) or vehicle (DMSO), in the presence or absence of brefeldin 1A (BREF 10 uM) or SM875 (10 uM). At the end of each time-point (2, 4, 8 and 24 h) cells were washed with PBS and then lysed in lysis buffer. For chase experiments, RK13 cells were seeded on 24-well plates at a confluence of 30%. After 24 h cells were treated with Doxyciclin (0.01 mg/ml) for 24 h. The medium containing doxyciclin was then removed and cells kept in fresh medium for 4 h before adding SM875 (10 uM). After 5, 19 and 24 h of incubation cells wells were washed with PBS and lysed.

### Plasmids

Cloning strategies used to generate cDNAs encoding for WT or EGFP-tagged PrP have been described previously ^93^. The EGFP-PrP construct contains a monomerized version of EGFP inserted after codon 34 of mouse PrP. The identity of all constructs was confirmed by sequencing the entire coding region. All constructs were cloned into the pcDNA3.1(+)/hygro expression plasmid (Invitrogen). The Strep-FLAG pcDNA3.1(+)/G418 NEGR-1 and the anti-NEGR-1 primary antibody were kindly provided by Giovanni Piccoli (University of Trento, Italy) ^94^. All plasmids were transfected using Lipofectamine 2000 (Life Technologies), following manufacturer’s instructions.

### Western blotting & Antibodies

Samples were lysed in lysis buffer (Tris 10 mM, pH 7.4, 0.5% NP-40, 0.5% TX-100, 150 mM NaCl plus complete EDTA-free Protease Inhibitor Cocktail Tablets, Roche, #11697498001), diluted 2:1 in 4X Laemmli sample buffer (Bio-Rad) containing 100 mM Dithiothreitol (CAS No. 3483-12-3, SigmaAldrich), boiled 8 min at 95 °C and loaded on SDS-PAGE, using 12% acrylamide pre-cast gels (Bio-Rad) and then transferred to polyvinylidene fluoride (PVDF) membranes (Thermo Fisher Scientific). Membranes were blocked for 20 min in 5% (w/v) non-fat dry milk in Tris-buffered saline containing 0.01% Tween-20 (TBS-T). Blots were probed with anti-PrP antibodies D18 (kindly provided by D. Burton, The Scripps Research Institute, La Jolla, CA) or 6D11 (1:5,000) in BSA 3% in TBS-T overnight at 4 °C, anti-NEGR-1 antibody (1:5,000, kindly provided by Giovanni Piccoli, University of Trento, Italy), anti-Thy-1 (1:5,000, OX7, Abcam) or anti-LC3 (1:2,000, ab51520, Abcam), all an in 5% (w/v) non-fat dry milk. After incubation with primary antibodies, membranes were washed three times with TBS-T (10 min each), then probed with a 1:8,000 dilution of horseradish conjugated goat anti-human (Jackson Immunoresearch) or anti-mouse (Santa Cruz) IgG for 1 h at RT. After 2 washes with TBS-T and one with Milli-Q water, signals were revealed using the ECL Prime western blotting Detection Kit (GE Healthcare), and visualized with a ChemiDoc XRS Touch Imaging System (Bio-Rad). In all the experiments, with the exception of PK-treated samples, the final quantification of proteins detected by primary antibodies were obtained by densitometric analysis of the western blots, normalizing each signal on the corresponding total protein lane (obtained by the enhanced tryptophan fluorescence technology of stain-free gels, BioRad).

### Dot Blotting

Samples were spotted on PVDF membrane in a 96-well dot blot system. A 96-well dot blot apparatus (Schleicher & Schuell) was set up with a 0.45-μm-pore-size polyvinylidene difluoride (PVDF) membrane (Immobilon-P; Millipore), and each dot was rinsed with 500 μl of TBS. Under vacuum, cell lysates diluted 1:10 in TBS were added to the apparatus and rinsed with 500 μL of TBS. The membrane was then blocked with 5% milk-0.05% Tween 20 (Sigma) in TBS (TBST-milk) for 30 min and probed with the anti-PrP antibody 6D11 (1:4,000) followed by goat anti-mouse IgG (Pierce). Signals were revealed using enhanced chemiluminescence (Luminata, Bio-Rad) and visualized by a ChemiDoc XRS Touch Imaging System (Bio-Rad).

### Quantitative Real Time PCR

Following treatments, cells were harvested from 24 wells plates and RNA was extracted using TRIzol (Invitrogen) or RNeasy Plus mini kit (Quiagen). An 800-ng aliquot per each sample was reverse transcribed using High Capacity cDNA Reverse Transcription kit (Applied Biosystems) according to the manufacturer’s instructions. Quantitative RT-PCR was performed in a CFX96 Touch thermocycler (Bio-Rad) using PowerUp SYBR Green Master mix (Invitrogen) for 40 cycles amplification. Mouse PrP set 1 and set 2 were used to amplify endogenous and transgenic PrP, respectively (table below). Relative quantification was normalized to mouse or human HPRT (hypoxanthine-guanine phosphoribosyltransferase) as a housekeeping control.

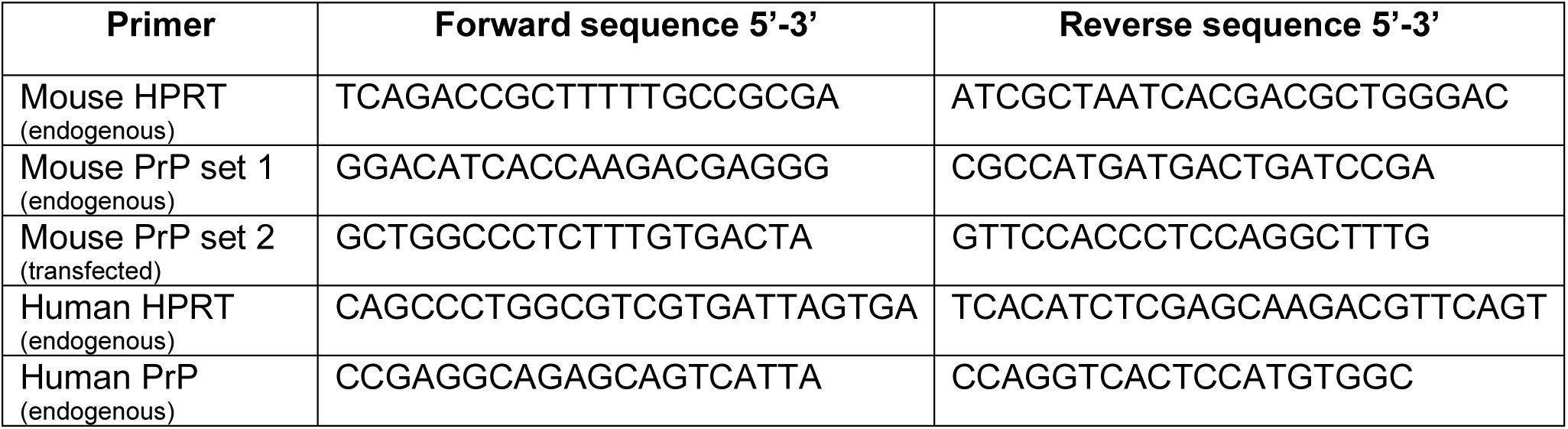

### Immunofluorescence and High Content Imaging

Immunocytochemistry was performed on inducible RK13 cells treated for 2, 4, 8, or 24 h with doxycycline 0.01 (1x) or 0.1 (10x) mg/mL, in the presence or absence of SM875 (10 µM). Cells were seeded on CellCarrier-384 Ultra microplates (Perkin Elmer) at a concentration of 6,000 cells/well, grown for approximately 24 h, to obtain a semi-confluent layer (60%) and then treated with the compound. Cells were fixed for 12 min at RT by adding methanol-free paraformaldehyde (PFA, Thermo Fisher Scientific) to a final concentration of 4%. Wells were then washed three times with PBS, and permeabilized for 1 min with PBS containing a final concentration of 0.1% Triton X-100. Wells were washed three times with PBS and cells were incubated with blocking solution (FBS 2% in PBS) for 1h at RT. The anti-PrP primary antibody (D18) was diluted in the blocking solution and added to the wells to a final concentration of 1:400. After three washes with PBS, the secondary antibody (Alexa 488-conjugated goat anti-human IgG diluted 1:500 in blocking solution) was incubated for 1 h at RT. Hoechst 33342 (Thermo Fisher Scientific) diluted in 0.5 mM PBS was then added after two additional washes. In another set of experiments, cells expressing EGFP-PrP were plated on CellCarrier-384 Ultra microplates (Perkin Elmer) at a concentration of 12,000 cells/well and grown for approximately 24 h, to obtain a semi-confluent layer (60%). SM875 was administered at a final concentration of 0.1, 0.3, 1, 3 or 10 µM, in two replicate wells. Vehicle (DMSO, volume equivalent) was used as a negative control. Cells were treated for 24 h and then fixed for 12 min at RT by adding methanol-free paraformaldehyde (Thermo Fisher Scientific) to a final concentration of 4%. Plates were then washed twice with PBS and counterstained with Hoechst 33342. The cell localization of EGFP-PrP and inducible PrP was monitored using an Operetta High-Content Imaging System (Perkin Elmer). Imaging was performed in a widefield mode using a 20X High NA objective (0.75). Five fields were acquired in each well over two channels (380-445 Excitation-Emission for Hoechst and 475-525 for EGFP and Alexa 488). Image analysis was performed using the Harmony software version 4.1 (Perkin Elmer).

### Cell viability

Cells were seeded on 24-well plates at approximately ∼60% confluence. Compounds at different concentrations or vehicle control (DMSO, volume equivalent) were added after 48 h, medium was replaced the second day, and then removed after a total of 48 h of treatment. Cells were incubated with 5 mg/mL of 3-(4,5-dimethylthiazol-2-yl)-2,5-diphenyltetrazolium bromide (MTT, Sigma Aldrich) in PBS for 15 min at 37 °C. After carefully removing MTT, cells were resuspended in 500 µL DMSO, and cell viability values obtained by a plate spectrophotometer (BioTek Instruments, VT, USA), measuring absorbance at 570 nm.

### Production of recombinant PrP

RecHuPrP23-231 was expressed by competent E. coli Rosetta (DE3) bacteria harboring pOPIN E expression vector containing the wild type I109 bank vole Prnp gene ^95^. Bacteria from a glycerolate maintained at -80 °C were grown in a 250 ml Erlenmeyer flask containing 50 ml of LB broth overnight. The culture was then transferred to two 2 L Erlenmeyer flasks containing each 500 ml of minimal medium supplemented with 3 g/L glucose, 1 g/L NH_4_Cl, 1M MgSO_4_, 0.1 M CaCl_2_, 10 mg/mL thiamine and 10 mg/mL biotin. When the culture reached an OD_600_ of ∼0.9-1.2 AU, Isopropyl β-D-1-thiogalactopyranoside (IPTG) was added to induce expression of PrP overnight under the same temperature and agitation conditions. Bacteria were then pelleted, lysed, inclusion bodies collected by centrifugation, and solubilized in 20 mM Tris-HCl, 0.5M NaCl, 6M Gnd/HCl, pH = 8. Although the protein does not contain a His-tag, purification of the protein was performed with a histidine affinity column (HisTrap FF crude 5 ml, GE Healthcare Amersham) taking advantage of the natural His present in the octapeptide repeat region of PrP. After elution with buffer containing 20 mM Tris-HCl, 0.5M NaCl, 500 mM imidazole and 2 M guanidine-HCl, pH = 8, the quality and purity of protein batches was assessed by BlueSafe (NZYTech, Lisbon) staining after electrophoresis in SDS-PAGE gels. The protein was folded to the PrP^C^ conformation by dialysis against 20 mM sodium acetate buffer, pH = 5. Aggregated material was removed by centrifugation. Correct folding was confirmed by CD and protein concentration, by measurement of absorbance at 280 nm. The protein was concentrated using Amicon centrifugal devices and the concentrated solution stored at -80 °C until used.

### Dynamic mass redistribution

The EnSight Multimode Plate Reader (Perkin Elmer) was used to carry out DMR analyses. Immobilization of full-length (residues 23-230) or mouse N-terminally truncated (111-230) recombinant PrP (15 µL/well of a 2.5 µM PrP solution in 10 mM sodium acetate buffer, pH 5) on label-free microplates (EnSpire-LFB high sensitivity microplates, Perkin Elmer) was obtained by amine-coupling chemistry. The interaction between Fe^3+^-TMPyP, SM875 and SM940 diluted to different concentrations (0.03-100 μM, eight 1:3 serial dilutions) in assay buffer (10 mM PO_4_, pH 7.5, 2.4 mM KCl, 138 mM NaCl, 0.05% Tween-20) and PrP, was monitored after a 30 min incubation at RT. All the steps were executed by employing a Zephyr Compact Liquid Handling Workstation (Perkin Elmer). Data were obtained by normalizing each signal on the intra-well empty surface, and then by subtraction of the control wells. The Kaleido software (Perkin Elmer) was employed to acquire and process the data.

### Temperature-dependent detergent insolubility assay

An 800 µM stock solution of freshly purified mouse recombinant PrP (residues 111-231) was diluted 1:10 in sodium acetate (10 mM NaAc, pH 7) to obtain 80 µM aliquots. To avoid precipitation of recombinant PrP, aliquots were flash frozen in liquid nitrogen, stored at -80 °C, and then kept on ice during their use. In order to carry out the assay, recombinant PrP was diluted to a final concentration of 0.5 µM in precipitation buffer (10 mM NaAc, 2% TX100, pH 7), split in 8 identical aliquots, and incubated for 1 h at different temperatures (25, 37, 45, and 55 ° C), in the presence or absence of each molecule, or vehicle control (DMSO, volume equivalent). Each sample was then carefully loaded onto a double layer of sucrose (60% and 80%) prepared in precipitation buffer and deposited at the bottom of ultracentrifuge tubes. Samples were then subjected to ultracentrifugation at 100,000 *x g* for 1 hour at 4 °C. The obtained protein pellets were diluted in 2X LMSB and then analyzed by western blotting.

### Detection of prions in L929 fibroblasts

L929 fibroblasts were grown in culturing medium and passaged 5-7 times after infection with a 0.5% homogenate of Rocky Mountain Laboratory (RML) prion strain (corresponding to ∼1 × 10^5^ lethal dose at 50%, derived from corresponding prion-infected mice; courtesy of Dr. Roberto Chiesa, Mario Negri Institute, Milan). In order to test the anti-prion effects of compounds, cells were seeded in 24-well plates (day 1) at approximately 60% confluence, with different concentrations of each molecule, or vehicle control (DMSO, volume equivalent). Medium containing fresh compounds or vehicle was replaced on day 2, and cells were split (1:2) on day 3, avoiding the use of trypsin by pipetting directly onto the well surface. Cells were collected on day 4 in PBS and centrifuged at 3.500 rpm x 3 min. The resulting pellets were then rapidly stored at -80 °C. To evaluate prion loads, cell pellets were resuspended in 20 µL of lysis buffer (Tris 10 mM, pH 7.4, 0.5% NP-40, 0.5% TX-100, 150 mM NaCl) and incubated for 10 min at 37 °C with 2.000 units/mL of DNase I (New England BioLabs). Half of the resulting sample was incubated with 10 µg/mL of PK (Sigma Aldrich) for 1h at 37°C, while the other half was incubated in the same conditions in the absence of PK. Both PK-treated and untreated samples were then mixed 1:2 with 4X Laemmli sample buffer (Bio-Rad) containing DTT, boiled for 8 min at 95°C and ran by SDS-PAGE. The final quantification of PK-resistant PrP species was obtained by densitometric analysis of the western blots, normalizing each signal on the corresponding PK-untreated lane.

### Crystallization

Recombinant mouse PrP (residues 111-230, 800 µM) was heated to 45 °C and slowly cooled to 20°C using a thermal cycler, either in the presence or absence of SM875 (2 mM). Massive precipitation was observed in the presence of SM875. Pellet was removed by centrifugation and a 40% reduction in PrP concentration was estimated by UV absorbance at 280 nm. Showers of very thin needles appeared in 2-4 days (0.2 M (NH_4_)_2_SO_4_, 10 mM CdCl_2_, 9-14% LMW PEG smear, 9-14% MMW PEG smear, pH 6-8) irrespectively of whether the protein was incubated with SM875 or not. Data collection was performed at the Elettra synchrotron, XRD1 beamline (Trieste, Italy). Crystals were weakly and anisotropically diffracting to 3.7 Å in the best directions. Multiple diffraction patterns were present, as single needles were impossible to isolate. Orthorhombic crystal system and unit cell dimensions could be identified (a = 36.1 Å, b = 51.8 Å, c = 55.9 Å), which are extremely similar to those reported for apo human PrP in PDB 3HAK: a = 32.5 Å, b = 49.1 Å, c = 56.9 Å.

### Statistical analyses of biological data

All the data were collected and analyzed blindly by two different operators. Statistical analyses, performed with the Prism software version 7.0 (GraphPad), included all the data points obtained, with the exception of experiments in which negative and/or positive controls did not give the expected outcome, which were discarded. No test for outliers was employed. The Kolmogorov-Smirnov normality test was applied (when possible, n≥5). Results were expressed as the mean ± standard errors, unless specified. In some case, the dose-response experiments were fitted with a 4-parameter logistic (4PL) non-linear regression model, and fitting was estimated by calculating the R^2^. All the data were analyzed with the one-way ANOVA test, including an assessment of the normality of data, and corrected by the Dunnet post-hoc test. Probability (*p)* values < 0.05 were considered as significant (*<0.05, **<0.01, ***<0.001).

## ACKNOWLEDGMENTS

We thank Ina Vorberg (DZNE, Bonn, Germany) for mouse L929 fibroblasts, Didier Villette (INRA, Toulouse, France) for RK13 cells, Roberto Chiesa (Mario Negri Institute in Milan, Italy) for RML brain homogenates and Dennis Burton (The Scripps Research Institute, La Jolla, CA) for D18-expressing CHO cells. The authors are grateful to the staff of the XRD1 beamline at Elettra (Trieste, Italy) for on-site assistance, and acknowledge a CINECA award under the ISCRA initiative, for the availability of high-performance computing resources and support. LT is supported by Fondazione Caritro (Bando Post Doc 2017) and Kennedy’s Disease Association (Research Grant 2018). Research of HCA was supported by the CJD Foundation, Inc. and the Alzheimer Forschung Initiative e.V. (AFI). JRR was funded by a grant (BFU2017-86692-P) from the Spanish Ministry of Economy and Competitiveness, partially funded by FEDER funds. The work was also supported by grants from Fondazione Telethon and Provincia Autonoma di Trento to SB (TCP13007), from Fondazione Telethon to EB (TCP14009), and by a fellowship from Fondazione Telethon to GS. SB and EB are Assistant Telethon Scientists at the Dulbecco Telethon Institute.

## AUTHOR CONTRIBUTIONS

Conceived and designed the computational analyses: GS, AA, AI, SO, AB, LT, LTo, MP, MLB, PF, EB; Conceived and designed the experimental analyses: GS, TM, SB,GP, GG, MC, HCA, GL, SB, IM, EB; Performed the computational analyses: GS, AA, AI, SO, AB, LT, MR, MLB, PF; Performed the experimental analyses: GS, TM, SB, PB, ML, GM, NLL, LCF, LL, BV, DG, GG, GL, MMP, IM; Analyzed the data: GS, TM, AA, SB, ML, AI, SO, AB, LT, GL, SB, JRR, IM, MLB, PF, EB; Wrote the paper: GS and EB. All authors edited the manuscript.

## FIGURE LEGENDS

**Supp. Figure 1.**
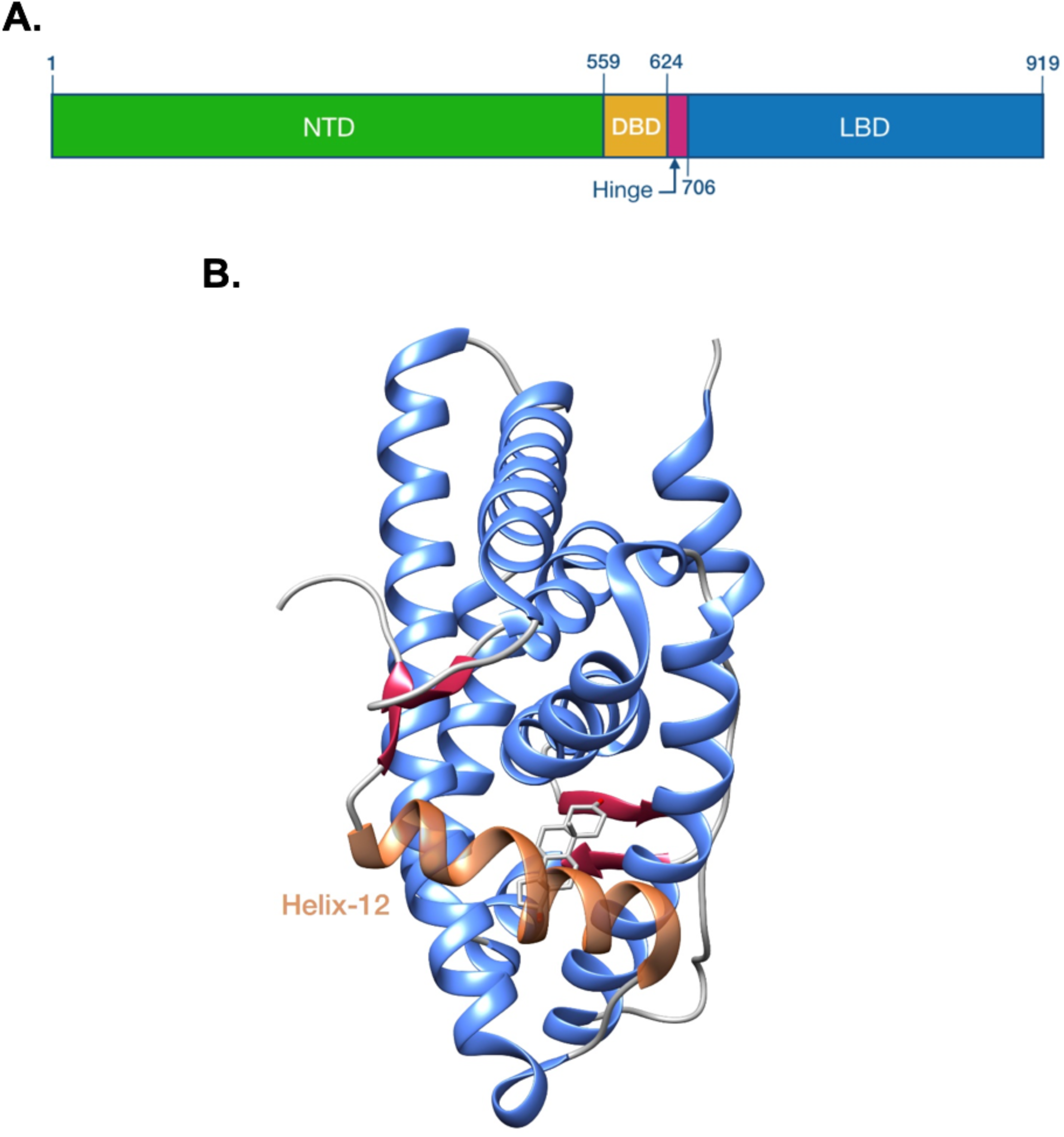
Structure of the AR. **A**. The picture illustrates a schematic representation of the structure of AR. The protein is organized in three domains: an intrinsically disordered N-Terminal Domain (NTD), responsible for the interaction with trans-activator proteins necessary for transcriptional activity; a DNA-Binding Domain (DBD); a C-terminal Ligand-Binding Domain (LBD) capable of binding androgens, which determine the activation of AR. **B**. Ribbon diagram of the native structure of the AR ligand-binding domain in complex with DHT (PDB 2AMA). The picture shows the topology of helix 12 (depicted in orange) covering the ligand-binding pocket.

**Supp. Table 1.** Solvent-accessible surface area (SASA) and relative surface area (RSA) for residue S791. The table illustrates SASA and RSA values for residue S791 of the AR ligand-binding domain from 75 different PDB entries. Based on a typical threshold to define a residue buried in the protein core of RSA < 20%, which corresponds to a serine SASA of 25 Å^2^, these data indicate that S791 is consistently excluded from the solvent in all the deposited structures (1.2 Å^2^ < SASA < 12 Å^2^; 1% < RSA < 9.5%).

| PDB ID | SASA [Å<sup>2</sup>] | RSA % |
| --- | --- | --- |
| 1E3G | 1.20 | 1.0 |
| 1GS4 | 2.90 | 2.3 |
| 1T5Z | 3.11 | 2.5 |
| 1T63 | 3.90 | 3.1 |
| 1T65 | 3.63 | 2.9 |
| 1XJ7 | 6.97 | 5.6 |
| 1XOW | 2.73 | 2.2 |
| 1XQ3 | 2.90 | 2.3 |
| 1Z95 | 2.91 | 2.3 |
| 2AM9 | 2.89 | 2.3 |
| 2AMA | 2.23 | 1.8 |
| 2AMB | 2.25 | 1.8 |
| 2AO6 | 2.87 | 2.3 |
| 2AX6 | 2.48 | 2.0 |
| 2AX7 | 2.31 | 1.8 |
| 2AX8 | 2.16 | 1.7 |
| 2AX9 | 2.00 | 1.6 |
| 2AXA | 1.88 | 1.5 |
| 2HVC | 3.65 | 2.9 |
| 2OZ7 | 3.54 | 2.8 |
| 2PIO | 3.16 | 2.5 |
| 2PIP | 3.95 | 3.2 |
| 2PIQ | 3.33 | 2.7 |
| 2PIR | 3.02 | 2.4 |
| 2PIT | 2.41 | 1.9 |
| 2PIU | 3.65 | 2.9 |
| 2PIV | 2.80 | 2.2 |
| 2PIW | 3.69 | 3.0 |
| 2PIX | 3.54 | 2.8 |
| 2PKL | 1.73 | 1.4 |
| 2PNU | 2.08 | 1.7 |
| 2Q7I | 2.09 | 1.7 |
| 2Q7J | 2.26 | 1.8 |
| 2Q7K | 2.08 | 1.7 |
| 2Q7L | 2.12 | 1.7 |
| 2YHD | 2.14 | 1.7 |
| 2YLO | 2.34 | 1.9 |
| 2YLP | 2.28 | 1.8 |
| 2YLQ | 3.07 | 2.5 |
| 2Z4J | 1.08 | 0.9 |
| 3B5R | 2.34 | 1.9 |
| 3B65 | 2.03 | 1.6 |
| 3B66 | 2.29 | 1.8 |
| 3B67 | 1.84 | 1.5 |
| 3B68 | 1.81 | 1.4 |
| 3L3X | 2.61 | 2.0 |
| 3L3Z | 2.65 | 2.1 |
| 3RLJ | 2.47 | 2.0 |
| 3RLL | 2.61 | 2.1 |
| 3V49 | 2.00 | 1.6 |
| 3V4A | 2.09 | 1.7 |
| 4HLW | 2.29 | 1.8 |
| 4K7A | 3.64 | 2.9 |
| 4OEA | 1.94 | 1.5 |
| 4OED | 3.40 | 2.7 |
| 4OGH | 1.65 | 1.3 |
| 4OH5 | 2.44 | 1.9 |
| 4OH6 | 2.95 | 2.4 |
| 4OHA | 2.10 | 1.7 |
| 4OIL | 2.89 | 2.3 |
| 4OIU | 3.21 | 2.6 |
| 4OJ9 | 2.04 | 1.6 |
| 4OJB | 2.35 | 1.9 |
| 4OK1 | 2.72 | 2.2 |
| 4OKB | 4.08 | 3.3 |
| 4OKT | 2.72 | 2.2 |
| 4OKW | 1.93 | 1.5 |
| 4OKX | 1.92 | 1.5 |
| 4OKX | 2.32 | 1.9 |
| 4QL8 | 2.50 | 2.0 |
| 5JJM | 11.76 | 9.4 |
| 5T8E | 5.05 | 4.0 |
| 5T8J | 3.33 | 2.7 |
| 5V8Q | 2.40 | 1.9 |
| 5VO4 | 2.10 | 1.7 |

**Supp. Figure 2.**
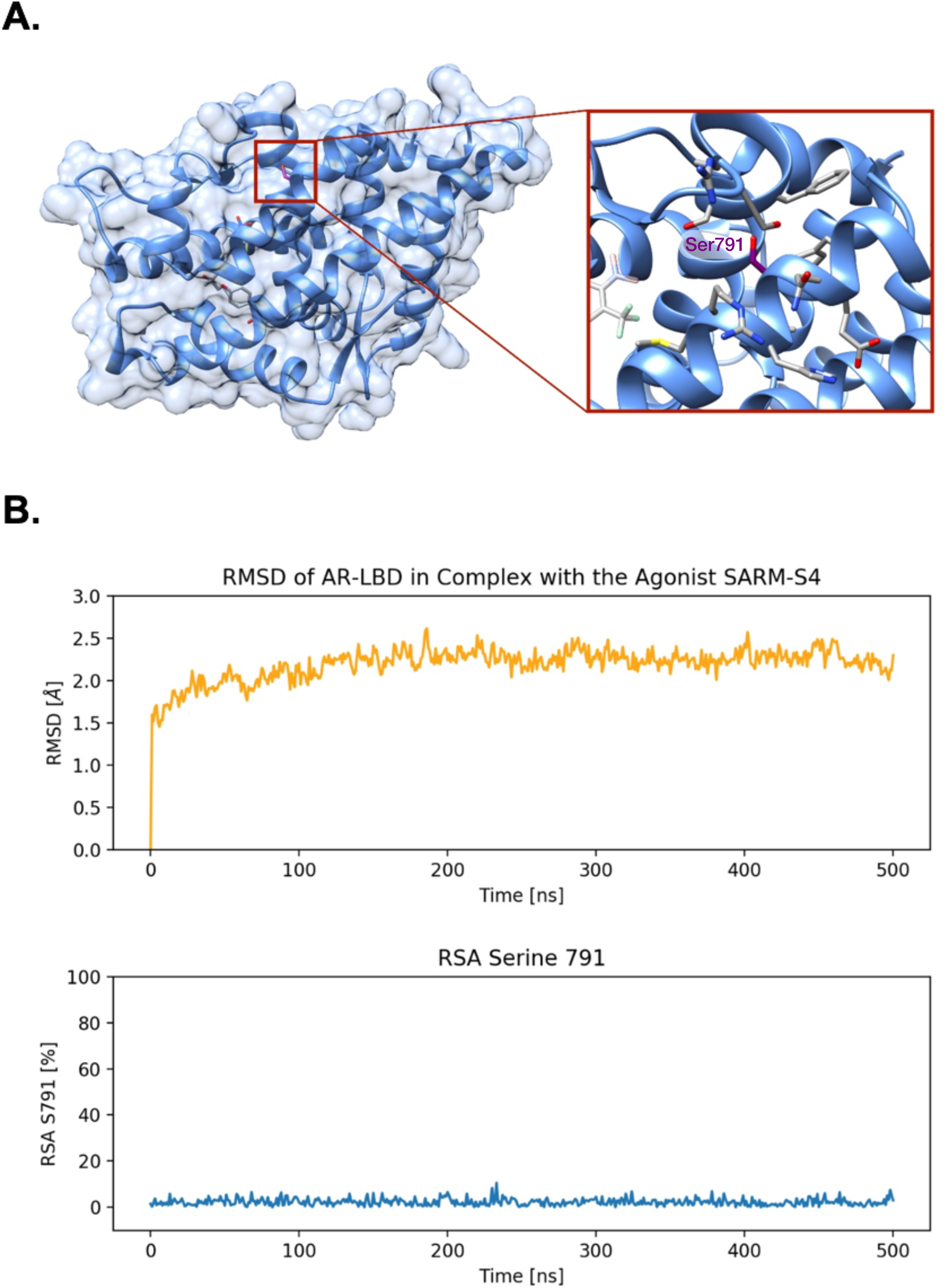
MD simulations of AR ligand-binding domain. **A**. Ribbon diagram of AR ligand-binding domain in complex with agonist SARM-S4 (PDB 3B68). The box shows the position of S791 buried inside the AR structure. **B**. All-atom MD simulations (500 ns) from initial configuration to RMSD convergence of the entire AR ligand-binding domain in complex with SARM-S4 (upper panel). The analysis shows that S791 is kept inside the protein core for the entire length of the simulation (as expressed as % RSA in the lower panel).

**Supp. Figure 3.**
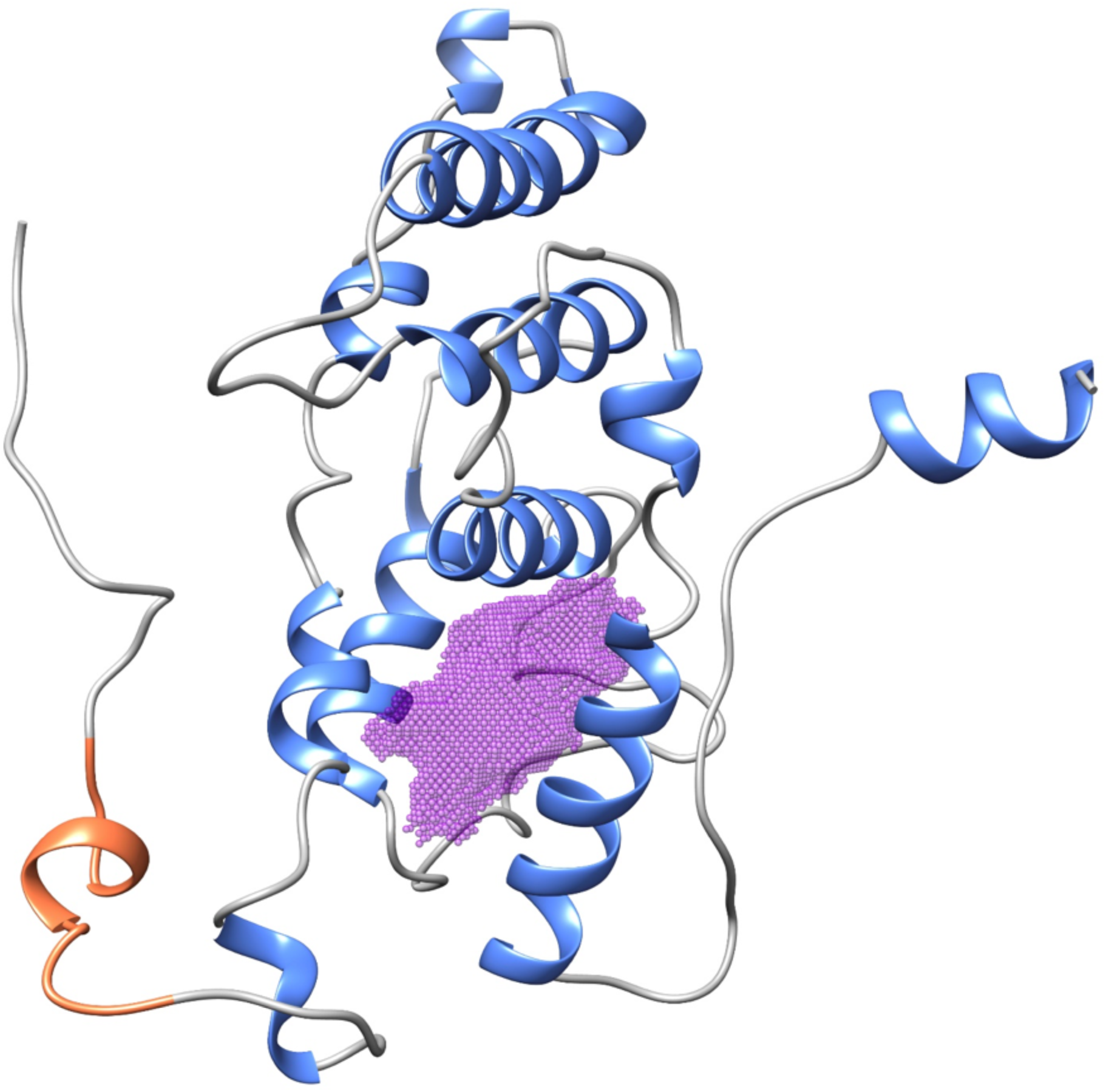
Druggable pocket in the AR intermediate state (I_1_). Druggable pocket in the AR intermediate state (I_1_): Ribbon diagram of the AR intermediate I_1_ with highlighted the ligand-binding pocket (purple dots) as identified by SiteMap and DogSiteScorer tools. The pocket, surrounded by helices 3 and 5, is accessible before the closing of the helix 12 (depicted in orange).

**Supp. Table 2.** Druggability descriptors of the pocket in the AR intermediate state (I_1_). The table reports the computed structural and druggability descriptors for the identified ligand-binding pocket in the I_1_ conformation of the AR. The reported values were calculated by SiteMap and DogSiteScorer algorithms. SiteScore, DScore, SimpleScore, and DrugScore are different scoring metrics for druggability assessments, whereas the exposure, enclosure and depth properties provide a different measure of how the site is prone to be a deep pocket. The balance property of the SiteMap tool expresses the ratio between the relative hydrophobic and hydrophilic characters of the site and is a key property in distinguishing between druggable and undruggable pockets.

| Descriptor | Threshold | AR I<sub>1</sub> Pocket |
| --- | --- | --- |
| SiteScore<sup>a</sup> | ≥ 0.8 | 1.10 |
| DScore<sup>a</sup> | ≥ 0.9 | 1.14 |
| Exposure<sup>a</sup> | ≤ 0.5 | 0.41 |
| Enclosure<sup>a</sup> | ≥ 0.7 | 0.84 |
| Balance<sup>a</sup> | ≥ 1.0 | 2.60 |
| Volume<sup>b</sup> | ≥ 300 Å<sup>3</sup> | 1040 Å<sup>3</sup> |
| Depth<sup>b</sup> | ≥ 10 Å | 12.1 Å |
| SimpleScore<sup>b</sup> | ≥ 0.5 | 0.72 |
| DrugScore<sup>b</sup> | ≥ 0.5 | 0.51 |
| Descriptors computed using SiteMap<sup>a</sup> and DogSiteScorer<sup>b</sup> |  |  |

**Supp. Figure 4.**
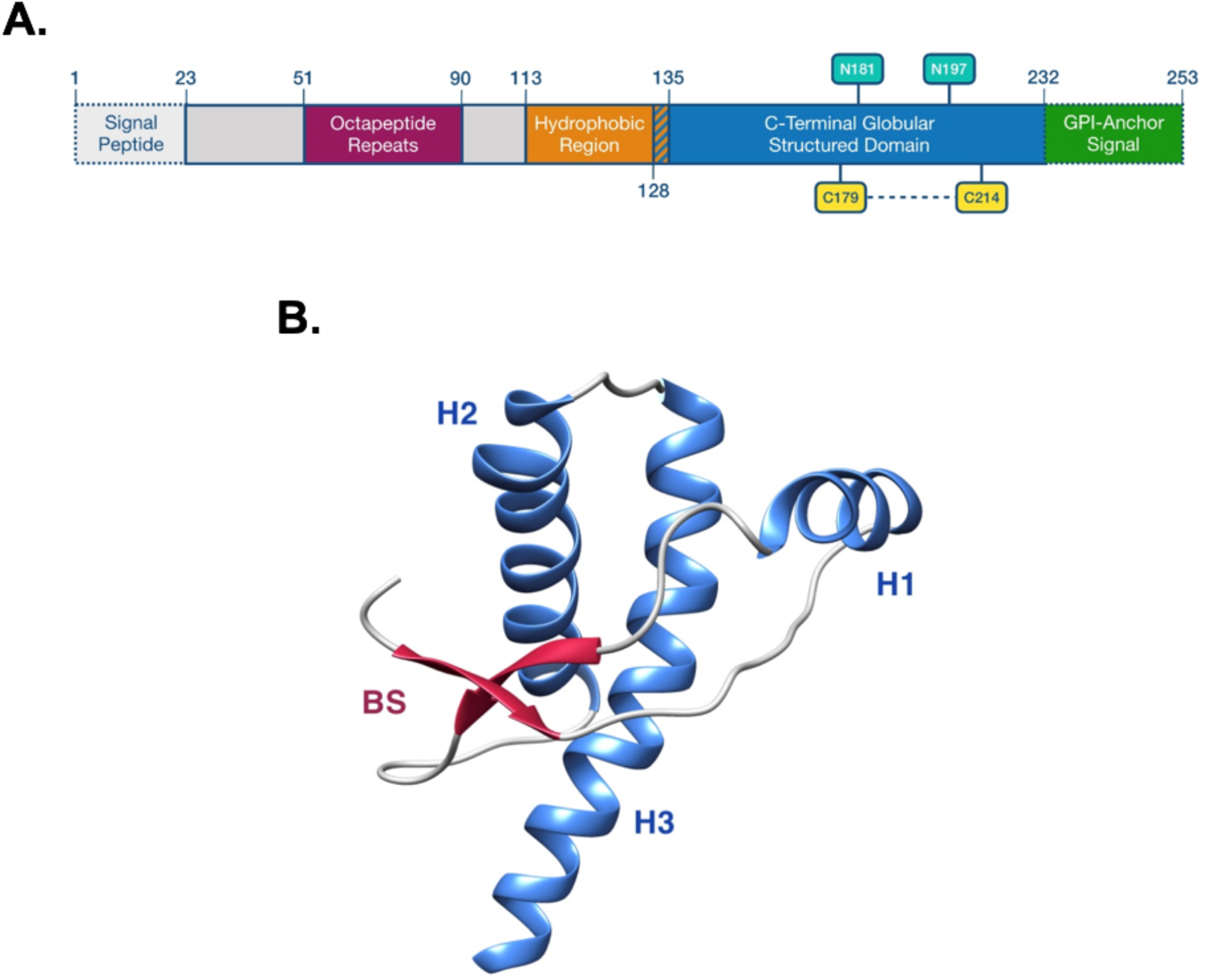
Representation of PrP structure. **A**. The picture illustrates a schematic representation of the structure of human PrP. The protein is organized as follows: a signal peptide (residues 1-22, light gray), removed during the biosynthesis of the protein in the ER, precedes five histidine-containing octapeptide repeats (residues 51-89, purple), which can bind divalent ions. The central region of the molecule includes a highly conserved hydrophobic domain (residues 113-134, orange). The C-terminal, globular domain (solved by NMR, PDB 1QLX, shown in panel **B**) includes two short anti-parallel β-strands (residues 127–129; and 166–168, light purple) and three α-helices (H1, H2, and H3, residues 143-152, 171-191 and 199-221, respectively; light blue). A C-terminal peptide (residues 232–253, green) is removed to allow the attachment of a glycosylphosphatidylinositol (GPI) moiety, which anchors the protein to the outer leaflet of the plasma membrane. The globular domain also contains two N-linked oligosaccharide chains (at Asn-181 and Asn-197, light green boxes) and a disulfide bond between residues 179 and 214 (yellow boxes).

**Supp. Movie.**
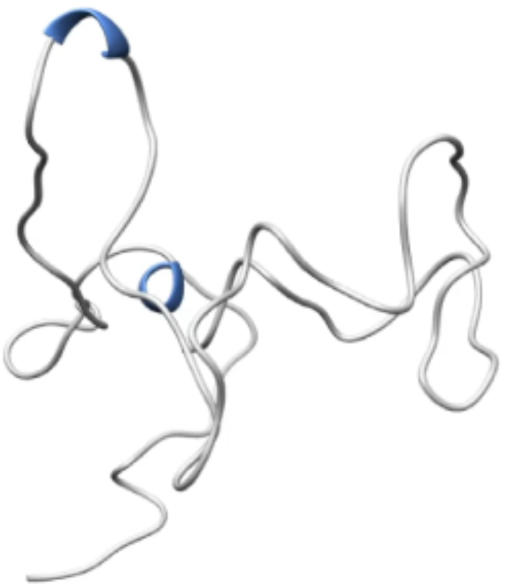
Full-atomistic reconstruction of PrP folding. Visualization of a representative least biased trajectory reconstructing the entire sequence of events leading the PrP polypeptide to reach the native state.

**Supp. Figure 5.**
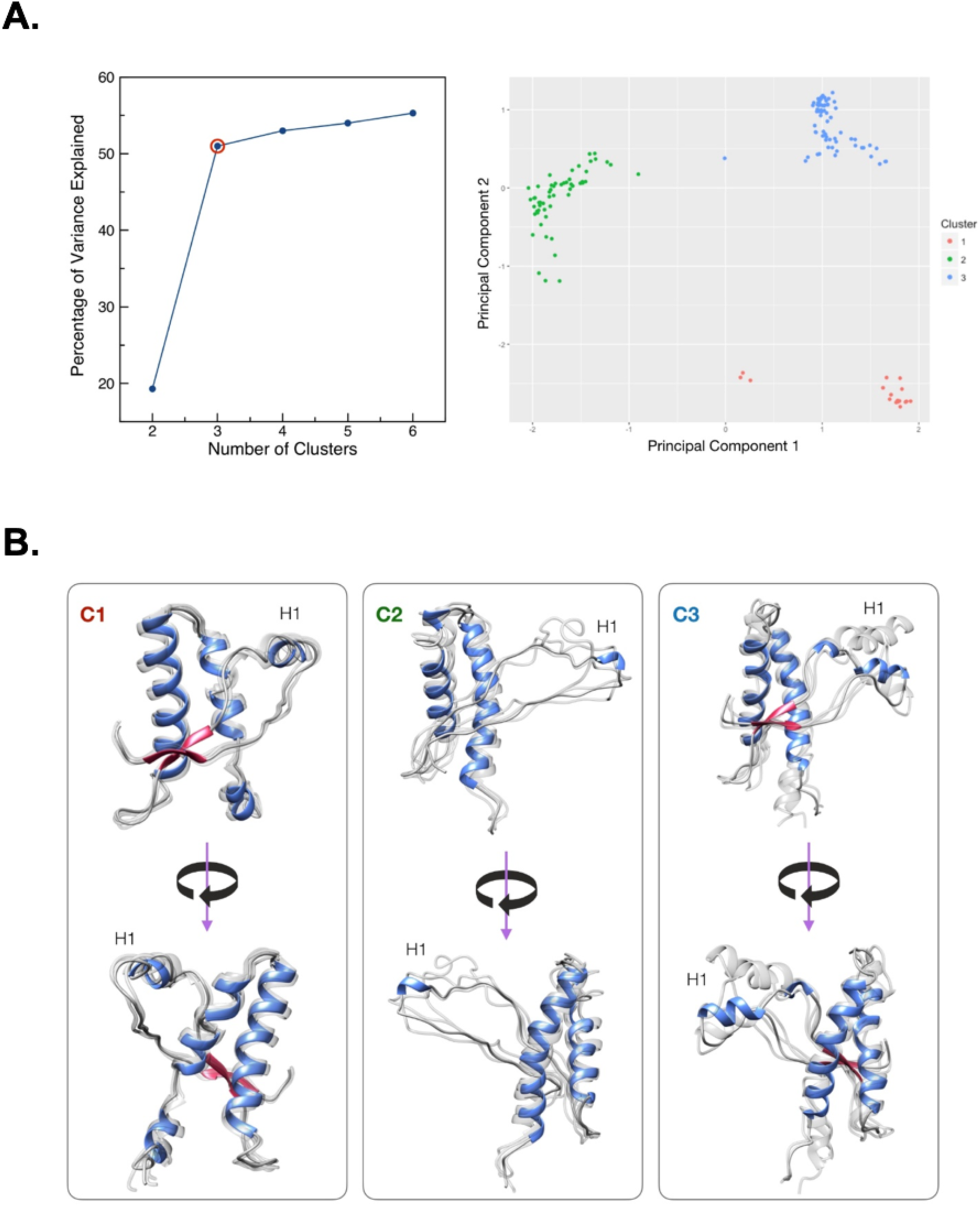
Clustering of PrP folding intermediate conformations. **A**. The left panel shows an elbow plot displaying the percentage of variance explained as a function of the number of clusters. The elbow criterion selects the number of clusters for *k*-mean at which the marginal gain obtained from the addition of a cluster drops (in this case *k* = 3, red circle). The right panel shows the clustering result represented in the principal component space. Conformations belonging to cluster 1, 2 and 3 are depicted in red, green, and blue, respectively. **B**. Structures corresponding to the three cluster (C) centers (helices, H, colored blue). Configurations obtained from C1, sampled from 2 out of 9 least biased trajectories, are characterized by docking of H1 onto H2 and H3, and an unproperly folded H2-H3 contact region. Both configurations C2 and C3 are characterized by a missing contact region between H1 and H2-H3, as compared to the native state, although the C3 conformations were largely the most represented, being sampled from 7 out of 9 least biased trajectories (while the C2 conformation only in 1 out of 9). Thus, the C3 center was selected for the following in-silico analyses.

**Supp. Table 3.**
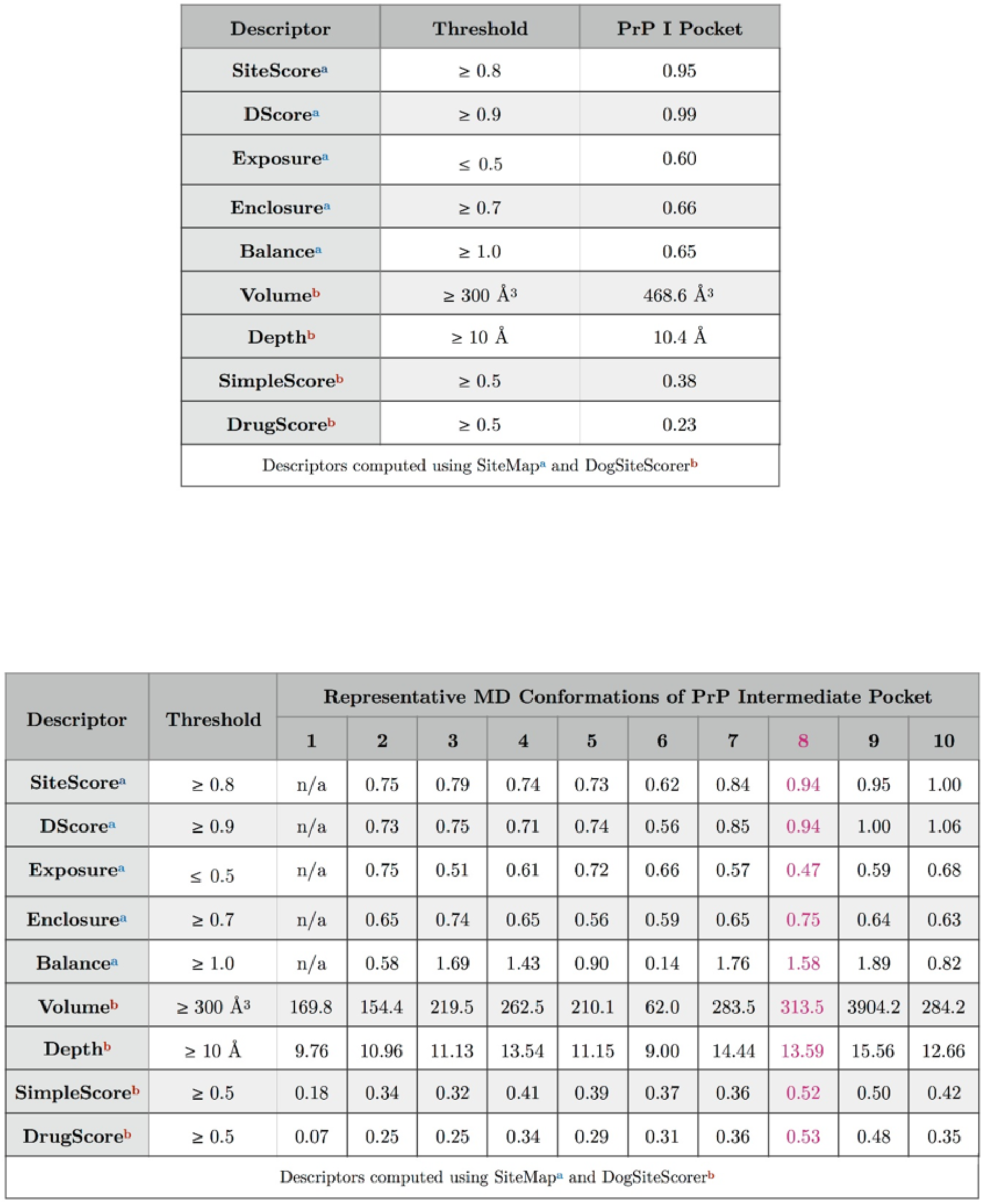
Druggability descriptors of the pocket in the PrP folding intermediate. The binding site identified in the PrP folding intermediate was refined by MD simulations. The table illustrates the computed SiteMap and DogSiteScorer descriptors for the ten representative MD pocket conformations. The analysis highlights the refined site in the MD conformation n.8, which possesses the desired thresholds for a well-defined, solvent-accessible druggable ligand-binding site. Noteworthy, this pocket was not present in the native form of PrP.

**SUPP. FIGURE 5**
| Descriptor | Threshold | PrP I Pocket |
| --- | --- | --- |
| SiteScore<sup>a</sup> | ≥ 0.8 | 0.95 |
| DScore<sup>a</sup> | ≥ 0.9 | 0.99 |
| Exposure<sup>a</sup> | ≤ 0.5 | 0.60 |
| Enclosure<sup>a</sup> | ≥ 0.7 | 0.66 |
| Balance<sup>a</sup> | ≥ 1.0 | 0.65 |
| Volume<sup>b</sup> | ≥ 300 Å<sup>3</sup> | 468.6 Å<sup>3</sup> |
| Depth<sup>b</sup> | ≥ 10 Å | 10.4 Å |
| SimpleScore<sup>b</sup> | ≥ 0.5 | 0.38 |
| DrugScore<sup>b</sup> | ≥ 0.5 | 0.23 |
| Descriptors computed using SiteMap<sup>a</sup> and DogSiteScorer<sup>b</sup> |  |  |

SUPP. TABLE 3
| Descriptor | Threshold | Representative MD Conformations of PrP Intermediate Pocket |  |  |  |  |  |  |  |  |  |
| --- | --- | --- | --- | --- | --- | --- | --- | --- | --- | --- | --- |
|  |  | 1 | 2 | 3 | 4 | 5 | 6 | 7 | 8 | 9 | 10 |
| SiteScore<sup>a</sup> | ≥ 0.8 | n/a | 0.75 | 0.79 | 0.74 | 0.73 | 0.62 | 0.84 | 0.94 | 0.95 | 1.00 |
| DScore<sup>a</sup> | ≥ 0.9 | n/a | 0.73 | 0.75 | 0.71 | 0.74 | 0.56 | 0.85 | 0.94 | 1.00 | 1.06 |
| Exposure<sup>a</sup> | ≤ 0.5 | n/a | 0.75 | 0.51 | 0.61 | 0.72 | 0.66 | 0.57 | 0.47 | 0.59 | 0.68 |
| Enclosure<sup>a</sup> | ≥ 0.7 | n/a | 0.65 | 0.74 | 0.65 | 0.56 | 0.59 | 0.65 | 0.75 | 0.64 | 0.63 |
| Balance<sup>a</sup> | ≥ 1.0 | n/a | 0.58 | 1.69 | 1.43 | 0.90 | 0.14 | 1.76 | 1.58 | 1.89 | 0.82 |
| Volume<sup>b</sup> | ≥ 300 Å<sup>3</sup> | 169.8 | 154.4 | 219.5 | 262.5 | 210.1 | 62.0 | 283.5 | 313.5 | 3904.2 | 284.2 |
| Depth<sup>b</sup> | ≥ 10 Å | 9.76 | 10.96 | 11.13 | 13.54 | 11.15 | 9.00 | 14.44 | 13.59 | 15.56 | 12.66 |
| SimpleScore<sup>b</sup> | ≥ 0.5 | 0.18 | 0.34 | 0.32 | 0.41 | 0.39 | 0.37 | 0.36 | 0.52 | 0.50 | 0.42 |
| DrugScore<sup>b</sup> | ≥ 0.5 | 0.07 | 0.25 | 0.25 | 0.34 | 0.29 | 0.31 | 0.36 | 0.53 | 0.48 | 0.35 |
| Descriptors computed using SiteMap<sup>a</sup> and DogSiteScorer<sup>b</sup> |  |  |  |  |  |  |  |  |  |  |  |

**Supp. Table 4.** Virtual screening results. The table reports the Asinex ID code, laboratory-internal code (SM code), and predicted HYDE affinity range (K_iHYDE_) for the 30 virtual hits selected as PrP intermediate ligands. The virtual screening campaign was carried out by means of BioSolveIT tools integrated into an in-house developed KNIME workflow. A funnel-like approach involving the prediction of pharmacodynamics, physicochemical, and ADMET properties as well as molecular diversity and 3D visualization of ligand binding interactions guided the hit selection procedure.

| Compound ID | SM code | K<sub>iHYDE</sub> |
| --- | --- | --- |
| ASN19380133 | 875 | 40275 nM > K<sub>i</sub> > 405 nM |
| BAS00670252 | 923 | 17266 nM > K<sub>i</sub> > 174 nM |
| BAS01809156 | 924 | 468 nM > K<sub>i</sub> > 5 nM |
| BAS03200180 | 925 | 23544 nM > K<sub>i</sub> > 237 nM |
| BAS00312802 | 926 | 43535 nM > K<sub>i</sub> > 438 nM |
| BAS02102991 | 927 | 13673 nM > K<sub>i</sub> > 138 nM |
| BAS00781325 | 929 | 538 nM > K<sub>i</sub> > 5 nM |
| BAS00287978 | 930 | 2618 nM > K<sub>i</sub> > 26 nM |
| BAS01849776 | 931 | 2795 nM > K<sub>i</sub> > 28 nM |
| BAS00855482 | 932 | 23755 nM > K<sub>i</sub> > 239 nM |
| BAS01029925 | 933 | 126 nM > K<sub>i</sub> > 1 nM |
| ASN02751074 | 934 | 5153 nM > K<sub>i</sub> > 52 nM |
| BAS05541419 | 935 | 7775 nM > K<sub>i</sub> > 78 nM |
| ASN02241765 | 936 | 6024 nM > K<sub>i</sub> > 61 nM |
| ASN15755504 | 937 | 2296 nM > K<sub>i</sub> > 23 nM |
| BAS15232183 | 938 | 5101 nM > K<sub>i</sub> > 51 nM |
| BAS01095083 | 939 | 20390 nM > K<sub>i</sub> > 205 nM |
| ASN16356774 | 940 | 6121 nM > K<sub>i</sub> > 62 nM |
| ASN05397475 | 941 | 2020 nM > K<sub>i</sub> > 20 nM |
| BAS01195857 | 942 | 1583 nM > K<sub>i</sub> > 16 nM |
| BAS17126510 | 943 | 26641 nM > K<sub>i</sub> > 268 nM |
| ASN17325626 | 944 | 5761 nM > K<sub>i</sub> > 58 nM |
| BAS03374806 | 945 | 19591 nM > K<sub>i</sub> > 197 nM |
| BAS01515776 | 946 | 20171 nM > K<sub>i</sub> > 203 nM |
| BAS00382671 | 947 | 40423 nM > K<sub>i</sub> > 407 nM |
| BAS01990851 | 948 | 15124 nM > K<sub>i</sub> > 152 nM |
| BAS03321293S | 949 | 1811 nM > K<sub>i</sub> > 18 nM |
| BAS01923101 | 950 | 25764 nM > K<sub>i</sub> > 259 nM |
| BAS01058340 | 951 | 7910 nM > K<sub>i</sub> > 80 nM |
| BAS09812417 | 952 | 11821 nM > K<sub>i</sub> > 119 nM |

**Supp. Figure 6.**
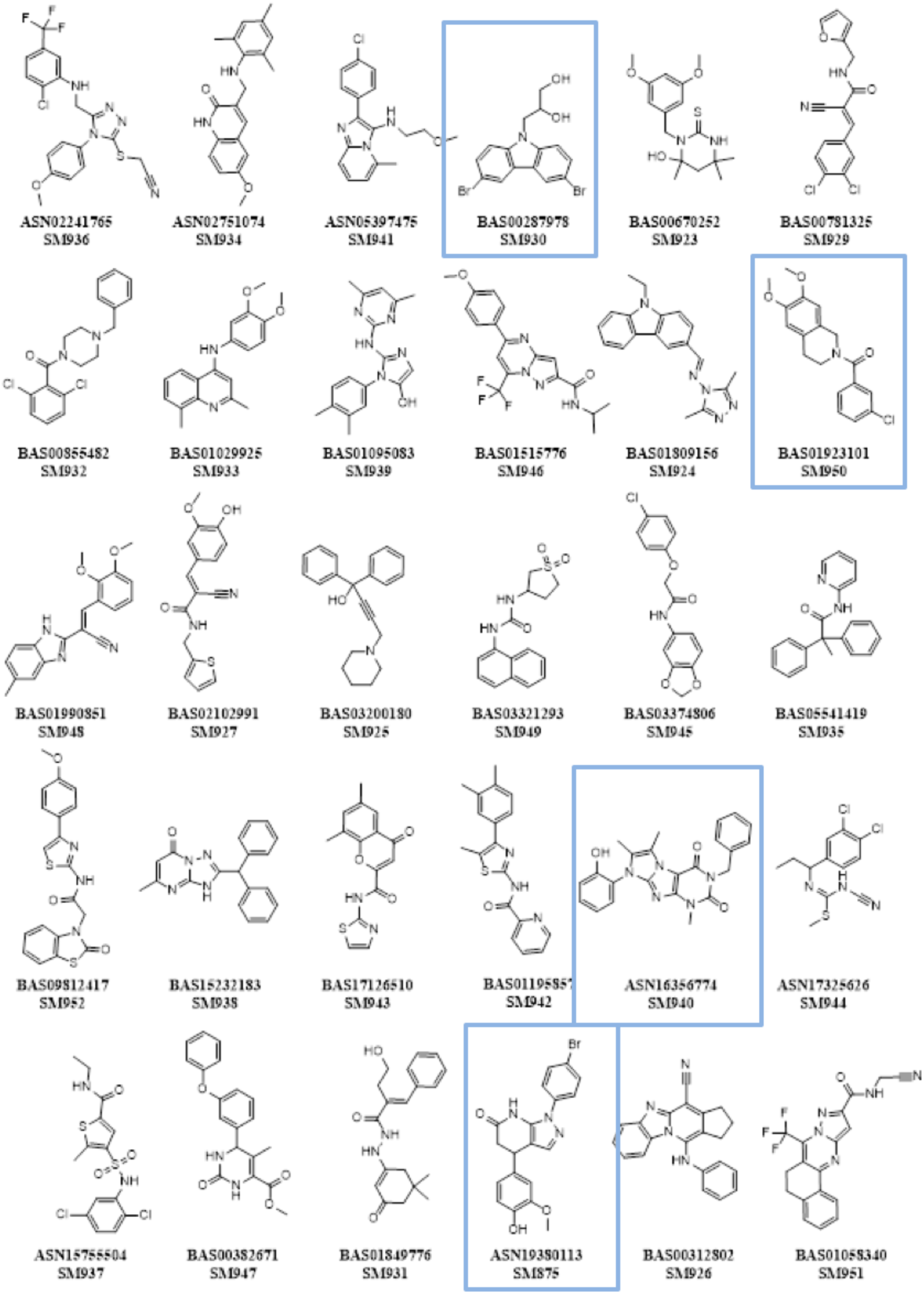
Chemical structures of the 30 virtual hits. The figure illustrates the 2D structure, the Asinex ID code, and the *in-house* code (SM) for the 30 virtual hits selected for biological evaluation. Each molecule was tested for its ability to lower the expression of full-length PrP (assayed by western blotting) without decreasing the expression of NEGR-1, a GPI-anchored protein following the same expression pathway of PrP and used here as control of effect specificity. Four compounds resulted as positive in these analyses: SM875, SM930, SM940, and SM950 (blue boxes).

**Supp. Figure 7.**
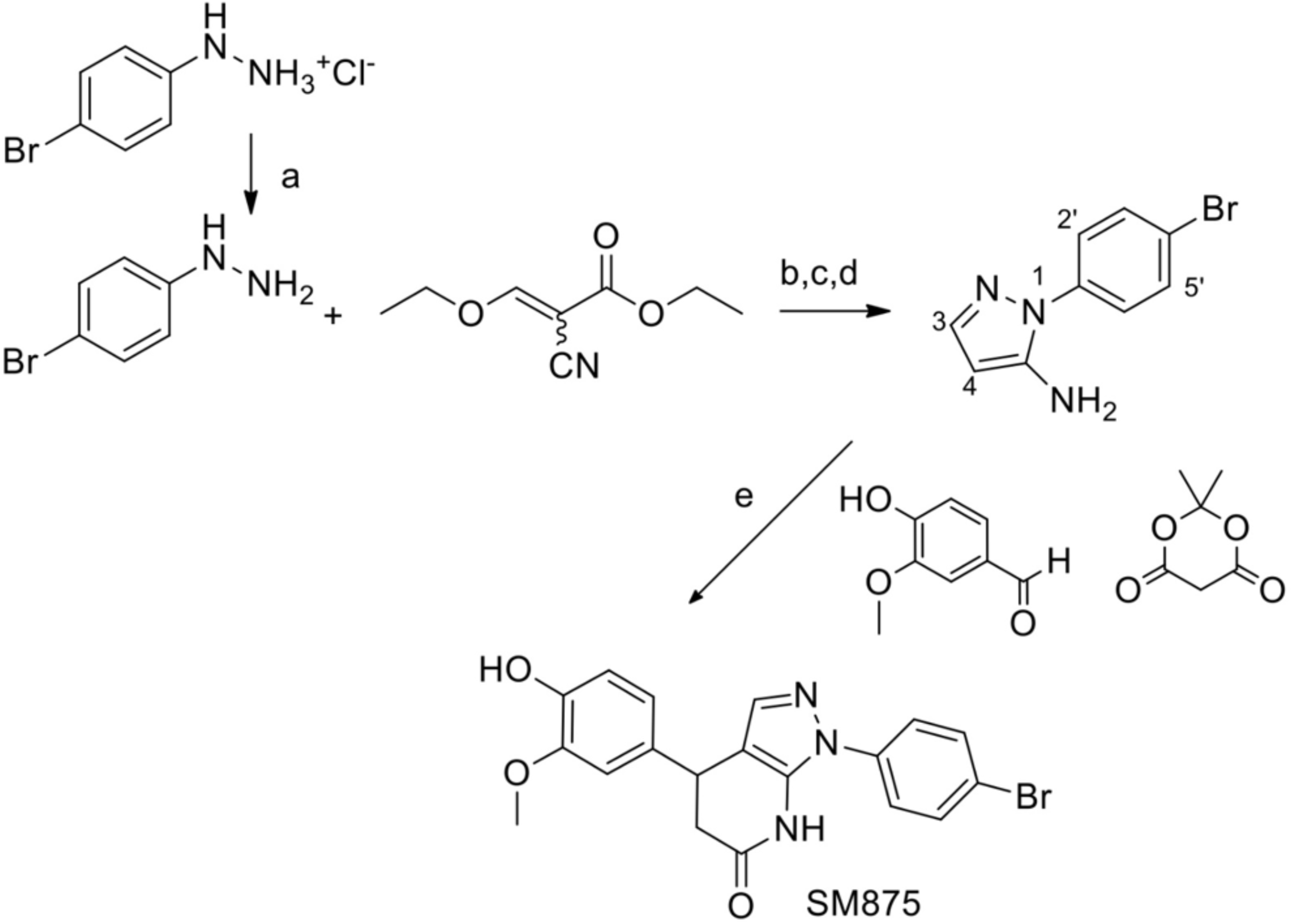
Synthetic scheme of SM875. Reagents and conditions: (a) NaHCO_3_ aqueous solution; (b) ethanol reflux, 2 h; (c) 1:1 methanol/aqueous 2M NaOH, reflux 1h; (d) 180°C, 10 min; 61%; (e) ethanol reflux 2.5 h, HPLC purification, 25%. Arbitrary numbering of synthetic intermediate [1-(4-bromophenyl)-1H-pyrazol-5-amine] is used for ^1^H-NMR assignment: [δ_H_ 7.58(d, J 8.7 Hz, H-3’and H-5’), 7.47(d, J 8.7 Hz, H-2’and H-6’), 7.41(s, H-3), 5.62 (s, H-4)].

**Supp. Figure 8.**
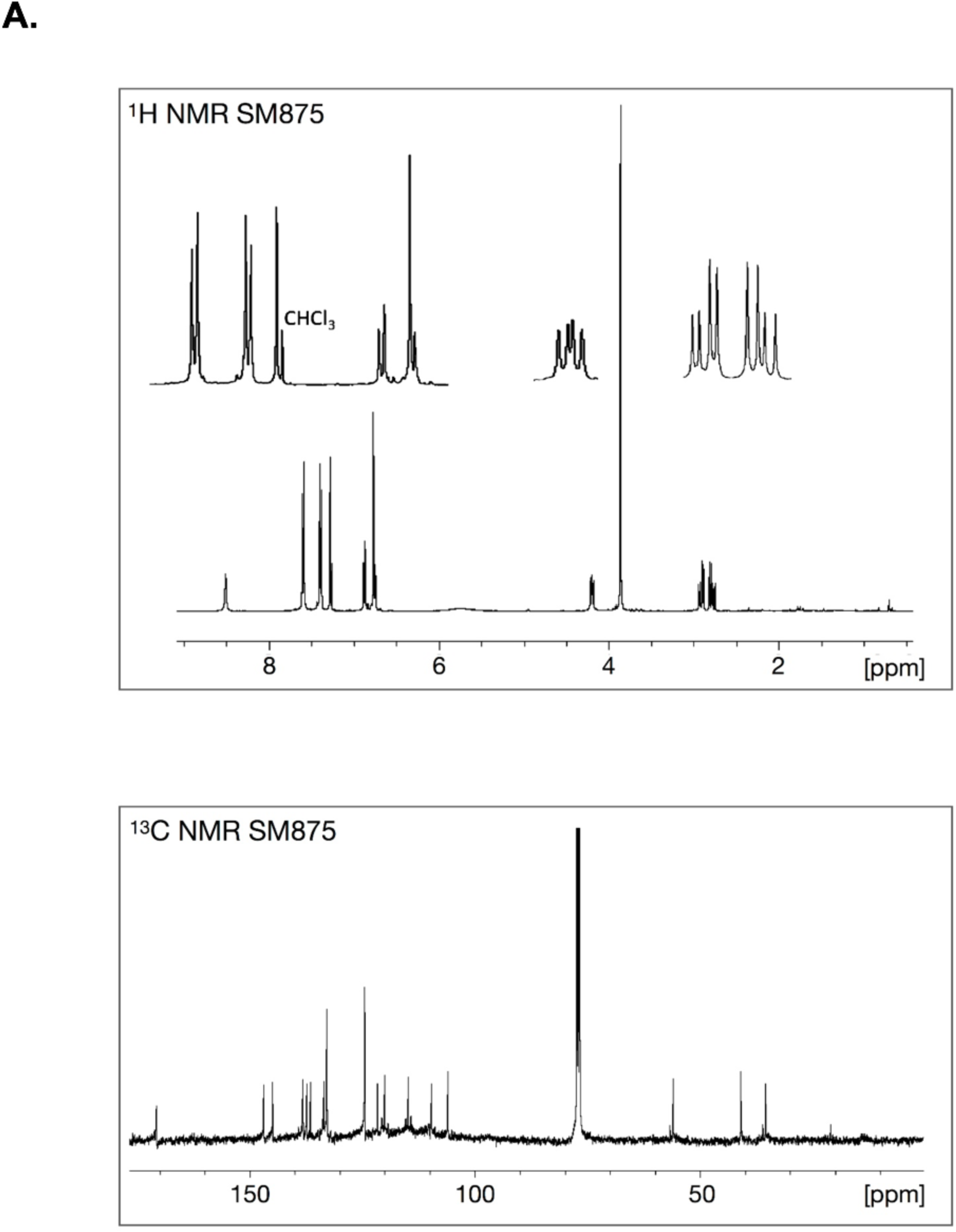
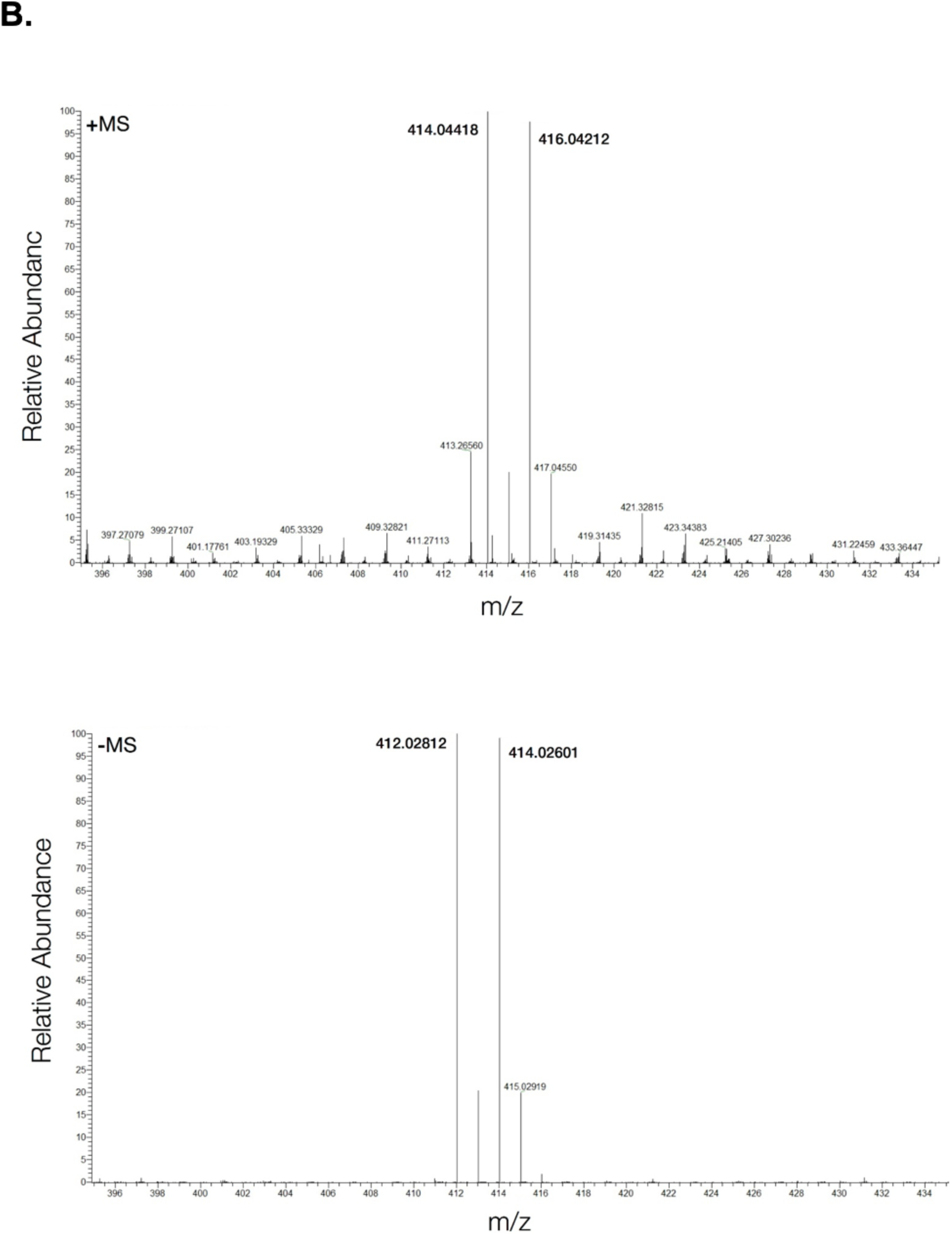
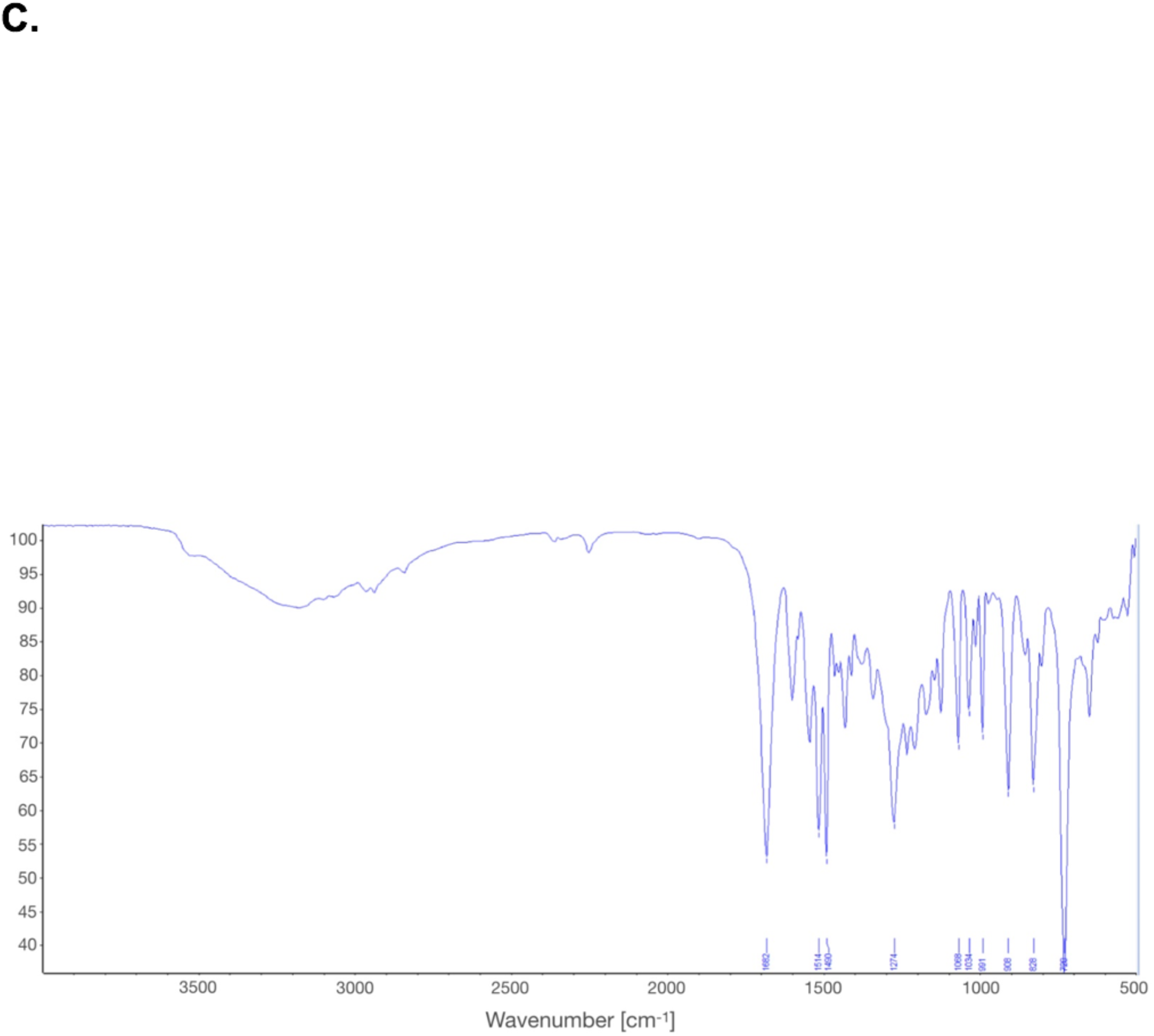
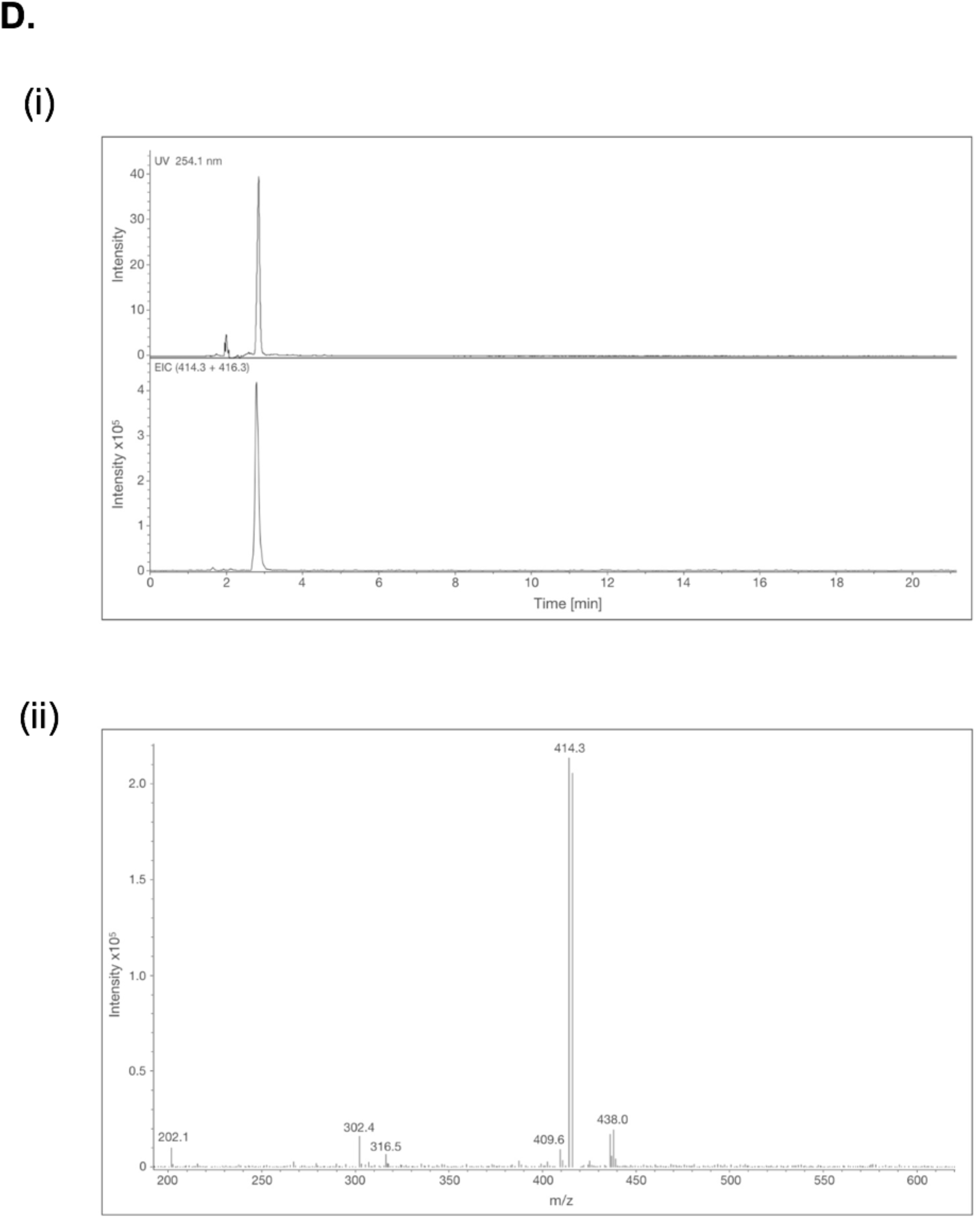
Structural characterization of SM875. **A**. ^1^H-NMR spectrum of SM875 (upper panel, 400 MHz) in CDCl_3_ with residual CHCl_3_ at 7.25 ppm. ^13^C-NMR spectrum (lower panel, 100 MHz) of SM875 in CDCl_3_. **B**. Upper panel: high resolution ESI-MS positive ion mode measurement of SM875 (methanol solution) by direct infusion: [M+H]^+^ ions at *m/z* 414.0443 (calculated for C_19_H_15_^79^BrN_3_O_3_ = 414.0448) and m/z 416.0421 (calculated for C_19_H_15_^81^BrN_3_O_3_ = 416.0427). Lower panel: high resolution ESI-MS negative ion mode measurement of SM875 (methanol solution) by direct infusion : [M-H]^-^ ions at *m/z* 412.0282 (calculated for C_19_H_15_^79^BrN_3_O_3_ = 412.0302) and m/z 414.0260 (calculated for C_19_H_15_^81^BrN_3_O_3_ = 414.0282); **C**. FT-IR spectrum of SM875 (cm^-1^): 1681s, 1543s, 1515s, 1490s, 1274s, 991m, 828m, 728 vs; **D**. Upper panel: UV chromatogram at l =254 nm of SM875 (LC-ESI(+)-MS-UV,; RP18, acetonitrile/water 70:30, flow 1 mL·min^-1^). Lower panel: MS chromatogram by extracted-ion current of the [M+H]^+^ peaks at *m/z* 414 and 416 (1:1) corresponding to the two expected major isotopomers (1:1) C_19_H_17_^79^BrN_3_O_3_ and C_19_H_17_^81^BrN_3_O_3_.

**Supp. Table 5.**
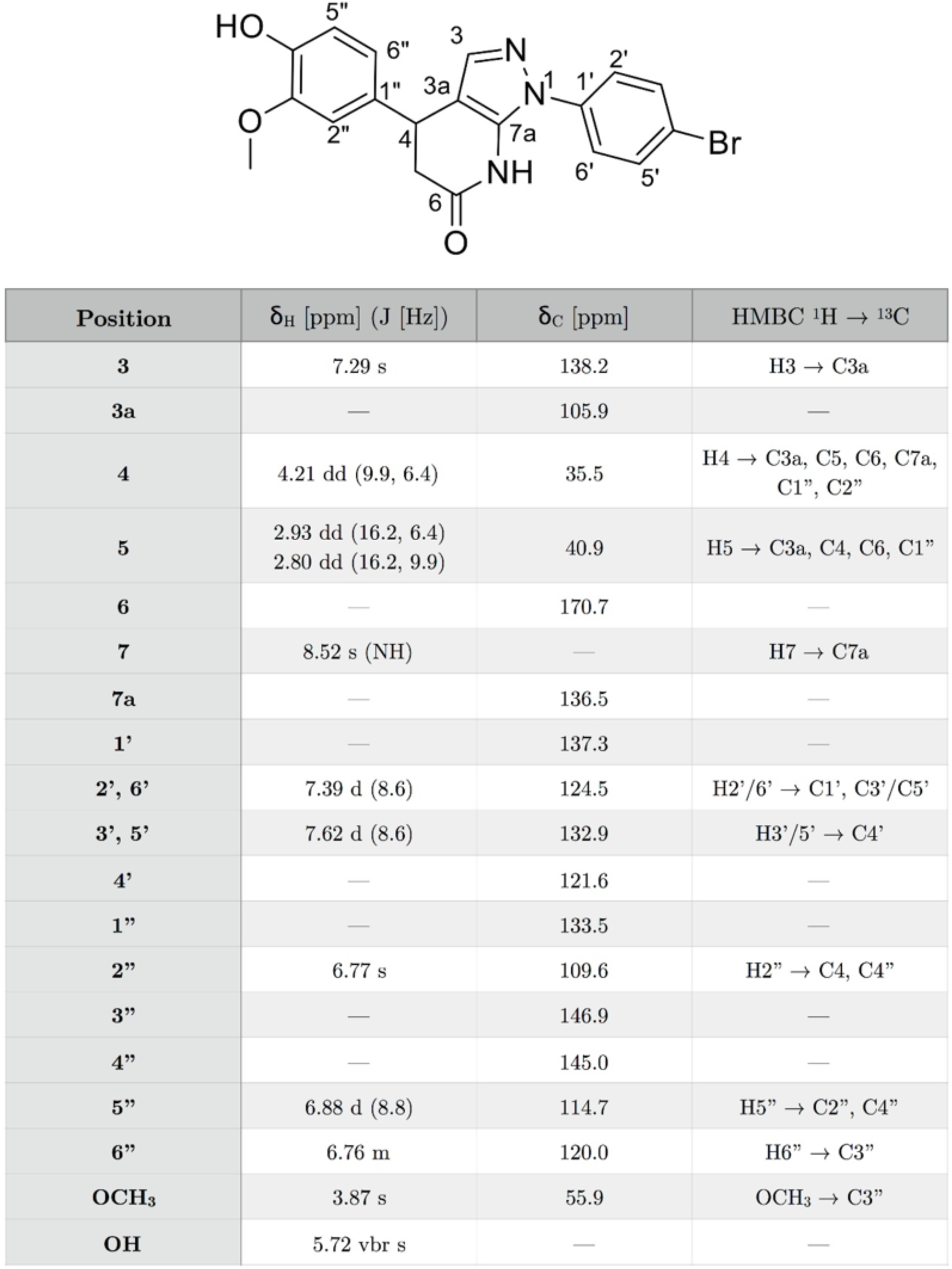
NMR spectral data of SM875. NMR data obtained in CDCl_3_ (^1^H at 400 MHz, ^13^C at 100 MHz) are reported. Resonances are assigned according to the carbon numbering shown on the chemical structure. In the second column are reported the ^1^H NMR parameters [δ_H_ (J)], in the third column the ^13^C chemical shifts (δ_C_) and in the fourth column the most relevant long-range heteronuclear correlations (obtained by 2D-NMR HMBC pulse sequence).

**Supp. Figure 9.**
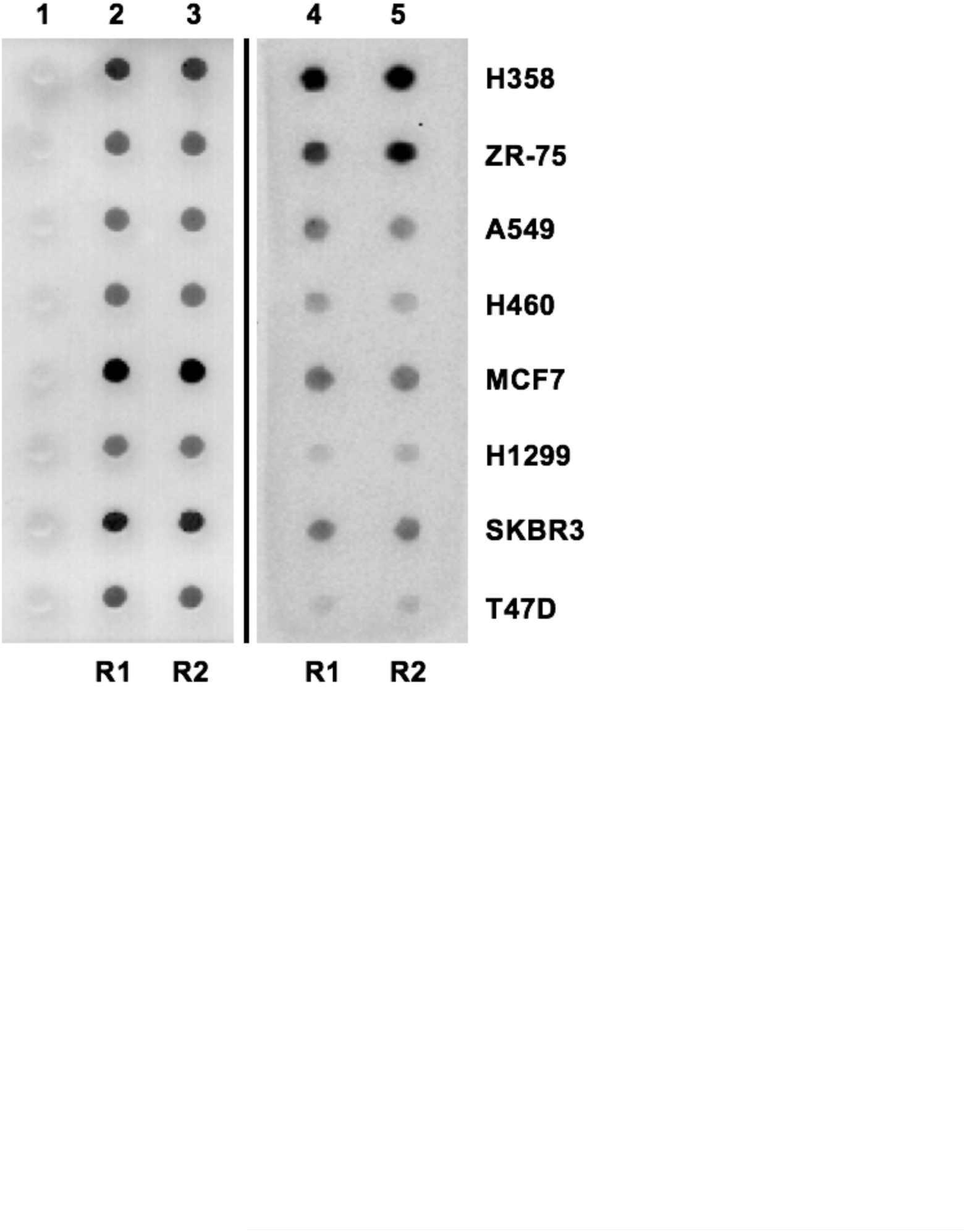
Evaluation of PrP expression in different cancer cell lines. The endogenous expression of PrP in eight different cultured cell lines (indicated) from lung and breast cancers belonging to the Human Tumor Cell Line collection of the National Cancer Institute (NCI) was evaluated by dot-blotting. Cell lysates (lanes 2-5) from two different experiments (R1 and R2) were spotted on to a PVDF membrane, together with a buffer control (lane 1), stained with Ponceau red (lanes 1-3) or incubated with anti-PrP primary antibody (D18) followed by an HRP-conjugated secondary antibody (lanes 4-5). ZR-75 show the highest expression of PrP among the different cell lines.

**Supp. Figure 10.**
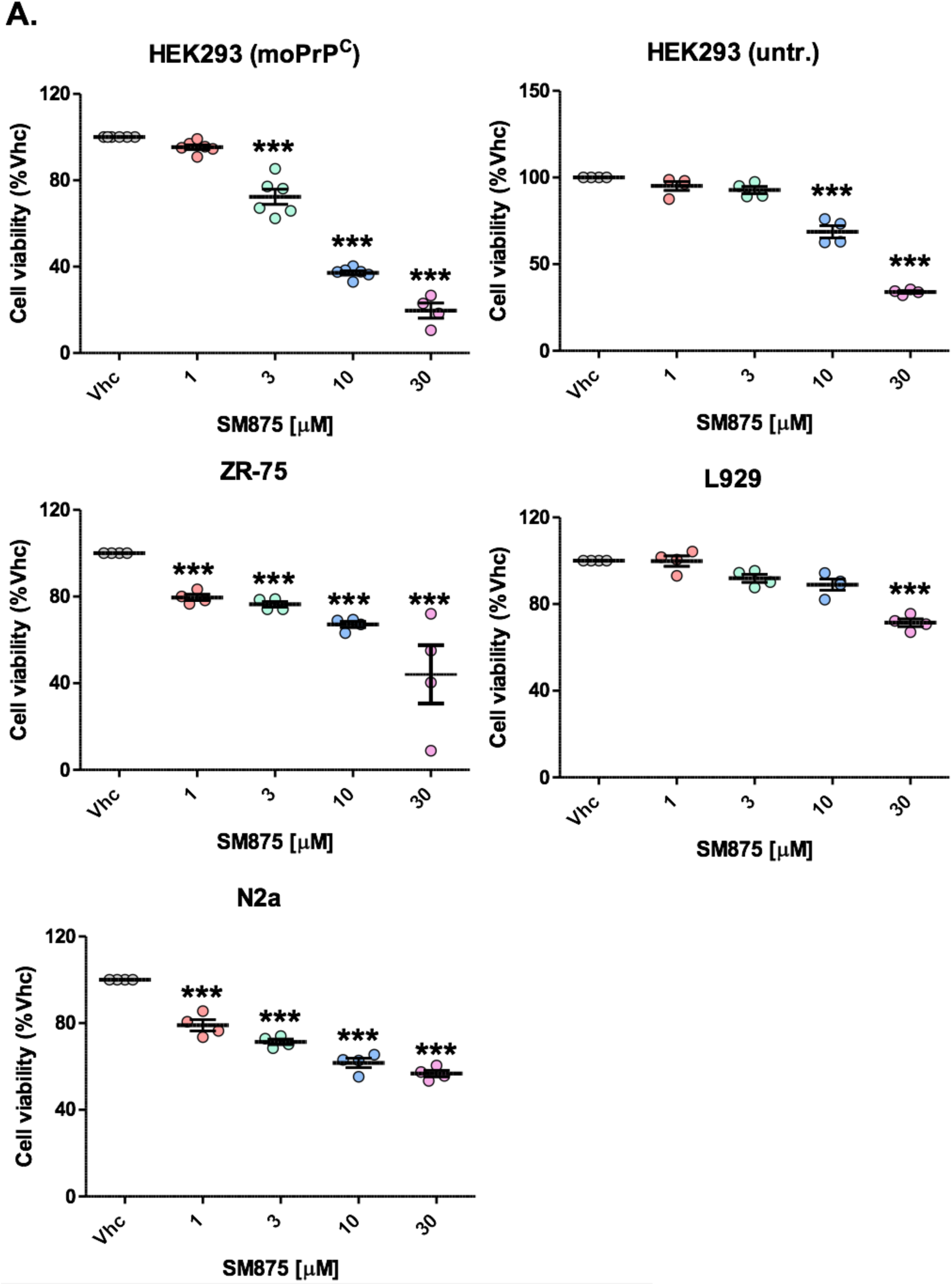
Estimation of SM875 cytotoxicity. In several experiments, we noticed evident cytotoxicity of SM875 at concentrations near to those at which the compound exerted PrP-lowering effects. While experimental controls in the different assays assess effect specificity, the observed cytotoxicity may still represent an important confounding factor. Therefore, we quantified the toxicity of the molecule in the different cell lines, and at the same incubation time and concentrations employed in the study, by using the MTT cell viability assay. Results show that SM875 variably affects cell viability in the different cell lines. In particular, the compound shows high cytotoxicity in moPrP-transfected HEK293 (∼28% reduction of cell viability at 3 µM), N2a (∼21% cell death at 1 µM) and ZR-75 cells (∼24-33% reduction between 1 and 10 µM). Conversely, SM875 appears to be much less toxic in untrasfected N2a cells (no significant alteration of cell viability at 3 µM, ∼32% reduction at 10 µM) and L929 fibroblasts (cell viability almost completely unaltered up to 10 µM, ∼29% reduction at 30 µM). These results clearly indicate that SM875 needs to be optimized to increase its potency and reduce its intrinsic toxicity, before becoming suitable for in vivo tests (*** p < 0.005).

**Supp. Figure 11.**
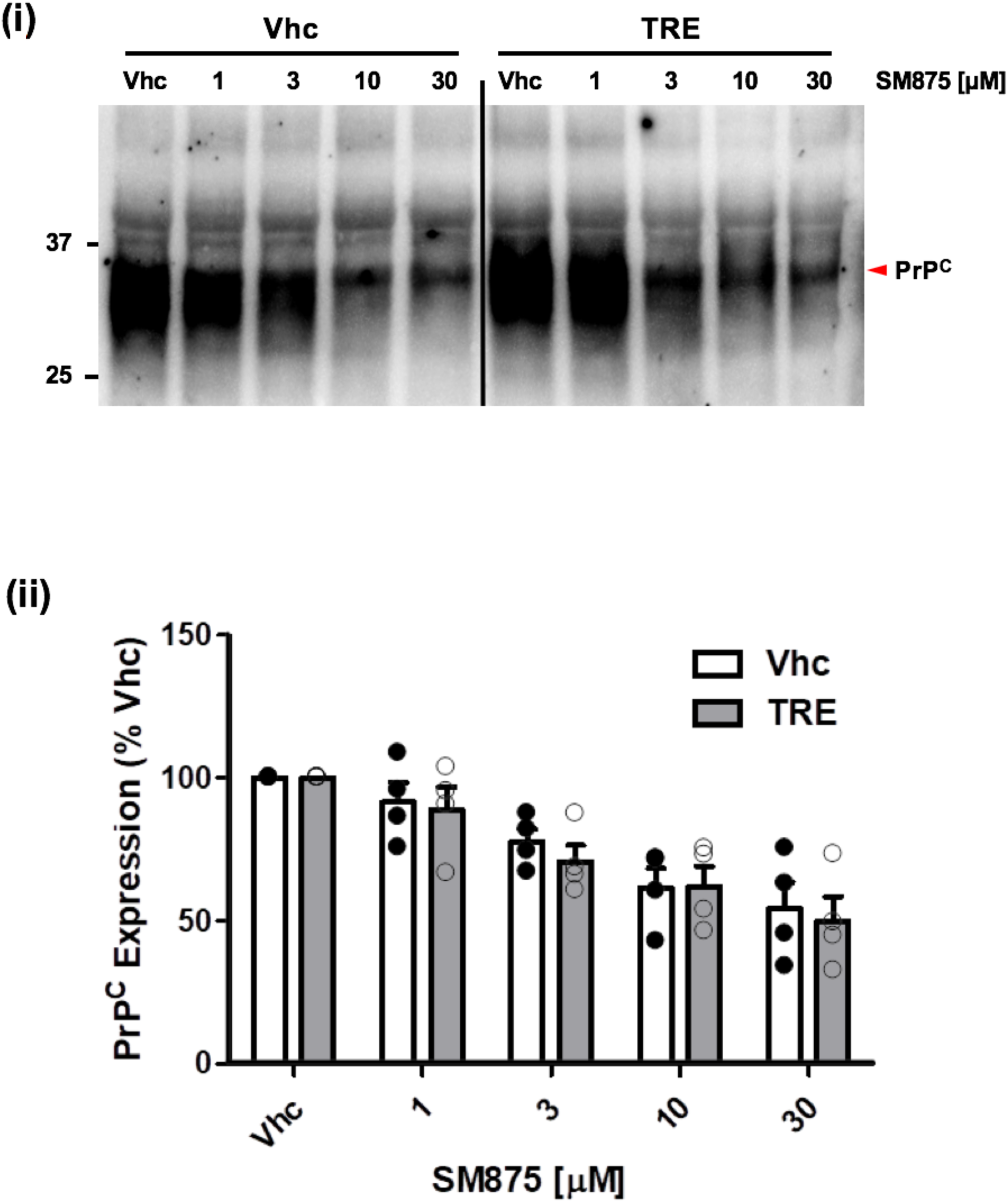
Stimulation of autophagy does not alter the PrP-lowering effects of SM875. Since inhibition of autophagy-mediated lysosomal degradation rescued the decrease of PrP induced by SM875, we sought also to test whether stimulating this pathway would alter the activity of the compound. Thus, we treated ZR-75 cells with different concentrations of SM875 (indicated) or vehicle (DMSO, volume equivalent), in the presence or absence of autophagy inducer TRE (100 µM) and detected PrP by western blotting **(i)**. Signals were detected by probing membrane blots with anti-PrP antibody (D18). We observed that SM875 lowers the levels of full-length PrP (indicated by the red arrow) in a dose-dependent fashion independently from the presence or absence of TRE; (**ii**) the graphs show the densitometric quantification of full-length PrP from independent replicates (n ≥ 4). Each signal was normalized on the corresponding total protein lane (detected by UV) and expressed as the percentage of vehicle (Vhc)-treated controls.

**Supp. Figure 12.**
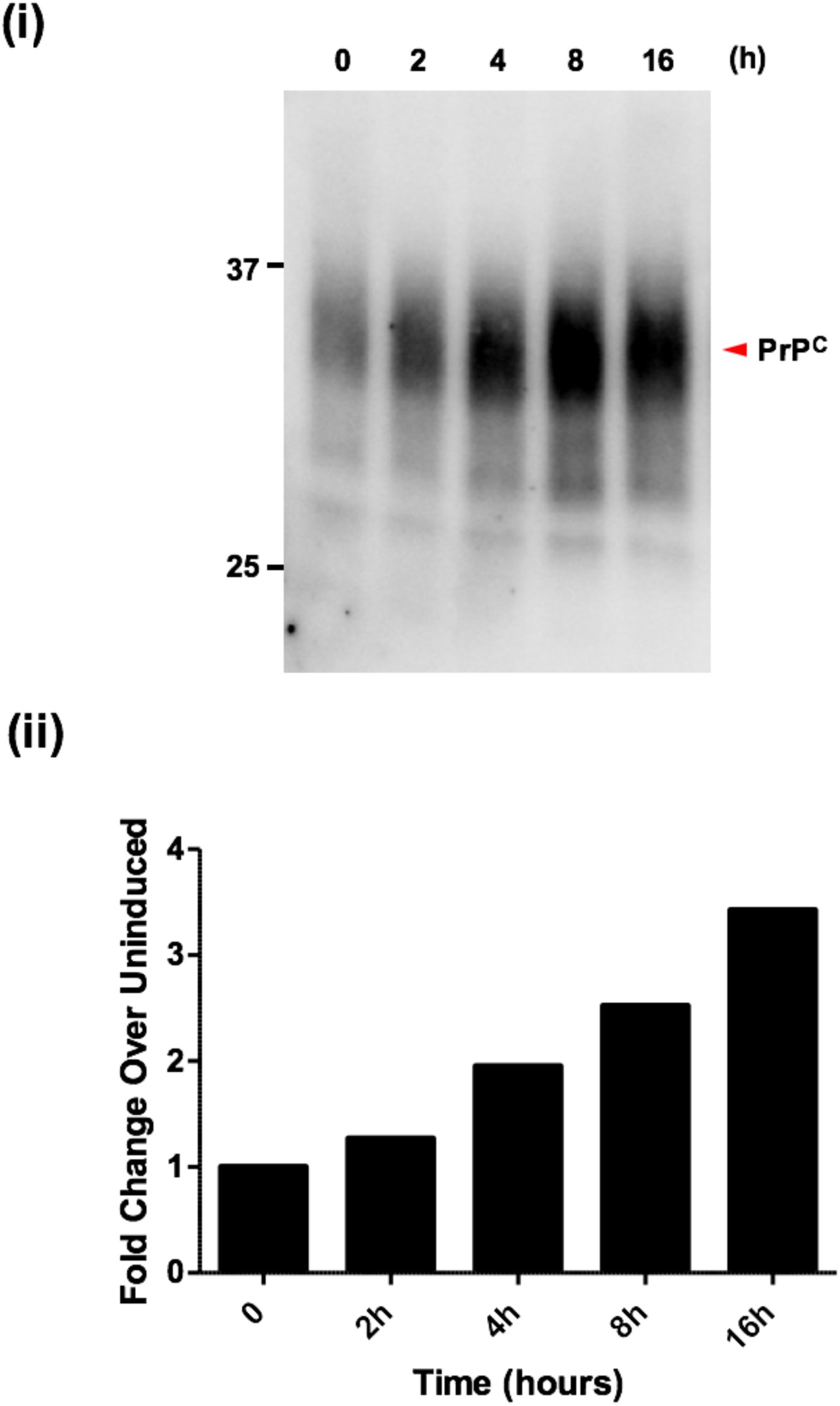
Expression of PrP in doxycycline-inducible RK13 cells. **(i)** The expression of PrP upon induction with doxycycline (0.01 mg/mL) was evaluated by western blotting over 16 h. Signals were detected by probing membrane blots with anti-PrP antibody (D18); **(ii)** The graph shows the densitometric quantification of full-length PrP, obtained by normalizing each signal on the corresponding total protein lane (detected by UV) and expressed as fold change of vehicle-treated control.

**Supp. Figure 13.**
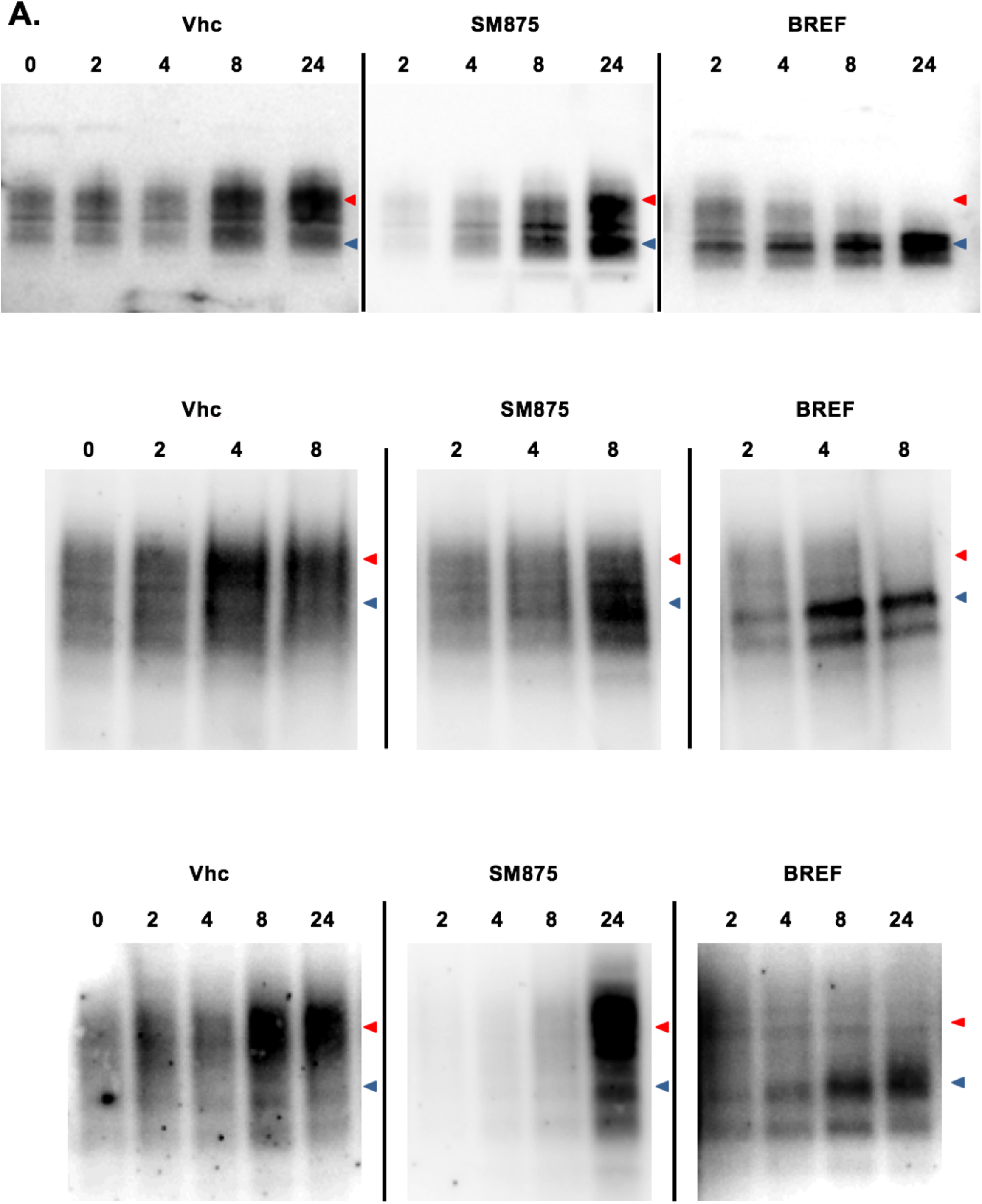
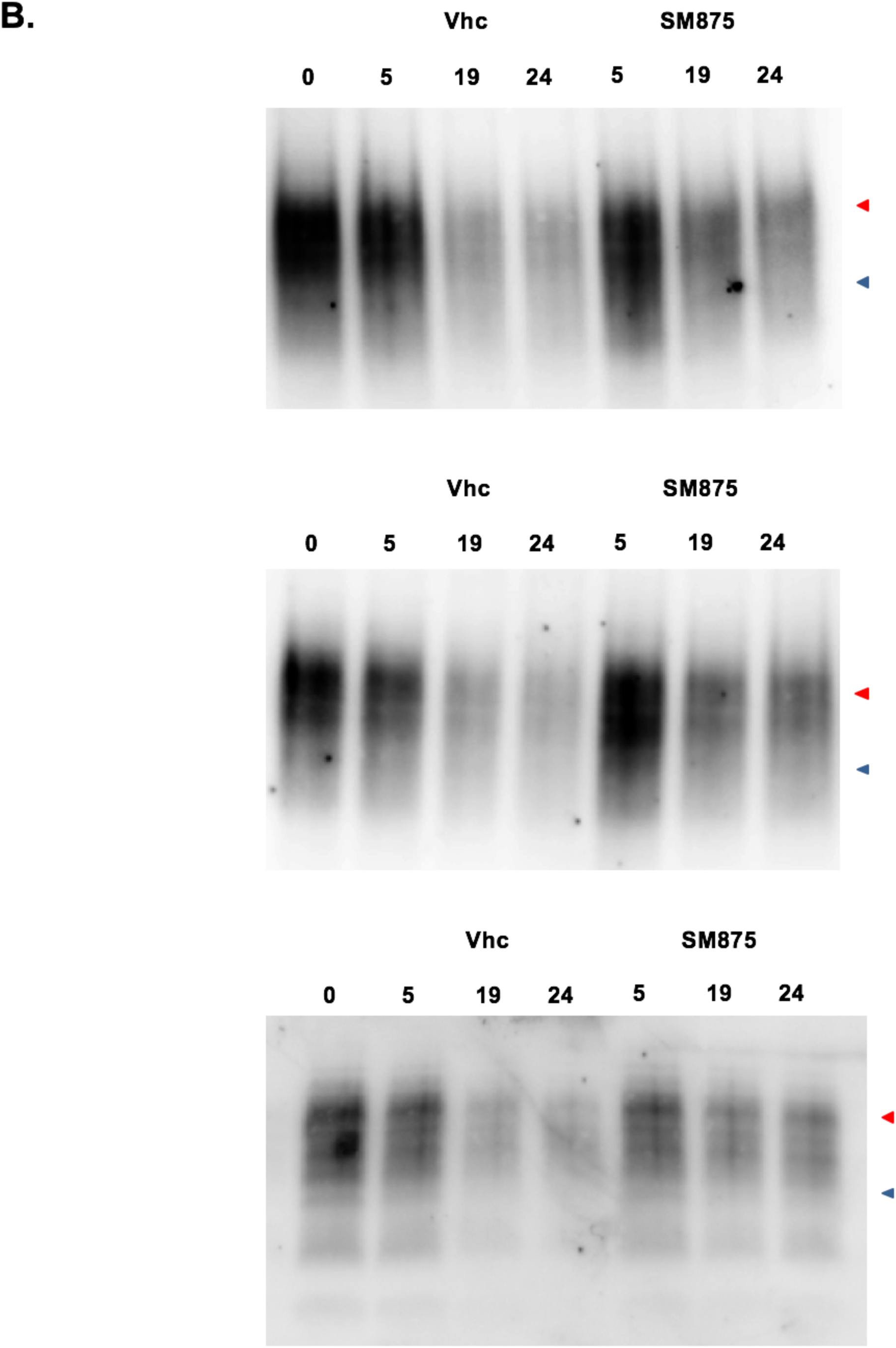
Replicates of experiments presented in figure 6. **A**. Individual replicates of experiments shown in Figure 6B, panel (i). **B**. Individual replicates of experiments presented in Figure 6B, panel (ii). Signals were detected by probing membrane blots with anti-PrP antibody (D18).

**Supp. Figure 14.**
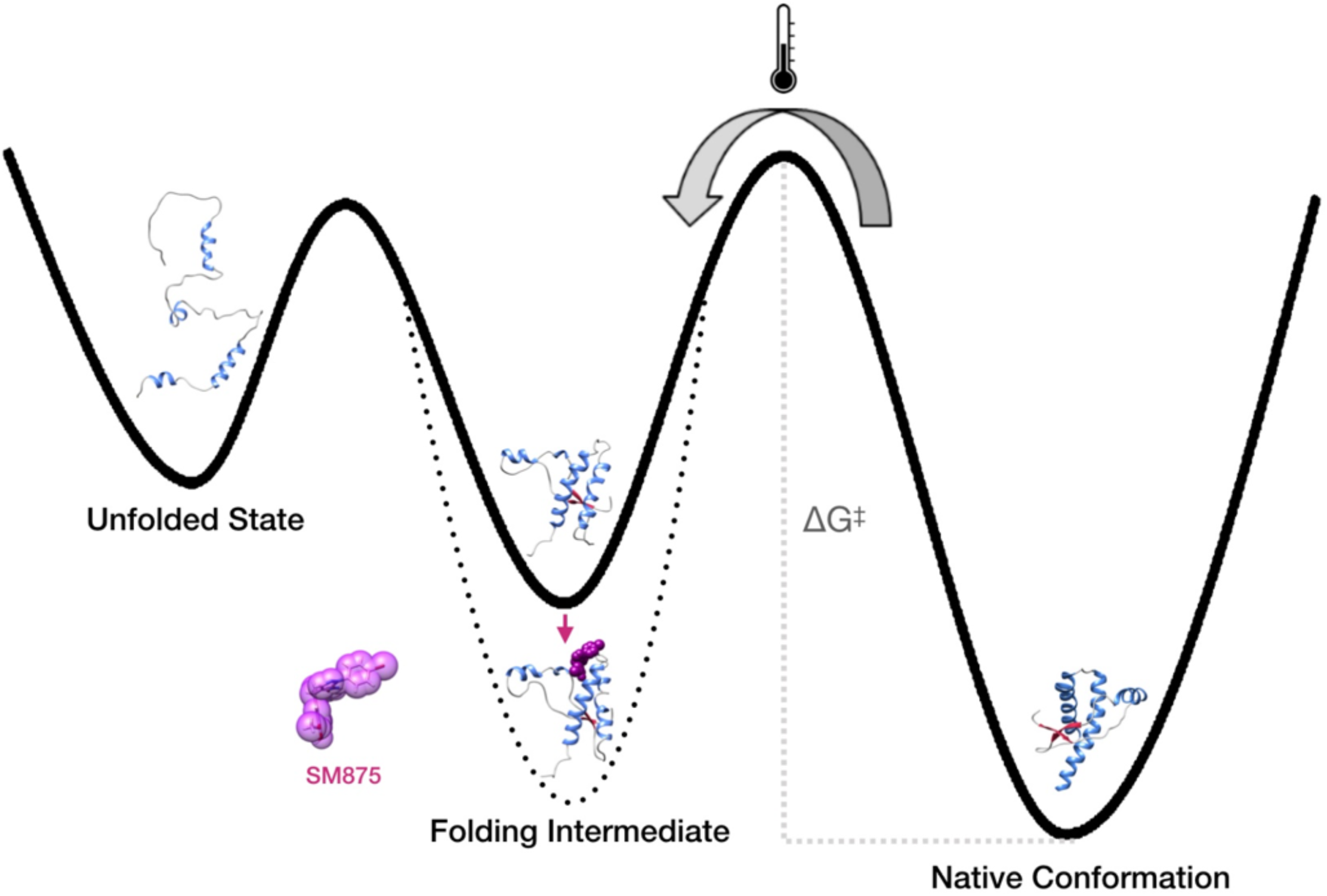
Scheme of the experimental layout used to try co-crystallizing the PrP folding intermediate bound to SM875. In an attempt to structurally characterize the PrP folding intermediate identified in-silico, we designed a novel experimental paradigm to induce the appearance of alternative PrP conformers, separated from the native state by a relevant energy barrier (ΔG^‡^). This goal was tentatively achieved by the partial, thermal unfolding of mouse recombinant PrP (111-230, 800 µM; temperature raised to 45°C; the reported melting temperature of PrP is approximately 65°C, ^77^, and then adding compound SM875.

**Supp. Table 6.**
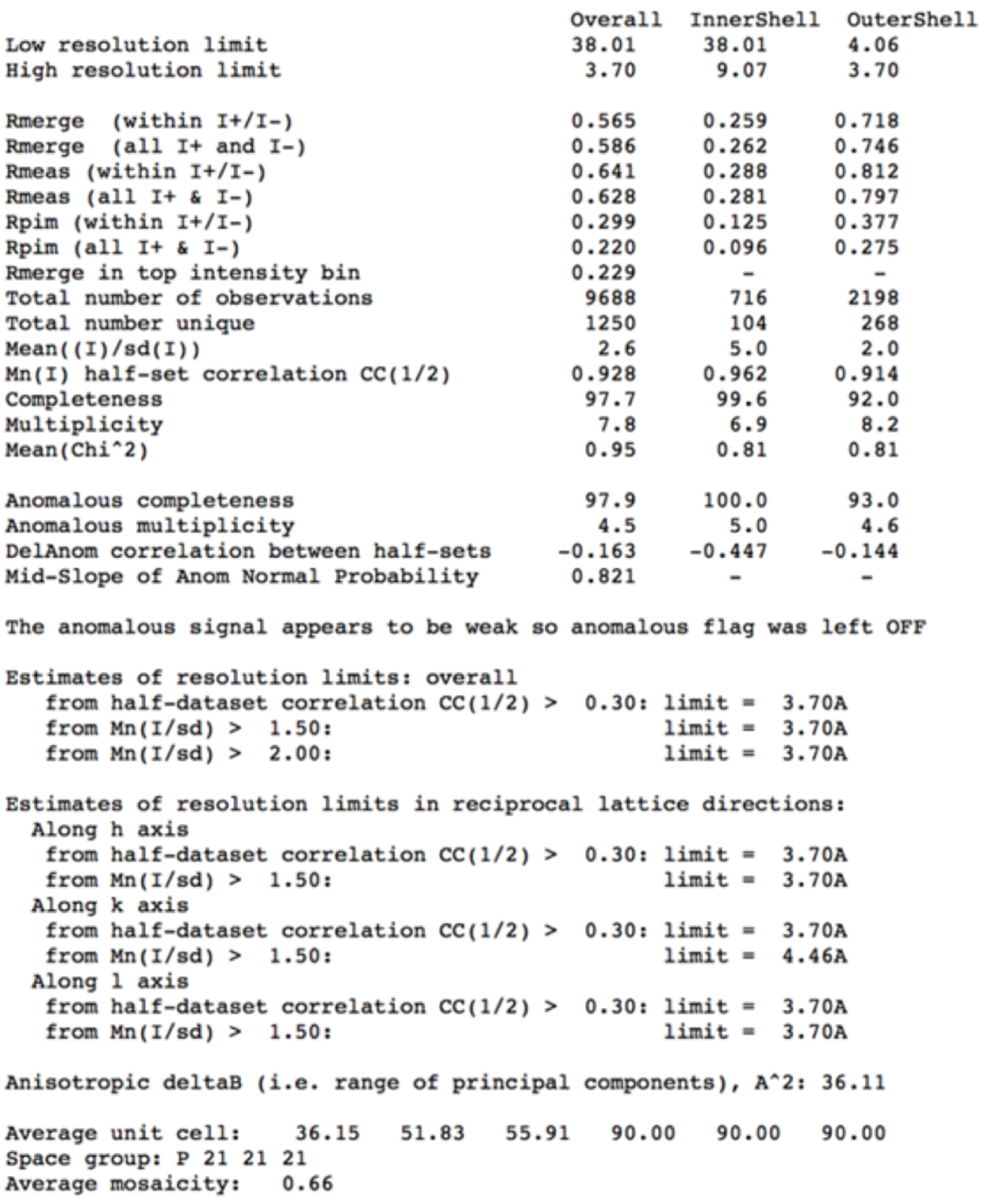
Crystallographic data collection statistics. Diffraction images were indexed and integrated with the XDS software. Reflection intensities were merged, and crystallographic data collection statistics calculated with Aimless (CCP4 suite).

**Supp. Figure 15.**
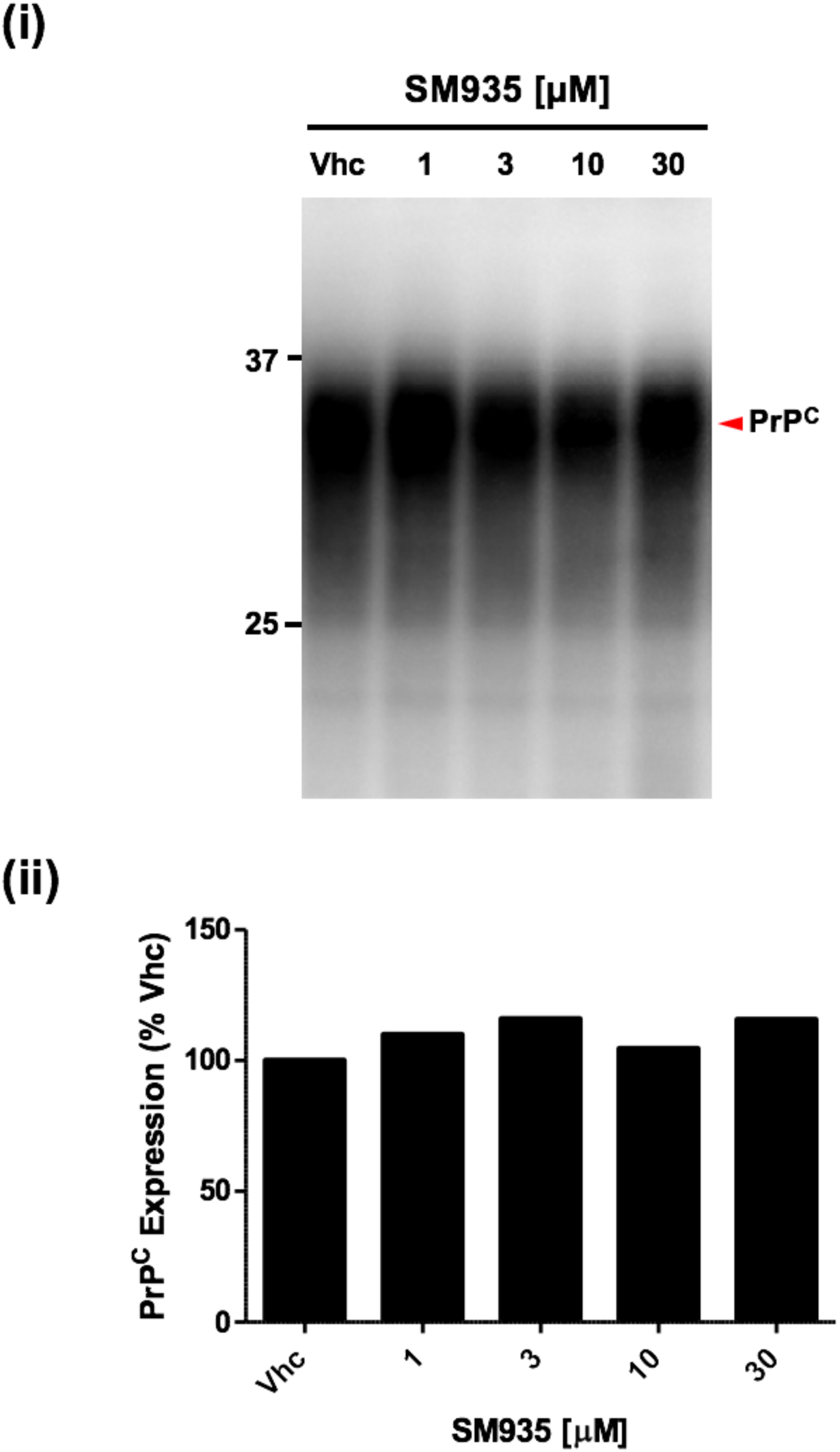
Lack of PrP-lowering effect for compound SM935. HEK293 cells stably transfected with mouse PrP were exposed to different concentrations of SM935 (indicated) or vehicle (Vhc, DMSO, volume equivalent) for 48 h, lysed, and analyzed by western blotting **(i)**. Signals were detected by using a specific anti-PrP (D18). Red arrow indicates the expected sizes of fully glycosylated PrP. The compound shows no effects on the expression of PrP, even at the highest concentration tested (30 µM); **(ii)** the graph shows the densitometric quantification of the levels of full-length PrP from two independent replicates (n = 2). Each signal was normalized on the corresponding total protein lane (detected by UV) and expressed as the percentage of the level in Vhc-treated controls.

